# Claudin-dependent clonal expansion couples epicardial activation to regenerative fate acquisition

**DOI:** 10.64898/2026.10.01.753275

**Authors:** Sofia Kagioglou, Inika Prasad, Tuba Majid, Christopher Y. T. Yen, Sabrina F. Syamsul, Lea M. Vettermann, Delfina Perez, Alberto S. Ceccarelli, Linda R. Micali, Zeyu Yao, Osvaldo Chara, Elif Eroglu

## Abstract

Epicardial activation is a conserved response to injury across vertebrates, yet only in regenerative species do epicardium-derived cells (EPDCs) acquire the capacity to invade the injury site and contribute to cardiac regeneration. How regenerative competence is established in epicardial cells downstream of activation remains poorly understood. Using multicolour lineage tracing we find that epicardial cells of the salamander species *Pleurodeles waltl* undergo a transient phase of clonal expansion that precedes EPDC accumulation at the injury site. Depletion of cCpE-sensitive Claudins from otherwise intact tight junctions alters the organisation and the mechanical state of the epicardium, impairs clonal expansion and reduces the capacity of epicardial cells to delaminate. Consequently, EPDCs fail to activate a regenerative transcriptional programme and complete their transition into injury-induced migratory EPDCs that are required to sculpt a Tenascin-C-rich matrix supporting cardiomyocyte protrusion at the injury site. Together, our findings demonstrate that epicardial activation alone is insufficient for regeneration and identify Claudin-dependent clonal expansion as a mechanism that couples epicardial activation to the acquisition of a regenerative EPDC state.

## Introduction

The epicardium, a layer of mesothelial cells covering the surface of the heart, plays a vital role in heart development and repair. In organisms capable of adult heart regeneration, such as salamanders and teleost fish, a subset of epicardial cells undergoes epithelial-to-mesenchymal transition (EMT), delaminates from the mesothelial layer, and gives rise to epicardium-derived cells (EPDCs) that migrate to the injury site and support myocardial regeneration through paracrine signalling, extracellular matrix (ECM) deposition and contribution to diverse cell types ^1–8^. The postnatal mammalian epicardium is also activated by injury, with epicardial cells re-expressing foetal epicardial genes, proliferating, and acquiring mesenchymal characteristics, yet the EPDCs generated are retained within a thickened epicardial layer on the cardiac surface rather than invading the injury site ^6,10^. Whether this reflects incomplete specification of EPDC identity or a non-permissive environment is unclear, but in either case the constraint lies in the transitions that follow activation. Which transitions these are, and why they proceed in regenerative species but stall in mammals, remains unresolved.

Single-cell studies of the regenerating zebrafish epicardium have identified injury-responsive subpopulations and cell states that are required for regeneration yet constitute only a fraction of the epicardium, including a *hapln1a*-expressing subpopulation that accounts for a minority of *tcf21*⁺cells within the injury site and a transient *ptx3a*⁺*col12a1b*⁺ activated epicardial progenitor cell (aEPC) state ^6,7^. The injury response itself is considerably broader: induction of the aEPC marker *ptx3a*, undetectable prior to injury, extends throughout the ventricular mesothelial epicardium by 1-day post-amputation and becomes confined to the wound by 3 days ^6^. Whether this reflects a restricted subset of cells capable of generating migratory EPDCs, or a capacity that is more broadly distributed across the activated epicardium but adopted by only a minority of cells, cannot be determined from transcriptional profiling or from population-level cell ablation alone and requires resolving the behaviour of individual epicardial cells.

Irrespective of how regenerative competence is distributed among epicardial cells, the transition into EPDCs requires cells to leave the mesothelial layer. Cells that proliferate, rearrange, and ultimately delaminate must continuously remodel their contacts with neighbours, and the capacity to do so may itself determine which activated cells contribute to the generation of migratory EPDCs. Tight junctions are central to such remodelling, regulating paracellular permeability, apical-basal polarity, cytoskeletal mechanics, and cell-cell signalling ^11–14^. Tight junction strands are formed by polymerization of claudins (CLDNs), a family of transmembrane proteins expressed in cell- and tissue-specific combinations that tune junctional function ^15–17^. Critically, tight junctions are remodelled continuously at the molecular level while remaining structurally intact, allowing cells to change shape and exchange neighbours without compromising barrier integrity ^18–21^.

Tight junctions have been studied extensively in organs with sealing epithelia that exert strict permeability control, but remain largely unexplored in the heart, in part because cardiomyocytes lack tight junctions, relying instead on gap junctions, adherens junctions, and desmosomes at the intercalated disc ^22–25^. In contrast to cardiomyocytes, epicardial cells form bona fide tight junctions ^8,26^. We previously showed that the epicardium of the regenerative salamander species *Pleurodeles waltl* (*P. waltl;* Iberian-ribbed newt) expresses *Cldn6*, *Cldn7,* and *Cldn15*, and that perturbation of tight junctions with the C-terminus of *Clostridium perfringens* enterotoxin (cCpE), which targets a subset of CLDNs including CLDN6 and CLDN7, but not CLDN15, impairs cardiac regeneration ^8,27–26^. However, how junctional composition regulates the epicardial injury response has remained unclear.

Here we use a multicolour fluorescent reporter line to assess the clonal dynamics of the regenerating salamander epicardium and show that only a minority of activated epicardial cells amplify, in a transient window that precedes EPDC accumulation at the injury site. Mathematical modelling shows that this clonal expansion cannot be explained by cell proliferation and delamination alone and predicts that clones fragment during regeneration. Upon depletion of cCpE-sensitive CLDNs from epicardial tight junctions, clones fail to expand beyond two cells. At the molecular level, CLDN6 is distributed heterogeneously between the junctions of individual epicardial cells, and its depletion leaves Zonula Occludens-1 (ZO-1)⁺ cell-cell contacts intact while altering the coordinated organisation and mechanical state of the epicardial layer. Spatial and single-nucleus transcriptional profiling identifies an injury-enriched *Tenascin-C*^+^ EPDC state that fails to emerge following CLDN depletion, leaving the mesenchymal compartment dominated by a non-migratory matrix-remodelling EPDC population normally abundant in the uninjured heart. Finally, we show that border-zone cardiomyocytes fail to protrude into an injury site where the Tenascin-C-rich ECM scaffold is present but disorganised.

## Results

### EPDC accumulation at the injury site follows the peak of epicardial proliferation

To determine how restoration of the epicardial layer relates temporally to EPDC accumulation at the injury site, we used the previously established TAT-Cre recombinase mediated lineage tracing strategy, which allows selective labelling of the salamander epicardium in a genetic marker-independent manner ^8^. We utilized salamanders carrying the conditional multi-colour reporter tgTol2(CAG:Nucbow) (hereafter *Nucbow*), in which recombination generates random combinations of nuclear fluorescent proteins ^30,31^ (Fig. 1a). As expected, microinjection of TAT-Cre into the pericardial cavity of the *Nucbow* reporter animals 40 hours prior to the cryoinjury labelled the outermost pan-cytokeratin (pan-CK)-expressing epicardial layer (Fig.1a, b, Extended Data Fig. 1a). At 3 days post cryoinjury (d.p.ci.), the apical tip where the cryoprobe had been applied was devoid of pan-CK^+^ epicardial cells (Fig.1b). A modest increase in the percentage of pHH3^+^ epicardial cells relative to the unoperated controls indicated the onset of injury-induced proliferation (Fig.1c). Extracellular matrix glycoprotein Tenascin-C (TNC) localized to the leading edge of the migrating epicardium and to the border zone between the remaining healthy myocardium and the injury area (Fig.1b). By 5 d.p.ci., TNC^+^ pan-CK^+^ cell layer extended towards the apical tip and TAT-Cre labelled epicardial cells were detectable in this region (Fig.1b). Percentage of pHH3^+^ mitotic epicardial cells increased until 7 d.p.ci., by which time a single layer of pan-CK^+^ epicardial cells covered the injury site (Fig.1b, c). Concomitantly, the first TAT-Cre labelled epicardial progeny (EPDCs) appeared within the injury area, indicating that epicardial cell delamination begins as the epicardial layer is restored (Fig. 1b, d). Consistent with this, expression of the epithelial-to-mesenchymal transition (EMT) gene *Twist-1* was detected in epicardial cells surrounding the border region at 7 d.p.ci. (Extended Data Fig. 1b).

**Fig. 1.**
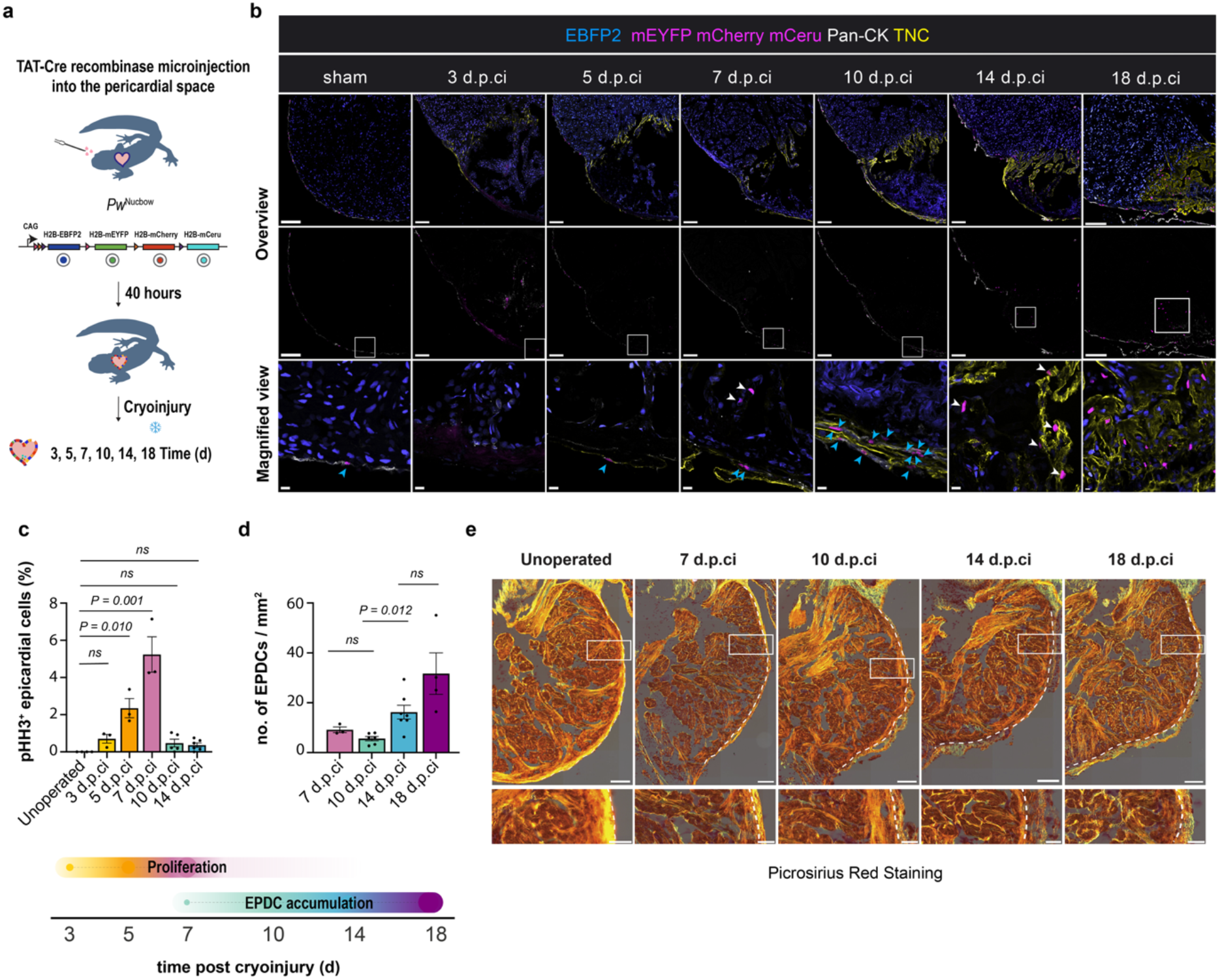
Epicardial proliferation precedes epicardium-derived cell (EPDC) accumulation at the injury site. **a**, Experimental design for permanent labelling of epicardial cells in *Nucbow* reporter *P. waltl*. Schematic of the *Nucbow* reporter construct showing Cre-dependent stochastic expression of H2B-tagged EBFP2, mEYFP, mCherry, and mCerulean (mCeru) fluorescent proteins. **b**, Representative immunostainings for Pan-CK, Tenascin-C (TNC) and mCherry. Lineage traced epicardium and progeny (mCherry^+^, mEYFP^+^, and mCeru^+^) are pseudocoloured in magenta. Scale bars, overview 200 μm, magnified view 20 μm. Blue arrowheads mark TAT-Cre labelled epicardial cells. White arrowheads mark TAT-Cre labelled EPDCs at the injury site. **c**, Epicardial proliferation peaks at 7 d.p.ci. Phospho-histone H3 (pHH3)^+^ Pan-CK^+^ cells were quantified in unoperated (*n* = 4) and cryoinjured hearts at 3, 5, 7, 10 and 14 d.p.ci. *n* = 3 for 3, 5, and 7 d.p.ci, *n* = 5 for 10 d.p.ci, and *n* = 6 for 14 d.p.ci. **d**, Number of TAT-Cre labelled EPDCs at the injury site, quantified at 7, 10, 14, and 18 d.p.ci, and normalized to the injury size. *n* = 3 for 7 d.p.ci, *n* = 6 for 10 d.p.ci, *n* = 7 for 14 d.p.ci, *n* = 4 for 18 d.p.ci. **(c, d)** Each data point represents an animal. Data represent mean ± s.e.m. Kruskal-Wallis test with Dunn’s post-hoc multiple comparisons test. *ns* (not significant)*; P* > 0.05. **e**, Representative picrosirius red staining images of the unoperated and cryoinjured hearts at 7, 10, 14 and 18 d.p.ci. Overview images (top) and magnified views of the regions in the white boxes (bottom). The myocardium–epicardium border is marked by dashed lines. Scale bars, overview 200 μm, magnified view 50 μm.

By 10 d.p.ci., the epicardium at the apical tip had become multilayered and exhibited strong TNC expression, while EPDC density within the injury area remained unchanged relative to 7 d.p.ci. (Fig. 1b, d). The percentage of pHH3⁺ epicardial cells declined between 7 and 10 d.p.ci. and remained low, though still detectable, through 14 d.p.ci. (Fig. 1c). EPDC density increased approximately three-fold between 10 and 14 d.p.ci. (Fig. 1d). Absolute EPDC numbers per section increased in parallel, indicating that the increased EPDC density was not a consequence of changes in the injury size (Extended Data Fig. 1c). Picrosirius Red staining revealed progressive thickening of the epicardial compartment and accumulation of collagen fibres in response to cryoinjury (Fig. 1e).

To determine what underlies the increase in EPDC accumulation at the injury site, we performed lineage tracing in *P. waltl* carrying the conditional reporter *tgTol2(CAG:loxP-GFP-loxP-Cherry)* (hereafter *GFP-loxP-Cherry*), in which recombination induces cytoplasmic Cherry expression ^31^. Analysis of pHH3 staining at 10 d.p.ci. detected no pHH3^+^Cherry^+^ EPDCs within the injury area (262 EPDCs analysed; *n* = 3 animals) (Extended Data Fig. 1d) suggesting that expansion of the EPDCs at the injury site results from continued recruitment of epicardial cells from the outer layer rather than local proliferation at the injury site. Taken together, our results reveal that EPDCs first appear within the injury site as the epicardial layer is restored, but their density increases substantially only after the peak of epicardial proliferation.

### Regenerating epicardium undergoes transient clonal expansion

We next asked how the proliferative response is distributed among individual epicardial cells. To address this question, we performed clonal analysis of epicardial cells in the injury-proximal epicardium, between the atrioventricular canal and the injury border, at 7, 10 and 14 d.p.ci. using *Nucbow* animals (Fig. 2a). The spectral diversity of the *Nucbow* reporter, combined with sparse labelling induced by TAT-Cre recombinase, enabled us to distinguish individual epicardial clones based on their colour identity (Fig. 2b, c) ^8,30^. In sham-operated hearts, pan-CK⁺ epicardial cells exhibited a broad distribution of nuclear colours spanning the visible spectrum, allowing neighbouring cells to be unambiguously distinguished (Fig. 2b, c). To quantitatively define clonal relatedness, we calculated colour spread coefficients in sham-operated hearts, representing pairwise distances between nuclei in three-dimensional RGB colour space (Fig. 2d) ^32^. This distribution established the baseline range of colour variation in the absence of injury and was used to define a colour spread coefficient threshold of 0.05 for clonal identity (Fig. 2d, e, Extended Data Fig. 1e). In both sham-operated and cryoinjured hearts, labelled epicardial cells were considered clonally related if they exhibited a colour spread coefficient below 0.05 and were adjacent among annotated nuclei (Fig. 2e).

**Fig. 2.**
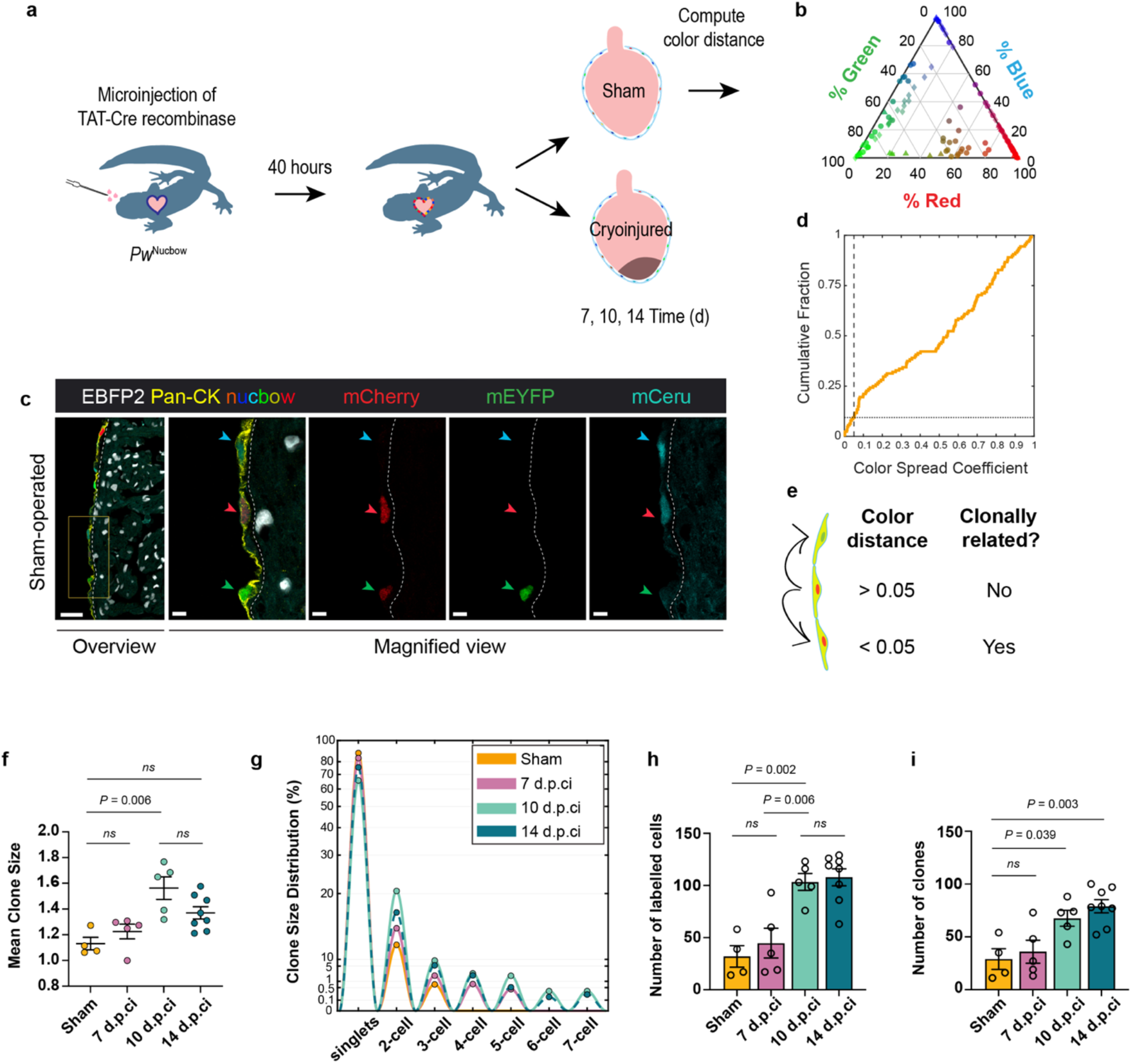
Epicardial cells expand clonally during heart regeneration. **a**, Experimental design for clonality analysis in multicolour fluorescent *Nucbow* reporter *P. waltl*. **b**, Quantification of TAT-Cre-labelled epicardial cell colours in sham-operated hearts plotted as relative channel weights in a ternary plot (red, green, blue channels). *n* = 4 animals depicted as different shapes, 128 cells in total. **c**, Representative images of EBFP2, Pan-CK, mCherry, mEYFP and mCeru. Arrowheads show epicardial cells labelled with distinct colour combinations. Scale bar, overview 50 μm, magnified view 10 μm. The dashed line marks the epicardium-myocardium border. **d**, Cumulative distribution of colour spread coefficients in sham-operated hearts used to assess colour similarity among labelled cells. Dashed lines indicate the threshold (0.05) for clonal assignment. **e,** Schematic illustrating the classification of clonal relationships based on colour distance and spatial proximity; neighbouring labelled cells below the defined threshold (<0.05) are considered clonally related. **f-i,** Quantification of clonal dynamics during regeneration. Mean clone size **(f)**, clone size distribution **(g)**, number of TAT-Cre labelled Pan-CK^+^ epicardial cells analyzed **(h)** and number of clones detected **(i)** in sham-operated and cryoinjured hearts at 7, 10, and 14 d.p.ci. Sham-operated: *n* = 4 animals; 7 and 10 d.p.ci: *n* = 5 animals each, 14 d.p.ci.: *n* = 8 animals. Three sections were analysed per animal. Analysis was restricted to the epicardial region between the atrioventricular canal (avc) and the injury border, corresponding to the injury-proximal epicardium. In **(g)** data points represent the mean percentage of clones within each clone size category per animal; lines connect means across clone size categories and are interpolated for visualization only. In **(f, h, i)** each data point represents one animal. Data are shown as mean ± s.e.m. Statistical significance was determined using Kruskal-Wallis test with Dunn’s multiple comparisons test **(f)** and one-way ANOVA with Tukey’s multiple comparisons test **(h, i)**, *ns; P* > 0.05.

At 3 days after sham operation, labelled epicardial cells were predominantly singlets, with only occasional multi-cell clones detected (Fig. 2f, g). At 7 d.p.ci., mean clone size was not significantly different from sham (Fig. 2f), although clones of up to five cells were detectable (Fig. 2g). By 10 d.p.ci., the number of labelled epicardial cells had increased 3.2-fold compared to sham (Fig. 2h), together with an increase in the total number of clones identified (Fig. 2i). Mean clone size also increased significantly compared to sham, with the clone size distribution shifting rightward and clones of up to seven cells detected (Fig. 2f, g).

By 14 d.p.ci., mean clone size was no longer significantly different from sham (Fig. 2f), while the number of labelled cells and the number of clones remained elevated (Fig. 2h, i). A sustained clone number with declining clone size suggests that the spatial organization of labelled cells changes between 10 and 14 d.p.ci. Taken together, these data demonstrate that epicardial cells undergo a transient phase of clonal expansion during salamander heart regeneration, peaking at 10 d.p.ci. and declining by 14 d.p.ci., as EPDCs accumulate at the injury site.

### Junctional CLDN composition sets the collective organization of the regenerating epicardium

Accommodation of clonally expanding cells within the epicardial layer requires cell rearrangements, changes in packing, or delamination, all of which depend on remodelling of cell-cell contacts ^33^. Having previously found that perturbing tight junctions impairs heart regeneration ^8^, we examined junctional CLDN6 together with the scaffolding protein ZO-1 at 10 d.p.ci., using whole-mount imaging of the apical epicardial surface (Fig. 3a). CLDN6 localized to ZO-1^+^ cell-cell contacts, but its levels varied between individual junctions, including junctions belonging to the same cell (Fig. 3b, c).

**Fig. 3.**
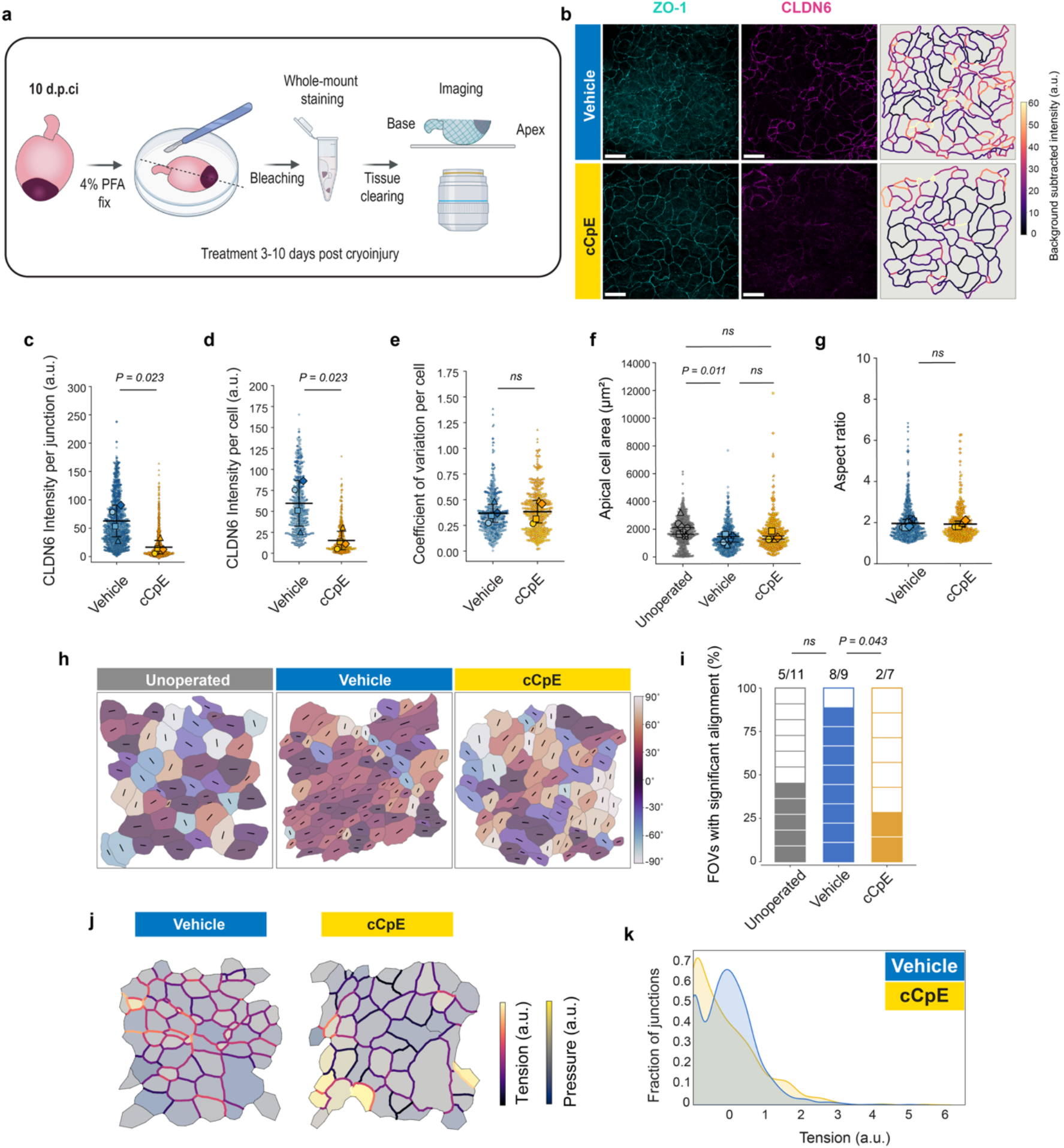
Cell organization and mechanical state of the regenerating epicardium depend on junctional CLDN composition. **a,** Schematic of the whole-mount imaging workflow to visualise epicardial apical surface (top view). **b,** Representative top view images of vehicle- and cCpE-treated epicardium at 10 d.p.ci. From left to right, ZO-1 and CLDN6 immunostaining, quantitative maps of mean junctional CLDN6 intensity (a.u.). Colour scale is shared between conditions; values are background subtracted. Values above 60 are shown as the maximum colour. Scale bars, 50 μm. **c, d,** Quantification of CLDN6 intensity per cell–cell junction (**c**) and per cell (**d**) in vehicle- and cCpE-treated hearts at 10 d.p.ci. Each point represents one junction (**c**) or one cell (**d**). Vehicle, *n* = 1387 junctions from 477 cells; cCpE, *n* = 1115 junctions from 383 cells; 4 animals per group. Data were collected from 6-7 fields of view (FOVs) per condition. Values are background-subtracted. **e,** Coefficient of variation of CLDN6 intensity across the junctions of individual cells. *n* = 476 (vehicle) and *n* = 383 (cCpE) cells from 4 animals per group. **f,** Quantification of apical cell area. *n* = 620 cells (unoperated) from 6 animals, *n* = 672 cells (vehicle) from 6 animals, and *n* = 373 (cCpE) cells from 4 animals. **g,** Aspect ratio of apical cell surface. Vehicle: *n* = 672 cells from 6 animals, cCpE: *n* = 373 cells from 4 animals. **(c-g)** Small symbols depict individual cells; large symbols depict per-animal means **(c-e, g)** or medians **(f)**. Bars are the mean ± S.D. of animal means **(c-e, g),** mean ± IQR of animal medians **(f)**. Statistical significance was determined using Student’s t test **(c-e, g)** and One-way ANOVA with Tukey HSD post-hoc test **(f)**. *ns; P* > 0.05. **h,** Representative orientation maps of the apical epicardial cell surface in unoperated, vehicle-, and cCpE-treated hearts at 10 d.p.ci. Cells are coloured by the orientation of their long axis relative to the image axes; black lines indicate the long axis of each cell. **i,** Proportion of field of views (FOVs) in which cell orientation differed significantly from a uniform distribution (Rayleigh test on doubled angles, *P* < 0.05). Each block represents one FOV; filled blocks indicate FOVs with significant alignment, and the fraction above each bar gives the number of aligned fields. Unoperated, 11 FOVs from 6 animals; vehicle, 6 FOVs from 6 animals; cCpE, 7 FOVs from 4 animals. Groups were compared by restricted permutation test with animal as the permutation block (permutations: 5005 for unoperated versus vehicle, 715 unoperated versus cCpE, 210 vehicle versus cCpE), with Holm correction across three pairwise comparisons; unoperated vs vehicle, *P* = 0.082; unoperated vs cCpE, *P* = 0.400. **j,** ForSys-generated colormaps indicating the relative tensions and pressures in arbitrary units (a.u.) for the vehicle and cCpE conditions (relative to the maximum value of tension or pressure in each FOV). **k,** Gaussian distributions fitted to the histograms of relative tension values for the vehicle- and cCpE-treated hearts. Mann-Whitney test, *P* = 0.006, and Kolmogorov-Smirnov test, *P =* 1.6 x 10^-13^.

To test whether junctional CLDN composition affects epicardial cell behaviour during regeneration, we sought to perturb it. Manipulation of a single CLDN gene often fails to alter permeability, owing to functional redundancy within the CLDN family, necessitating tools that can target several CLDNs simultaneously ^13,34^. One such tool is the non-toxic C-terminus of *Clostridium perfringens* enterotoxin (cCpE), which preferentially targets non-junctional CLDNs and induces their endocytosis, thereby depleting the pool available for incorporation into tight junctions during junctional remodelling ^35^. We previously showed that *P. waltl* CLDN6 is a receptor for CpE ^8^. To confirm that cCpE targets epicardial CLDNs *in vivo*, we generated a fluorescently tagged cCpE fusion protein (mCherry–cCpE). mCherry–cCpE reduced transepithelial electrical resistance (TEER) in Caco-2 monolayers and, when injected intraperitoneally, labelled pan-CK⁺ epicardial cells within one hour both in uninjured and injured hearts, localising to apical tight junctions (ZO-1⁺) and non-junctional lateral membranes (ZO-1⁻), whereas a CLDN-binding-deficient mutant did not (Extended Data Fig. 2a–d).

We treated cryoinjured animals daily with vehicle or cCpE from 3 to 10 d.p.ci. cCpE significantly reduced CLDN6 at cell-cell junctions, while ZO-1 remained continuous (Fig. 3b-d, Extended Data Fig. 2e, f). The relative variability of junctional CLDN6 was unchanged (Fig. 3e). Thus, cCpE altered junctional composition without disrupting cell–cell contacts, consistent with the epicardial expression of *Cldn15*, which is not a CpE receptor and can autonomously form tight junction strands ^8,17,36^.

We next asked how epicardial cells are organized within the regenerating layer. Apical cell area was reduced in the regenerating epicardium at 10 d.p.ci relative to unoperated hearts, reminiscent of the reduced apical area of crowded epithelia, where cells subsequently delaminate (Fig. 3f) ^37^. Epicardial cells in vehicle-treated hearts were frequently in alignment with their neighbours, whereas alignment tended to be less frequent in unoperated hearts (Fig. 3h, i). cCpE treatment did not affect median apical cell area or cell elongation (Fig. 3f, g), yet reduced the proportion of aligned fields, indicating a loss of local order rather than a change in cell shape (Fig. 3h, i, Extended Data Fig. 2g). Consistent with altered epicardial organization, force inference using ForSys showed that cCpE significantly increased the proportion of inferred low-tension junctions (Fig. 3j, k) ^38,36^. Together, these data show that depletion of cCpE-sensitive CLDNs from cell–cell contacts does not affect the size or shape of individual epicardial cells but disrupts their collective organization and the distribution of junctional tension.

### cCpE-sensitive CLDNs are required for clonal expansion and EPDC accumulation

Given the altered organization and junctional tension of the cCpE-treated epicardium, we asked how clonal expansion was affected. We injected TAT-Cre-labelled *Nucbow* animals daily with vehicle or cCpE from 3 d.p.ci. until analysis at 7, 10 and 14 d.p.ci. (Fig. 4a). Because the effect of cCpE is reversible, continued injection maintained loss of CLDN6 from cell–cell contacts through 14 d.p.ci., evidenced by reduced CLDN6–ZO-1 colocalisation (Extended Data Fig. 3a, b) ^8,28,36^. The treatment regimen also recapitulated the impaired regeneration phenotype previously observed following cCpE treatment from 3-10 d.p.ci. (Extended Data Fig. 3c, d) ^8^.

**Fig. 4.**
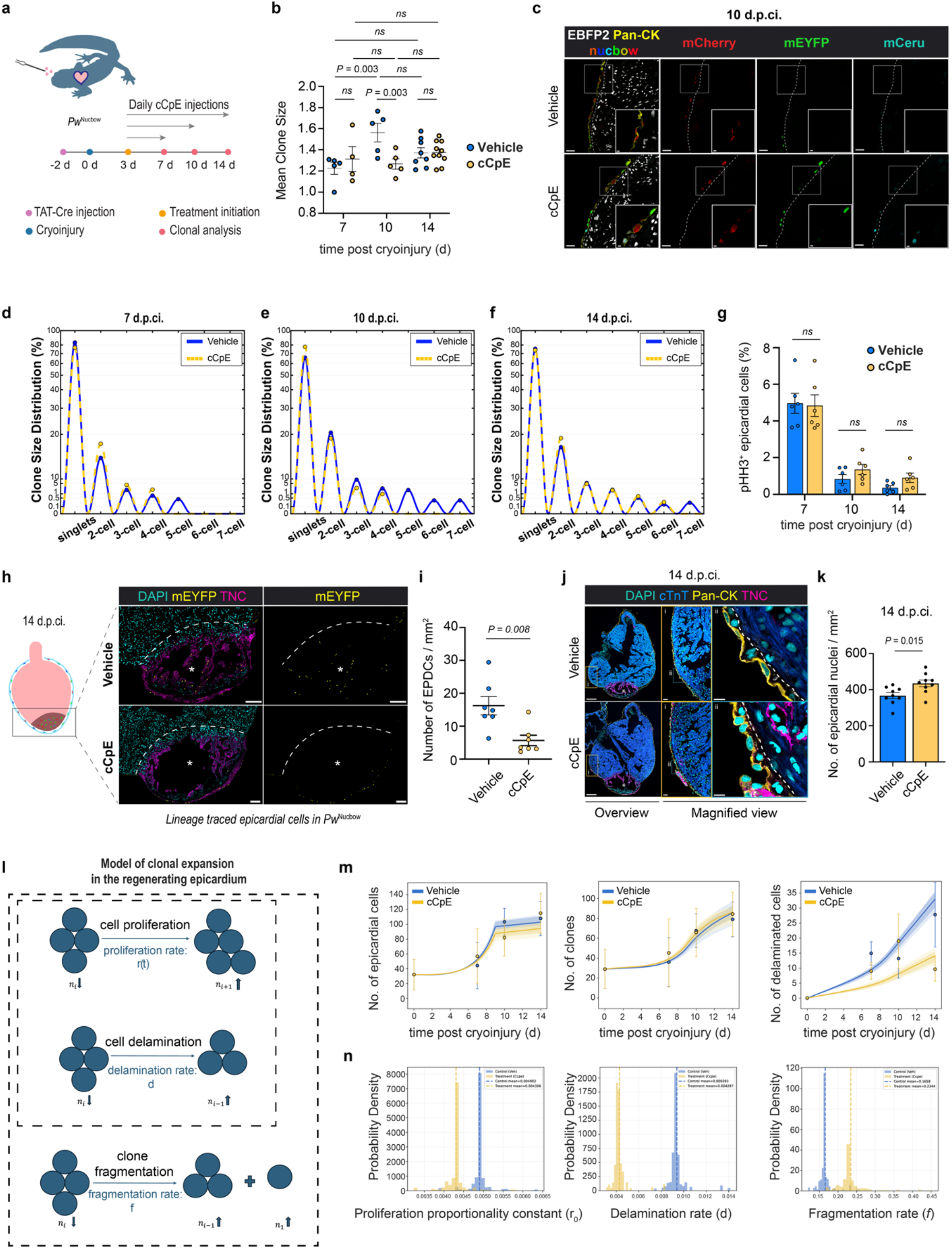
cCpE-sensitive CLDNs are required for clonal expansion and EPDC accumulation. **a**, Experimental design for clonality analysis. **b**, Mean clone size of labelled epicardial clones in vehicle- and cCpE-treated hearts at 7, 10, and 14 d.p.ci. Each data point represents one animal. Data are shown as mean ± s.e.m. Two-way ANOVA with Tukey’s test. **c**, Representative images of EBFP2, Pan-CK, mCherry, mEYFP and mCeru in vehicle- vs. cCpE-treated animals at 10 d.p.ci. Scale bar, overview 50 μm, magnified view 10 μm. **d–f,** Clone size distributions of labelled epicardial clones in vehicle- and cCpE-treated hearts at 7 **(d)**, 10 **(e)**, and 14 d.p.ci. **(f)**. Data points represent the mean percentage of clones within each clone size category per animal; lines connect means across clone size categories and are interpolated for visualization only. **(b, d-f)** Vehicle: *n* = 5 animals at 7 and 10 d.p.ci., *n* = 8 animals at 14 d.p.ci.; cCpE: *n* = 4 animals at 7 d.p.ci., *n* = 5 animals at 10 d.p.ci., *n* = 10 animals at 14 d.p.ci. **g,** Percentage of pHH3^+^ epicardial cells in vehicle- and cCpE-treated hearts at 7, 10, and 14 d.p.ci. Each data point represents one animal. *n* = 6 animals per timepoint per group. Two-way ANOVA with Tukey’s test (**b**, **g**). Data are shown as mean ± s.e.m. *ns; P* > 0.05. **h,** Representative images of lineage-traced mEYFP⁺ EPDCs at the injury site of vehicle- and cCpE-treated *Nucbow* reporter hearts at 14 d.p.ci. Asterisks indicate the injury site. Dashed lines mark the injury border. Scale bar, 200 μm. **i,** Number of lineage traced EPDCs within the injury area in vehicle- and cCpE-treated hearts at 14 d.p.ci normalized to the injury area. Each data point represents one animal. *n* = 7 animals per group. Data are shown as mean ± s.e.m. Welch’s t-test. **j,** Representative immunofluorescence images of vehicle- and cCpE-treated hearts at 14 d.p.ci. stained for DAPI, cardiac Troponin-T (cTnT), Pan-CK and TNC. Left, overview of the ventricle; asterisks mark the injury area. Middle and right, magnified views of the epicardial layer. Dashed line indicates the boundary between the epicardium and the underlying myocardium. Scale bars, 500 μm (overview), 50 μm (i), 10 μm (ii). **k,** Density of epicardial nuclei within the Pan-CK⁺ layer at 14 d.p.ci. Data are shown as mean ± s.e.m.; data points represent individual animals. *n* = 6 animals per group. Welch’s t-test. **l-n,** Mathematical modelling reveals shifts in cell proliferation and clone fragmentation as well as reduced cell delamination following cCpE treatment. **l,** Schematic of the mathematical model of epicardial clonal dynamics, comprising cell proliferation (rate r(t)), cell delamination (rate d) and clone fragmentation (rate f), in which a clone of i cells gives rise to a clone of i−1 cells and a new single-cell clone. Horizontal arrows indicate time; nᵢ denotes the number of clones containing i cells, with vertical arrows indicating whether that number increases or decreases as a result of each process. **m,** The model successfully fits the experimental time courses of the number of labelled epicardial cells (left), number of clones (center), and number of EPDCs at the injury site (right), used as a proxy for the number of delaminated cells for vehicle and cCpE-treated conditions. Solid lines represent the best fits to the experimental data, and shaded regions represent 1 and 2 standard deviations of the parameter distributions obtained from the fitting procedure. Individual points and the error bars correspond to the mean and standard deviation, respectively. (Vehicle/cCpE): for the No. of epicardial cells and No. of clones, 22/23 total, distributed per time point (T0, T7, T10, T14) as 4/4, 5/4, 5/5, 8/10; for the no. of delaminated cells, 17/16 total, distributed as 1/1, 3/4, 6/4, 7/7. **n,** Distributions of the model parameters reveal shifts towards lower values of the proliferation-related parameter *r*₀ (left) and higher values of the fragmentation rate *f* (right) following cCpE treatment. The distribution of the delamination rate *d* (center) is shifted towards lower values following cCpE treatment, and this difference was statistically significant according to the percentile bootstrap method (*P* = 1.06 × 10⁻³). Vehicle and cCpE-treated conditions are shown in blue and yellow, respectively. Vertical dashed lines indicate the mean of each distribution.

Mean clone size was comparable between vehicle- and cCpE-treated hearts at 7 d.p.ci., with overlapping clone size distributions (Fig. 4b, d). By 10 d.p.ci., however, cCpE-treated hearts exhibited a significant reduction in mean clone size compared with the vehicle (Fig. 4b, c). This reflected a selective loss of larger clones: the proportion of clones of three or more cells was reduced (percentage ± s.e.m.; 3.45 ± 1.15 in cCpE vs 13.71 ± 2.66 in vehicle, adjusted *P* = 0.023), with a trend towards a higher proportion of singlets (percentage ± s.e.m.; 77.75 ± 3.67 in cCpE vs 65.73 ± 4.65 in vehicle, adjusted *P* = 0.148), while the proportion of two-cell clones was unchanged (percentage ± s.e.m.; 18.80 ± 2.93 in cCpE vs 20.56 ± 3.06 in vehicle, adjusted *P* = 0.660) (Fig. 4e). Consequently, only 8.6 ± 2.7% of labelled cells resided in clones of three or more cells following cCpE treatment, compared with 30.6 ± 5.0% in vehicle-treated hearts. To exclude the possibility that clones expanded out of the section plane, we identified clonally related cells across five consecutive sections per animal by colour identity alone. Mean clone size remained approximately two-fold lower in cCpE-treated hearts (mean ± s.e.m.; 3.62 ± 0.72 in cCpE vs 7.53 ± 1.10 in vehicle; *P* = 0.039, n = 4 animals per group; Extended Data Fig. 3e), confirming the reduced clonal expansion.

At 14 d.p.ci., mean clone size and clone size distributions were no longer different between the treatment groups (Fig. 4b, f). The percentage of pHH3⁺ pan-CK⁺ epicardial cells did not differ between vehicle- and cCpE-treated hearts at 7, 10 or 14 d.p.ci. (Fig. 4g), nor did the percentage of TUNEL⁺ pan-CK⁺ cells at 10 d.p.ci. (Extended Data Fig. 3f). Together, these data show that depletion of cCpE-sensitive CLDNs restricts clonal expansion beyond two cells, without detectable changes in epicardial mitotic index or survival.

Although clone-size distributions converged by 14 d.p.ci., cCpE-treated hearts had a 3.6-fold reduction in the number of EPDCs at the injury site (Fig. 4h, i). Notably, this reduction was not apparent at 7 or 10 d.p.ci. (Extended Data Fig. 3g). The EPDC deficit therefore followed the divergence in clonal architecture, consistent with the time required for cells to delaminate and migrate to the injury site. Since EPDCs at the injury site do not proliferate at 10 d.p.ci. (see above), local proliferation cannot account for the difference in abundance. The proportion of TUNEL⁺ EPDCs at the injury site showed a trend towards an increase in cCpE-treated hearts that did not reach significance (Extended Data Fig. 3h). Epicardial cell density within the pan-CK⁺ layer was comparable between the treatment groups at 7 d.p.ci., but was significantly higher in cCpE-treated hearts by 10 d.p.ci. and remained elevated at 14 d.p.ci., when the epicardium also appeared buckled (Fig. 4j, k, Extended Data Fig. 3i). Retention of cells within the epicardium and their reduced accumulation at the injury site, without significant changes in proliferation or survival in either compartment, suggests impaired delamination from the epicardial layer.

### Modelling of clonal expansion in the regenerating epicardium

Reduced delamination would be expected to retain cells within clones and thereby increase clone size, yet clone size was reduced, indicating that additional processes shape clonal behaviour in the regenerating epicardium. To quantitatively assess the kinetics of epicardial clonal expansion, we developed a minimal mathematical model in which labelled epicardial cells can proliferate and delaminate (Fig. 4l). In the model, proliferation increases the size of individual clones: when a clone containing a single cell proliferates, it becomes a clone of two cells; when a clone of two cells proliferates, it becomes a clone of three cells; and so on. Thus, a single cell division within a clone of i cells results in a decrease in the number of clones containing i cells and a concomitant increase in the number of clones containing i + 1 cells (Fig. 4l). Delamination was modelled analogously, but with the opposite effect: delamination of one cell from a clone of i cells results in a decrease in the number of clones containing i cells and an increase in the number of clones containing i − 1 cells (Fig. 4l).

A clear prediction of this model is that, when the proliferation rate exceeds the delamination rate, the total number of cells within clones increases over time (Eq. 5; see Methods), consistent with our experimental observations (Fig. 2h). However, this model also predicts that neither proliferation nor delamination of clones larger than one cell changes the number of clones. In contrast, delamination of single-cell clones reduces their number over time (Eq. 6; see Methods), which contradicts the experimentally observed increase in clone number during regeneration (Fig. 2i). Hence, we concluded that transient clonal expansion of the epicardium cannot be explained by cell proliferation and delamination alone and that an additional process must account for the increase in clone number over time.

We reasoned that clones could fragment, with clonally related cells remaining in the section plane but losing contiguity and therefore being scored as independent clones. We therefore extended the model by introducing a fragmentation process in which a clone containing i cells is transformed into a clone containing i − 1 cells and a new clone containing a single cell (Fig. 4l). This process is similar to delamination in that the original clone loses one cell but differs in that the cell remains within the pan-CK^+^ epicardial layer and establishes a new clone rather than leaving the epicardium. We next asked whether the extended model including fragmentation could quantitatively reproduce the experimentally observed dynamics. We first parametrised the proliferation dynamics using the experimental time course of mitotic pHH3^+^ epicardial cells observed during regeneration (Fig. 1c). This left three unknown parameters: a scaling factor linking the number of pHH3^+^ cells to the proliferation rate (*r₀*), and the delamination (*d*) and fragmentation (*f*) rates. To estimate these parameters, we then simultaneously fitted the model to three experimental datasets: the time course of the total number of labelled epicardial cells (*N*), the number of clones (*N_c_*), and the number of EPDCs at the injury site as a proxy for the cumulative number of delaminated cells (*N_d_*). The fitted model successfully recapitulated the experimental dynamics of all three quantities (Fig. 4m), indicating that cell proliferation, delamination, and clone fragmentation are sufficient to explain transient clonal expansion of epicardial cells.

We next asked whether the same model could account for the altered regenerative response observed following the depletion of cCpE-sensitive CLDNs. We independently fitted the model to the three experimental datasets *N*, *N_c,_* and *N_d_* obtained under cCpE treatment. The distribution of the parameter linking the percentage of pHH3^+^ cells to the proliferation rate (*r₀*) was shifted towards lower values under cCpE treatment compared with vehicle-treated hearts (Fig. 4n), consistent with a modest reduction in proliferation that was not resolved by pHH3 staining (Fig. 4g). The fragmentation rate (*f*) showed a tendency towards higher values, whereas the delamination rate (*d*) was significantly reduced (percentile bootstrap method, *P* = 1.06 x 10^-3^). Together, these results indicate that clone fragmentation is required to reproduce the observed clonal dynamics, and that depletion of cCpE-sensitive CLDNs significantly reduces delamination, with tendencies towards reduced proliferation and increased fragmentation.

### Transcriptional and spatial heterogeneity of the regenerating epicardium

We next asked whether the altered epicardial dynamics following cCpE treatment affect the state of the EPDCs generated. We performed single-nucleus RNA sequencing (snRNA-seq) on cardiac nuclei isolated from unoperated, sham-operated, vehicle- and cCpE-treated hearts at 10 d.p.ci., when clonal architecture diverges between the treatment groups (Fig. 5a). Following removal of ambient RNA, low-quality nuclei, and predicted doublets, we integrated the datasets, yielding 67,491 high-quality nuclei that resolved into 27 clusters (Fig. 5b, Extended Data Fig. 4a-c). Using established marker genes, we identified the major cardiac cell populations, including cardiomyocytes, epicardial cells, endothelial cells, fibroblasts, smooth muscle cells, erythrocytes, and immune cells (Extended Data Fig. 4d). All major cell types were detected across experimental conditions, although their relative abundances varied between groups (Extended Data Fig. 5a, b). In parallel, we performed spatial transcriptomics on unoperated, vehicle-, and cCpE-treated hearts at 10 d.p.ci. (Fig. 5a, Extended Data Fig. 5c). Integration of the spatial transcriptomics and snRNA-seq datasets enabled mapping of transcriptionally defined cell populations within the tissue and provided independent validation of cell-type annotations (Extended Data Fig. 5d).

**Fig. 5.**
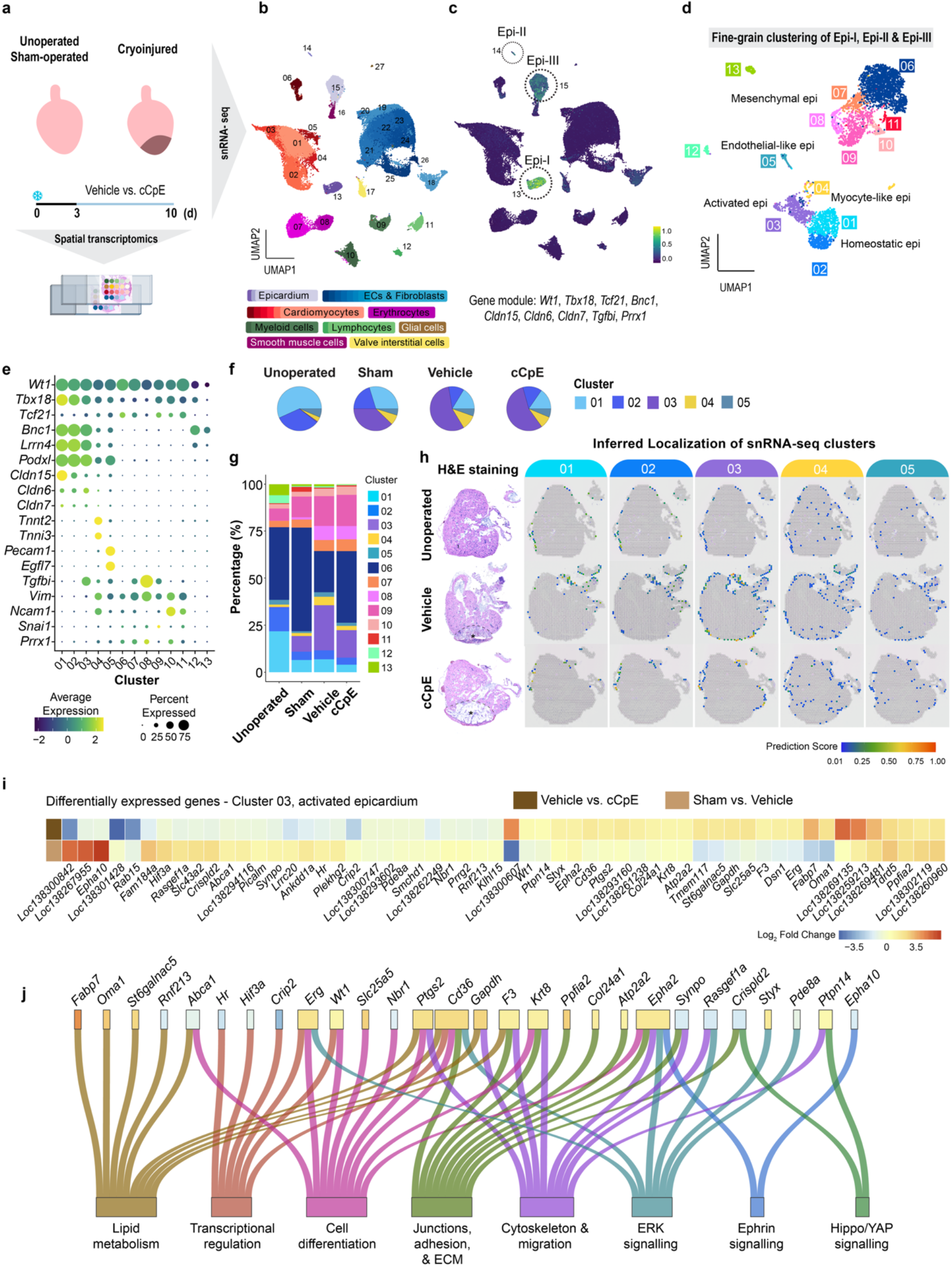
snRNA-seq and spatial transcriptomics reveal transcriptionally and spatially distinct epicardial populations. **a,** Schematic overview of the experimental design. Unoperated, sham-operated, 10 d.p.ci vehicle-treated and cCpE-treated hearts were collected for single-nucleus RNA-sequencing (snRNA-seq) and spatial transcriptomics (10x Visium). snRNA-seq: *n* = 4 pooled unoperated hearts, *n* = 3 pooled sham-operated ventricles, and *n* = 4 pooled cryoinjured ventricles per vehicle- and cCpE-treated group. Spatial transcriptomics: 1 section from *n* = 1 unoperated heart, 8 sections from *n* = 3 vehicle-treated hearts, and 6 sections from *n* = 3 cCpE-treated hearts. **b**, Uniform manifold approximation and projection (UMAP) representation of 67,461 cardiac nuclei following integration of all experimental conditions. Shades indicate cell clusters belonging to major cardiac cell types. **c,** UMAP highlighting epicardial lineage clusters (Epi-I, Epi-II and Epi-III) identified based on expression of an epicardial gene module comprising *Wt1, Tbx18, Tcf21, Bnc1, Cldn15, CldnC (Loc1383043C4), Cldn7, Tgfbi,* and *Prrx1*. **d,** UMAP showing fine grain clustering of Epi-I, Epi-II and Epi-III (4384 nuclei across all conditions), revealing 13 transcriptionally distinct epicardial subclusters corresponding to homeostatic (clusters 01-02), activated (cluster 03), myocyte-like (cluster 04), endothelial-like (cluster 05), mesenchymal (clusters 06-11) and ionocyte-like epicardial states (cluster 12-13). **e,** Dot plot showing the expression of known markers of mesothelial and mesenchymal epicardium, cardiomyocytes, endothelial cells and epithelial-to-mesenchymal transition. **f,** Pie charts showing the relative abundance of clusters in the mesothelial epicardial compartment (01–05) across experimental conditions. **g,** Proportions of epicardial subclusters across experimental conditions. **h,** Spatial localization of epicardial subclusters inferred by integration of snRNA-seq and spatial transcriptomic datasets. Corresponding HGE staining of the sections are shown alongside inferred cluster localizations. Dashed lines demarcate the injury area from the healthy myocardium. Asterisks indicate the injury site. Colour intensity indicates prediction scores > 0.01. **i,** Heatmap showing the log2 fold change of genes differentially expressed in activated epicardium (cluster 03) between vehicle- and cCpE-treated hearts (top row, adjusted *P* < 0.05). Log2 fold changes for the same genes during regeneration (sham-operated versus vehicle-treated hearts) are shown in the bottom row. **j,** Selected differentially expressed genes and associated biological processes, annotated using literature and GO term database ^81–87^.

To identify cells belonging to the epicardial lineage, we scored a gene module of established mesothelial and mesenchymal epicardial markers (Fig. 5c)^6,40–42^. Module scores were highest in three clusters designated Epi-I, Epi-II and Epi-III (Fig. 5c, Extended Data Fig. 5e). Moderate module expression was also detected in cluster 18; however, this cluster was excluded from further analysis owing to its high expression of endothelial markers including *Dkk2*, *Apln*, *Kdr*, *Cdh5* and *Vwf* (Extended Data Fig. 4d). To resolve subpopulations and cell states within the epicardial lineage, cells from Epi-I, Epi-II and Epi-III were subsetted and re-clustered, resulting in 13 transcriptionally distinct epicardial subclusters (Fig. 5d, Extended Data Fig. 6a-c).

Expression of the canonical mesothelial epicardium markers *Bnc1*, *Lrrn4* and *Podxl* identified clusters 01-05 as the mesothelial epicardium (Fig. 5e, f), in which cCpE-responsive CLDNs were expressed (Extended Data Fig. 6d). Clusters 01 and 02 lacked expression of *Tgfbi* and predominated the mesothelial epicardium in unoperated hearts, representing the homeostatic epicardium (Fig. 5e-g). Both mapped to the outer ventricular surface across all conditions (Fig. 5h). Cluster 03 expressed *Stra6*, *Runx1* and *Card10* (Extended Data Fig. 6c) and expanded following both sham operation and cryoinjury, consistent with an activated epicardial state (Fig. 5f, g, Extended Data Fig. 6e). Cluster 04 retained canonical epicardial markers while co-expressing cardiomyocyte-associated genes (*Tnnt2*, *Tnni3*, *Actn2*, *Trdn*, *Myh7b,* and *Mybpc3*) (Fig. 5e, Extended Data Fig. 6c). Cluster 05 similarly expressed epicardial markers but additionally expressed vascular and lymphatic endothelial-associated genes (*Egfl7*, *Notch4*, *Lyve1,* and *Mmrn2*), suggestive of an endothelial-like epicardial state (Fig. 5e, Extended Data Fig. 6c).

The remaining subclusters fell into two groups. Clusters 06–11 formed a distinct mesenchymal branch characterized by downregulation of *Bnc1*, *Lrrn4*, *Podxl*, *Cldn6*, *Cldn7,* and *Cldn15* and upregulation of *Vim*, *Ncam1*, *Snai1,* and *Prrx1*, consistent with EPDCs (Fig. 5e). In contrast, clusters 12 and 13 expressed ion transport genes (*Ca12*, *Slc26a4*, *Slc4a1*, and *Bsnd*), suggestive of an ionocyte-like identity with potential roles in pericardial fluid homeostasis (Extended Data Fig. 6c). Clusters 12 and 13 were most abundant in unoperated hearts and remained largely unaffected by cCpE treatment (Fig. 5g).

### The activated epicardial state is maintained but its transcriptional programme is altered

As the activated epicardium (cluster 03) was the most abundant mesothelial population in regenerating hearts and expressed cCpE-sensitive *Cldn* genes, we asked how this population was affected by cCpE treatment. The proportion of cluster 03 was unchanged between the treatment groups, and RNAscope *in situ* hybridization showed that *Stra6* was expressed in pan-CK⁺ epicardial cells of both vehicle- and cCpE-treated hearts, indicating that the activated state was maintained under cCpE treatment (Fig. 5f; Extended Data Fig. 6e). To determine whether cCpE treatment nonetheless induced transcriptional changes within the activated epicardium, we performed differential gene expression (DGE) analysis and identified 56 differentially expressed genes that were associated with junction and ECM organisation, cytoskeleton and migration, lipid metabolism, and pathways including Ephrin, Hippo/YAP and ERK1/2 signalling (Fig. 5i, j).

A subset of genes was upregulated during regeneration but attenuated upon cCpE treatment, including *Hif3a*, *Rasgef1a*, *Crispld2*, *Synpo*, *Hr*, *Plekhg2* and *Crip2* (Fig. 5i). Several have established roles in cytoskeletal remodelling, including *Plekhg2*, a Rho guanine nucleotide exchange factor driving Rac1- and Cdc42-dependent actin remodelling, and *Synpo*, an actin-associated protein (Fig. 5j) ^43,44^. *Hr*, which modulates H3K9 methylation and maintains heterochromatin, was also attenuated, suggesting that junctional CLDN composition may additionally affect epigenetic components of the regenerative response ^45,46^.

Conversely, cCpE treatment increased expression of several genes that were normally downregulated during regeneration, including *St6galnac5*, *Slc25a5*, *Erg*, *Oma1* and *Fabp7* (Fig. 5i). Among these, induction of *St6galnac5*, an α2,6-sialyltransferase that modifies cell-surface ganglioside composition, is notable given that increased sialylation has been associated with loss of regenerative competence in the neonatal mouse heart ^47^ ^48^. Together, these data show that the activated epicardial state is maintained, yet its transcriptional programme is altered by cCpE treatment, identifying candidate effectors downstream of junctional CLDNs.

### cCpE treatment alters EPDC composition and softens the epicardium

Given these changes in the activated epicardium, we next asked whether the EPDC populations were affected. Cluster 06, hereafter resident EPDCs owing to their predominance in uninjured hearts, expressed transcripts shared across the EPDC compartment, including *Dab1*, *Lix1*, *Phex*, *Col5a3* and *Vwce*, and comprised 72% of EPDCs in sham-operated hearts (Fig. 6a, b). Following cryoinjury, their proportion declined to 39% in vehicle-treated hearts, but less so in cCpE-treated hearts (52%) (Fig. 6b). Spatial mapping localized the transcriptional signature of resident EPDCs to the outer layer of the regenerating heart (Fig. 6c). Cluster 08 showed the converse pattern: it comprised 2% of EPDCs in sham-operated hearts, increased to 14% following cryoinjury in vehicle-treated hearts (Fig. 6b), and localized preferentially to the injury site (Fig. 6c).

**Fig. 6.**
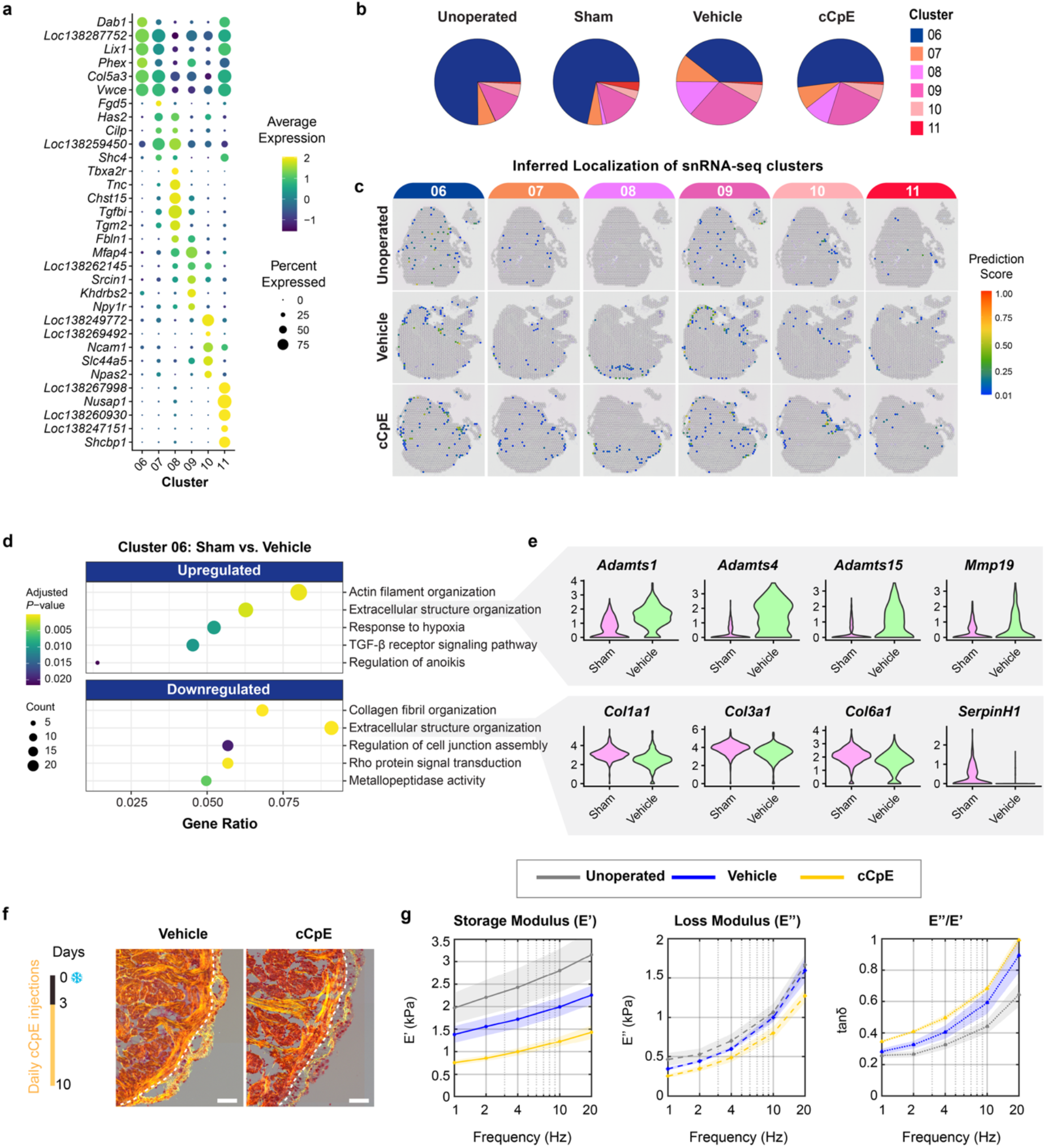
Mesenchymal EPDC populations undergo injury-induced remodelling. **a,** Dot plot showing expression of representative marker genes across EPDC clusters (06–11). **b,** Pie charts showing the relative abundance of clusters 06–11 across experimental conditions. **c,** Spatial localization of mesenchymal EPDC clusters inferred by integration of snRNA-seq and spatial transcriptomic datasets. Colour intensity indicates prediction score > 0.01 for each cluster. **d,** Dotplot showing selected Gene Ontology (GO) terms overrepresented in genes differentially expressed in Cluster 06 between sham-operated and vehicle-treated hearts **e,** Violin plots showing expression of selected extracellular matrix-associated genes in Cluster 06. **f,** Representative images of heart sections stained with Picrosirius Red in vehicle- and cCpE-treated animals at 10 d.p.ci. Dashed line marks the epicardium-myocardium border. Scale bars, 100 μm. **g,** Dynamic mechanical analysis (DMA) of the epicardium. Frequency dependence of the storage modulus (E′), loss modulus (E′′), and loss tangent (tan δ = E′′/E′) in unoperated, vehicle- and cCpE-treated hearts at 10 d.p.ci. Shaded regions indicate s.e.m. Statistical significance was assessed by pairwise comparison of DMA curves using linear hypothesis tests (MATLAB *coefTest*) on the fixed-effects coefficients of a linear mixed-effects model, with animal and measurement (nested within animal) as random-effects and condition and frequency as fixed-effects (Satterthwaite F-test, Bonferroni-corrected). No significant differences were detected between unoperated and vehicle-treated hearts for E′, E′′ or tan δ. Compared with vehicle-treated hearts, cCpE-treated hearts exhibited reduced E′ (*P* = 0.006) and increased tan δ (*P* = 0.016), whereas E′′ was unchanged (*P* = 0.367). Tan δ was also increased in cCpE-treated relative to unoperated hearts (*P* = 0.047).

To understand what the greater representation of resident EPDCs under cCpE treatment implies, we first characterised their response to injury by comparing sham-operated and vehicle-treated hearts. DGE analysis identified 624 genes differentially expressed in response to injury. Gene ontology (GO) analysis identified extracellular structure organisation among both up- and downregulated genes: matrix turnover genes were induced, whereas collagens and the collagen chaperone *Serpinh1* were reduced (Fig. 6d, e).

This transcriptional shift towards matrix remodelling, together with the localization of these cells to the outer layer of the heart (Fig. 6c) and their greater representation under cCpE treatment (Fig. 6b), led us to hypothesize that the mechanical properties of the epicardium and underlying matrix might be altered upon cCpE treatment. Picrosirius Red staining revealed a less compact subepicardial collagen network in cCpE-treated hearts compared with vehicle controls at 10 d.p.ci. (Fig. 6f). To determine whether these alterations were accompanied by changes in tissue mechanics, we performed dynamic mechanical analysis using nanoindentation (Extended Data Fig. 7a, b). The epicardial layer exhibited viscoelastic behaviour, with the storage modulus (E′) exceeding the loss modulus (E′′) across all frequencies tested (Extended Data Fig. 7c). Following cryoinjury, storage modulus declined, reaching significance by 14 d.p.ci., while the loss tangent increased, indicating a shift towards a softer and more fluid-like mechanical state, reminiscent of tissue fluidisation (Extended Data Fig. 7c-e) ^46^. Epicardium in cCpE-treated hearts exhibited a reduced storage modulus and an elevated loss tangent at 10 d.p.ci., indicating a softer and more fluid-like state than vehicle-treated hearts at this time-point (Fig. 6g). The stiffness of the underlying myocardium did not differ between treatment groups, excluding broad changes in cardiac mechanics (Extended Data Fig. 7f).

Taken together, cCpE treatment shifted the EPDC composition towards resident EPDCs, accompanied by untimely softening of the epicardium.

### Newly formed EPDCs fail to acquire the regenerative state

As delamination and the transitions that generate EPDCs occur within this altered mechanical environment, we next asked which states newly formed EPDCs acquire in vehicle- and cCpE-treated hearts. Clusters 07–10 expanded following cryoinjury in vehicle-treated hearts, identifying them as injury-enriched EPDC states. Their expansion was less pronounced in cCpE-treated hearts (Fig. 6b). Cluster 08, hereafter the regenerative EPDC state, was the most injury-enriched population and the one localizing to the injury site (Fig. 6c). To identify these populations *in vivo*, we performed RNAscope *in situ* hybridization at 10 d.p.ci., restricting our analysis to the outer epicardial layer and the subepicardium, where recently delaminated cells are found ^50^. We adopted a combinatorial marker approach to distinguish transcriptionally similar EPDCs. *Bnc1* marked the mesothelial epicardium (clusters 01-05) and its expression was confined to the pan-CK^+^ epicardial layer, while *Phex* broadly marked all EPDCs (clusters 06-11). Cells within the subepicardial compartment were assigned identity by combinations of *Phex*, *Cilp*, *Tnc* and *Fbln1* (Fig. 7a). In vehicle-treated hearts, cells within the subepicardium showed low *Cilp* expression and comprised both *Tnc^high^Fbln1^high^* (cluster 08) and *Tnc^low^Fbln1^high^* (cluster 06) cells (Fig. 7b, c). In cCpE-treated hearts, the subepicardial compartment was instead dominated by *Cilp*^high^ and *Tnc^low^* cells, corresponding to cluster 07, while *Tnc^low^ Fbln1*^high^ (cluster 06) cells appeared less frequent (Fig. 7b, c). The absence of *Tnc^high^Fbln1^high^* cells in the subepicardial compartment suggests a failure of newly formed EPDCs to reach the injury-enriched cluster 08 state.

**Fig. 7.**
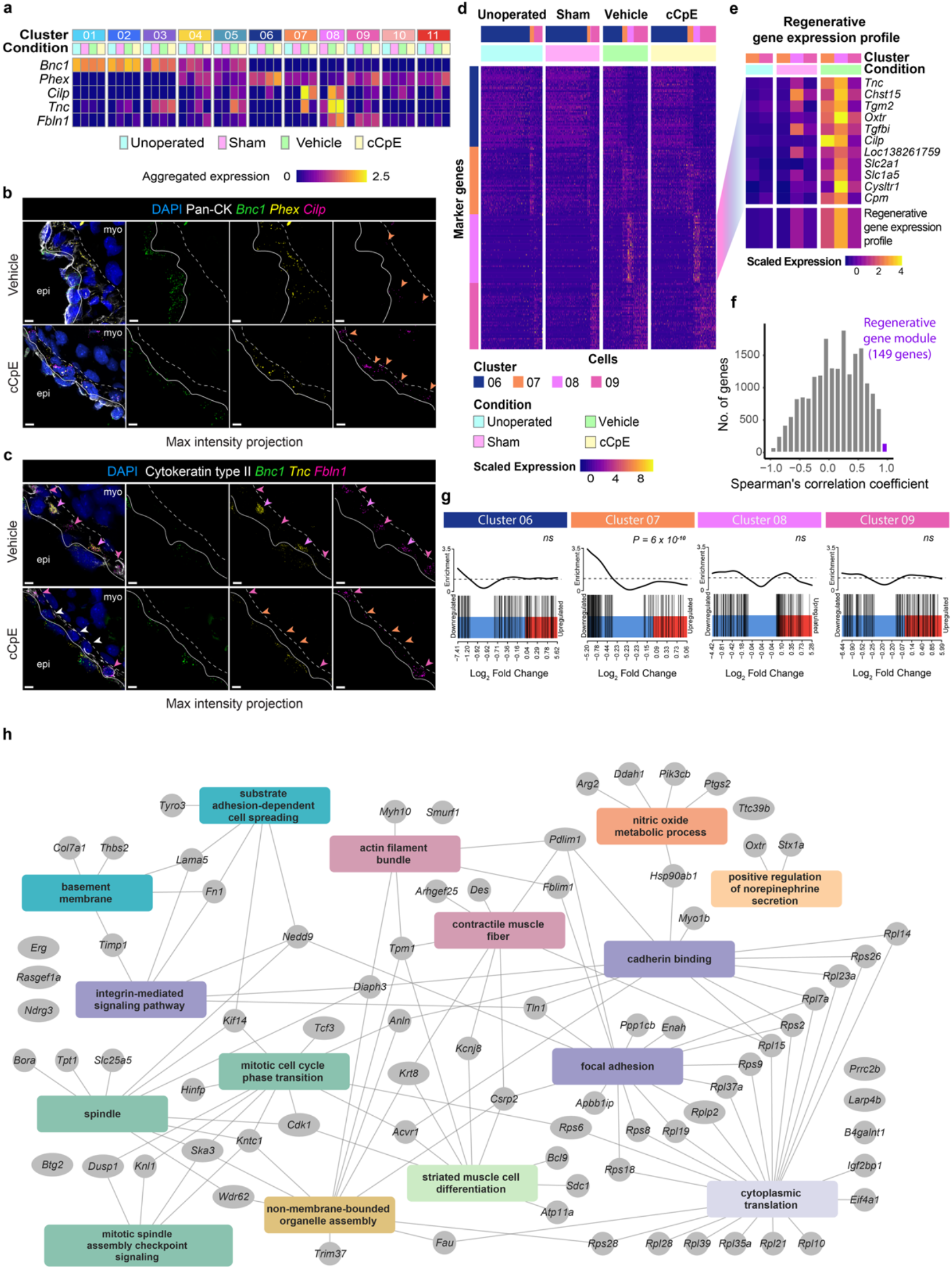
Depletion of cCpE-sensitive CLDNs attenuates emergence of regenerative EPDC state. **a**, Heatmap showing aggregated expression of genes selected for RNAscope *in situ* hybridization across clusters and conditions **b**, **c,** Representative images of RNAscope *in situ* hybridization against *Bnc1*, *Phex* and *Cilp* **(b)** and *Bnc1*, *Tnc* and *Fbln1* **(c)** in vehicle- and cCpE-treated hearts at 10 d.p.ci. Pan-CK and Cytokeratin type II delineate the epicardial and subepicardial regions. Solid lines indicate the boundary between the epicardial and subepicardial compartments, whereas dashed lines indicate the boundary between the sub-epicardium and myocardium. Arrowheads indicate putative subepicardial EPDCs. Maximum-intensity projections are shown. Scale bars, 10 μm. **d**, Heatmap showing scaled expression of the top marker genes ranked by log2 fold changes across mesenchymal EPDC clusters 06–06. The top markers for Cluster 08 are highlighted, a subset of which are expressed in Cluster 07 and 06 in cryoinjured conditions and shown in **e**, along with their aggregated expression across conditions excluding the cCpE-treated condition. Their expressions were averaged to define the putative regenerative gene expression profile. **f**, Distribution of Spearman’s correlation coefficients between all genes and the putative regenerative gene expression profile are shown in **e**. Genes with invalid correlation coefficients (4053 genes, see Methods) were excluded. Genes with correlation coefficients >0.6 were defined as the putative regenerative gene module (146 genes). **g,** Barcode plot showing the log2 fold-change based ranks of the putative regenerative gene module genes in vehicle- vs. cCpE-treated conditions for clusters 06-06. Indicated *P-*values were calculated using a competitive gene set enrichment test. **h**, Network representation of selected GO terms overrepresented (adjusted *P*-value < 0.05) in the regenerative gene module. Rectangular nodes show overrepresented GO terms; edges connect genes to the GO terms they belong to. Oval-shaped nodes represent ERK1/2 pathway components.

To assess this independently of RNAscope markers, we asked which genes define the cluster 08 state. Inspection of the top 50 markers for clusters 01-11 and their condition-specific expression patterns revealed that a subset of cluster 08 markers was highly expressed in clusters 07 and 09 following cryoinjury, but not in resident EPDCs (Fig. 7d, Extended Data Fig. 8a). We averaged the aggregated expression of cluster 08 markers following this pattern, using only unoperated, sham-operated and vehicle-treated conditions, to define a regenerative gene expression profile (Fig. 7e). Spearman correlation of all genes with this profile identified 146 genes (ρ > 0.6), hereafter called the putative regenerative gene module (Fig. 7f; Extended Data Fig. 8b). To determine whether activation of this module was affected by cCpE treatment, we tested for its enrichment among the log_2_ fold changes of all genes between vehicle- and cCpE-treated conditions in clusters 06-06. Significant enrichment was detected only in cluster 07, where module genes were preferentially downregulated following cCpE treatment, with no enrichment in clusters 06, 08, or 09 (Fig. 7g). Thus, in cluster 07 cells, most abundant in the cCpE-treated subepicardium, activation of the gene expression programme they share with cluster 08 was attenuated. GO term analysis revealed that the module was enriched for terms related to cell adhesion and motility, including integrin-mediated signalling, substrate adhesion-dependent cell spreading, and focal adhesion and included regulators of epicardial invasion and EPDC maturation such as *Smurf1* and *Myh10* (Fig. 7h) ^51,52^. The module therefore comprises genes associated with the capacity of EPDCs to traverse the subepicardium and reach the injury site.

Beyond cell invasion and migration genes, we also observed expression of ERK1/2 signalling pathway components within the module (Fig. 7h). We therefore tested whether ERK signalling was downstream of cCpE-sensitive CLDNs. Nuclear phospho-ERK1/2 (p-ERK1/2) levels in epicardial cells were comparable between vehicle- and cCpE-treated hearts (Extended Data Fig. 6a), indicating that cCpE-treatment does not alter nuclear ERK activity. However, inhibiting ERK phosphorylation using MEK inhibitor PD0325901 (PD) reduced the proportion of pHH3⁺ epicardial cells (Extended Data Fig. 6b), providing us with an alternative approach to block clonal expansion. Daily PD administration from 8 to 10 d.p.ci. reduced p-ERK1/2 levels and significantly lowered the proportion of clones containing three or more cells without affecting the proportion of two-cell clones, mirroring the clonal phenotype observed with the cCpE treatment (Extended Data Fig. 6c–f). Furthermore, *Tnc^high^Fbln1^high^* cells were absent from the subepicardial compartment of PD-treated hearts (Extended Data Fig. 6g, h). Thus, a second perturbation that restricts clonal amplification likewise prevents EPDCs from reaching the cluster 08 state.

### Cardiomyocyte protrusion into the TNC-rich ECM scaffold is impaired

Heart regeneration ultimately requires lost cardiomyocytes to be replaced ^53,54^. As border-zone cardiomyocytes dedifferentiate and give rise to most of the new muscle, we asked whether changes in the EPDCs affected them. Cardiomyocyte proliferation is unaffected by cCpE treatment at 10 d.p.ci. ^8^, so we reasoned that processes independent of proliferation might underlie the impaired regeneration. Unsupervised clustering of spatial transcriptomics data from unoperated, vehicle-, and cCpE-treated hearts at 10 d.p.ci. identified 22 clusters among which cluster 10 localized to the injury border and expressed the established border zone marker *Ankrd1* (Fig. 8a, b, Extended Data Fig. 10a-d) ^55^. Although spatial cluster 10 was detected at low frequency in unoperated hearts, it lacked the spatial organization observed following cryoinjury, consistent with its emergence as a distinct border-zone population during regeneration (Fig. 8a, b). DGE analysis of the border zone cluster between vehicle- and cCpE-treated hearts identified 100 differentially expressed genes, enriched for muscle development and differentiation, sarcomere organization, cell adhesion, ECM and collagen organisation, and cytoskeleton and actin organization (Fig. 8c; Extended Data Fig. 10e). We further noted coordinated upregulation of Z-disc and costamere components, including *Flnc*, X*irp1*, *Myot*, *Pdlim3* and *Des*, together with the basement membrane collagens *Col4a1* and *Col4a2* (Fig. 8c) ^56–61^.

**Fig. 8.**
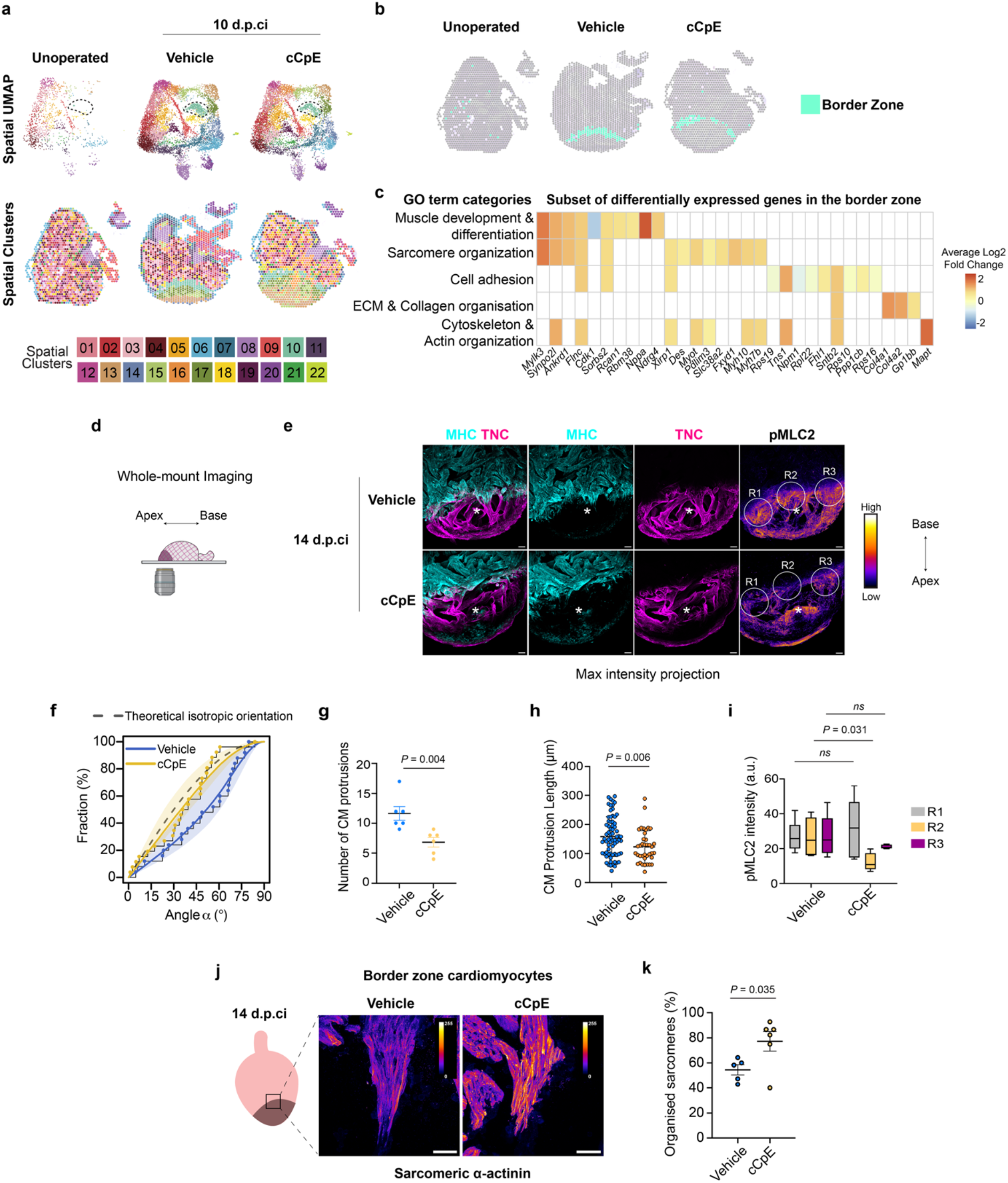
Cardiomyocytes fail to protrude into a disorganized ECM scaffold. **a,** Spatial transcriptomic clusters from unoperated, and vehicle- and cCpE-treated hearts at 10 d.p.ci shown as UMAP projections and spatially mapped tissue sections. **b,** Spots assigned to Cluster 10 (border zone) are highlighted. **c**, Heatmap showing selected categories summarizing significantly overrepresented GO terms for genes differentially expressed between vehicle- and cCpE-treated hearts in the border zone (Cluster 10) (adjusted *P* < 0.05, q < 0.05). Genes comprising these GO terms are coloured by their log2 fold changes. **d,** Schematic illustrating the whole-mount imaging approach used to visualize the injury site. **e,** Representative whole-mount immunostainings of the injury site for myosin heavy chain (MHC), TNC and phosphorylated myosin light chain 2 (pMLC2) in vehicle- and cCpE-treated hearts at 14 d.p.ci. Maximum-intensity projections are shown. pMLC2 signal is displayed using a Fire lookup table (LUT). R1–R3 denote regions of interest used for quantification of pMLC2 fluorescence intensity in **h**. Scale bars, 100 μm. **f,** Cumulative distribution plot of TNC^+^ bridge orientation at 14 d.p.ci. Each point represents an individual TNC^+^ scaffold. The grey dashed line represents the expected distribution for random orientation within a 3D sphere, while the solid-coloured lines represent fitted to the empirical cumulative distribution functions (ECDFs) for vehicle and cCpE. 65% confidence intervals of each treatment are shaded in the respective treatment colours. Vehicle, *n* = 25 scaffolds from 7 animals; cCpE, *n* = 26 scaffolds from 8 animals. Kolmogorov–Smirnov (KS) test: vehicle vs isotropic distribution, P = 0.030; cCpE vs isotropic distribution, P = 0.136. **g,** Number of cardiomyocyte (CM) protrusions at the injury site at 14 d.p.ci. Each point represents one individual animal (*n* = 6 per group), two-tailed Mann-Whitney test. **h,** Quantification of CM protrusion length at 14 d.p.ci. Each point represents an individual protrusion. Vehicle, *n* = 70 protrusions from 6 animals; cCpE, *n* = 38 protrusions from 6 animals. Median values are indicated, two-tailed Mann–Whitney test. **i,** Quantification of pMLC2 fluorescence intensity within regions of interest (R1–R3) shown in **e**. Statistical significance was assessed using a mixed-effects model (REML) followed by Tukey’s multiple-comparisons test. Vehicle, *n* = 6 animals; cCpE, *n* = 5 animals, with three measurements (R1–R3) obtained per heart. **j,** Representative images of border-zone CMs stained for sarcomeric α-actinin in vehicle- and cCpE-treated hearts at 14 d.p.ci. A schematic indicating the location of the imaged region within the injury border zone is shown on the left. Fluorescence intensity is displayed using a Fire LUT to visualise sarcomere organization. Scale bars, 50 μm. **k,** Percentage of border-zone muscle protrusions containing organised sarcomeres within 30 μm of the protrusion tip, in vehicle- and cCpE-treated hearts at 14 d.p.ci., assessed by sarcomeric α-actinin staining. Points represent individual animals; bars indicate mean ± s.e.m. Vehicle, *n* = 5 animals; cCpE, *n* = 6 animals. Welch’s t-test.

Border-zone cardiomyocytes extend actomyosin-rich protrusions into the injury site ^62,63^. Given the transcriptional changes suggesting altered sarcomere and cytoskeletal organisation, we asked whether these protrusions were affected at 14 d.p.ci., when EPDC accumulation at the injury site is reduced. To gain optical access to the injury site for whole-mount imaging, we bisected vehicle- and cCpE-treated hearts at 14 d.p.ci. along the median plane and performed immunofluorescence staining against TNC, myosin heavy chain (MHC), and phosphorylated myosin light chain 2 (pMLC2) (Fig. 8d, e). In vehicle-treated hearts, the TNC-rich ECM scaffold was organized into bridge-like structures reminiscent of trabeculae (Fig. 8e). MHC⁺ cardiomyocytes extended protrusions into these TNC-rich bridges, which spanned radially from the border zone towards the injury tip (Fig. 8e). Bridge orientation in vehicle-treated hearts deviated significantly from a theoretical isotropic distribution, whereas in cCpE-treated hearts it was not distinguishable from an isotropic distribution (Fig. 8f). TNC levels were unchanged between groups (Extended Data Fig. 10f). cCpE treatment reduced both the number and length of cardiomyocyte protrusions (Fig. 8g, h) and altered regional pMLC2 distribution, suggesting localized changes in actomyosin contractility (Fig. 8e, i). Border-zone cardiomyocytes in cCpE-treated hearts exhibited increased sarcomeric organization as assessed by α-actinin staining, consistent with retention of a mature contractile state (Fig. 8j, k). Together, these data show that following cCpE treatment the TNC-rich ECM scaffold is no longer detectably organised and border-zone cardiomyocytes extend fewer and shorter protrusions into the injury site, explaining in part the regenerative defect.

## Discussion

Our work demonstrates that regenerative competence of the salamander epicardium depends on the CLDN composition of its tight junctions and identifies claudin-dependent clonal expansion as a mechanism that couples injury-induced epicardial activation to the emergence of migratory EPDCs that contribute to muscle regeneration. Despite widespread epicardial activation in response to cryoinjury, only a minority of clones expand beyond the two-cell stage, indicating that heterogeneity within the regenerating epicardium extends beyond transcriptional state to include differences in clonal amplification. Our data show that cCpE-sensitive CLDNs are dispensable for the first division of an epicardial cell but required for the subsequent ones that give rise to larger clones. Inhibition of ERK1/2 phosphorylation between 8–10 d.p.ci. phenocopies this selective loss, placing the amplifying divisions after closure of the epicardial layer at 7 d.p.ci. Thus, widespread proliferation in the first week after cryoinjury restores the epicardial layer itself, whereas the subsequent CLDN- and ERK-dependent amplifying divisions are required for the generation of migratory EPDCs.

In the developing mouse heart, mitotic spindle orientation determines whether proliferating epicardial cells remain in the layer or invade the underlying myocardium ^64^. While spindle orientation might also be important in the regenerating epicardium, our data favour a model in which delamination is determined by the local context created by repeated divisions within a clone, rather than at each cell division. Reduced clone size decreased the rate of delamination even though divisions still occurred. Work in other epithelia suggests mechanical forces might generate such local context ^37,65–67^. In the *Drosophila* notum, proliferation leads to crowding, which then induces delamination to balance growth ^37^. In the mammalian epidermis, crowding is sufficient to trigger differentiation in the basal epidermal layer. Interestingly, cells that initiate differentiation and subsequently delaminate are often neighbours of dividing cells, as junctions at the interface of a mitotic cell undergo remodelling increasing the likelihood of delamination at that site ^68^. Several of our observations are in line with this framework. In the regenerating epicardium, cell density increased during the phase of clonal amplification (Extended Data Fig. 3i), and apical cell area was reduced compared to unoperated hearts (Fig. 3f), consistent with crowding in the epicardial layer. Epicardial density was further increased in cCpE-treated hearts, yet our mechanical characterization revealed a softer and more fluid-like epicardial tissue (Fig. 4, 6). Tissue fluidity is associated with cell-cell rearrangements within a layer, suggesting that crowding in cCpE-treated hearts might be accommodated within the epicardium rather than relieved by cell delamination. Such rearrangements would also separate clonally related cells, consistent with the tendency towards higher fragmentation in our mathematical model and with the reduction in clone size despite decreased delamination. Although challenging, live imaging of the regenerating heart would resolve whether epicardial cells exchange neighbours more readily in cCpE treated hearts.

EPDCs emerging from an epicardial layer in which clonal amplification was impaired failed to activate the putative regenerative gene module and to acquire the regenerative EPDC state. The requirement for clonal amplification, rather than dispersed cell division, suggests that cell state transitions leading to the generation of a migratory EPDC depend on cellular neighbourhoods. This is akin to embryonic induction, where differentiation can depend on a critical mass of interacting cells, a phenomenon termed the community effect ^66^. Cells within an epicardial clone are neighbours originating from a common progenitor, coupled mechanically through shared cell-cell junctions and biochemically through contact-dependent and short-range paracrine cues. Amplification would thus generate a patch of like neighbours in which a new cell state can be established and maintained, making the community it produces, rather than lineage ancestry per se, the essential feature of clonal expansion.

A key question emerging from our work is how the molecular composition of tight junctions is coupled to the cellular decisions that govern clonal expansion and the generation of migratory EPDCs. The epicardium retains a continuous network of ZO-1⁺ cell–cell contacts following cCpE treatment, so regenerative competence depends on the molecular composition of otherwise intact junctions. The C-terminal cytoplasmic tails of CLDNs act as intracellular docking sites for signalling and scaffolding proteins ^70,71^. Changes in junctional CLDN composition could therefore influence not only the structural properties of cell–cell contacts, but also the signalling environment those contacts support. Consistent with this, perturbation of tight junction composition dysregulated genes associated with Ephrin, Hippo/YAP and ERK signalling rather than selectively affecting a single pathway (Fig. 5j). Nuclear phospho-ERK1/2 levels were nonetheless unchanged by cCpE treatment, suggesting that CLDNs do not act through nuclear ERK activity (Extended Data Fig. 6a). These signalling pathways have established roles in epicardial biology, and our results suggest an important upstream role for cCpE-sensitive CLDNs in orchestrating them ^72–76^. Notably, junctional CLDN6 content varies markedly between the individual contacts of a single epicardial cell during regeneration (Fig. 3). Signalling potential may therefore be distributed unevenly around the cell perimeter rather than being a uniform cellular property, which would allow cells to distinguish among their individual interfaces.

Decreased EPDC presence was accompanied by reduced cardiomyocyte protrusion into the injury site, a process previously attributed largely to permissive ECM remodelling, including MMP14-dependent collagen degradation^63,77,78^. Here, TNC abundance was preserved while the TNC-rich bridges were no longer detectably organised (Fig. 8). EPDCs may therefore contribute to muscle regeneration in part by organising ECM at the injury site into structures that guide directional cell movement, analogous to the hyaluronan-rich scaffolds that guide coronary growth ahead of the vascular front ^76^.

The mammalian epicardium also harbours CLDN-containing tight junctions, yet their molecular composition and function in response to injury remain largely unexplored. Whether tight junction composition similarly constrains epicardial behaviour in the mammalian heart remains to be tested. Efforts to harness the mammalian epicardium for cardiac repair have largely focused on reactivating or augmenting its injury-induced developmental programme ^80^, yet our work indicates that activation is a poor predictor of regenerative output: activation was widespread in response to injury, while only a minority of epicardial cells expanded clonally prior to generation of regenerative EPDCs. The 149 genes comprising the regenerative gene module we define (Fig. 7) provide a shortlist of candidates for assessing whether the corresponding state can be induced in mammalian EPDCs.

## Acknowledgements

We would like to thank members of the Eroglu and Chara Labs, Carlos Mario Ayala Torres and András Simon for scientific discussions, Florian Salomons for imaging support, Elin Arvidsson, Chad Donaldson, Nino Gopi Daisy Bai, Stefania Bauer, and Monica Morin for help with raising the animals and Roy Francis for support with spatial transcriptomics analysis. The authors acknowledge support from the National Genomics Infrastructure in Stockholm funded by Science for Life Laboratory, the Knut and Alice Wallenberg Foundation and the Swedish Research Council, and NAISS for assistance with massively parallel sequencing and access to the UPPMAX computational infrastructure. This work was supported by Swedish Research Council (2022-01365), Knut and Alice Wallenberg Foundation (2023.0078), ERC Starting Grant (Tightly Controlled, 101165259), Åke Wibergs Foundation (M22-0065, M23-0055) and Karolinska Foundations (2023-01243, 2024-02555). A.S.C. and O.C. were funded by a BBSRC grant BB/X014908/1 to O.C. O.C. was also funded by UADE grant P26T02 to O.C.

## Code Availability

The original scripts of the key bioinformatics analyses are available via GitHub at https://github.com/Elif-Eroglu-Lab/cCpE_paper_2026.git. All custom codes developed for Forsys analysis and mathematical modelling in this study are publicly available on Github (https://github.com/delfiperez/Claudin-dependent_clonal_expansion_couples_epicardial_activation_to_regenerative_fate_acquisition) and Zenodo (https://doi.org/10.5281/zenodo.23068476). Forsys is available on Github (https://github.com/borgesaugusto/forsys) and on Zenodo (https://doi.org/10.5281/zenodo.11282554). Any additional information required to re-analyse the data reported in this paper is available from the lead contact upon request.

---

**Extended Data Fig. 1, related to Figures 1 and 2.**
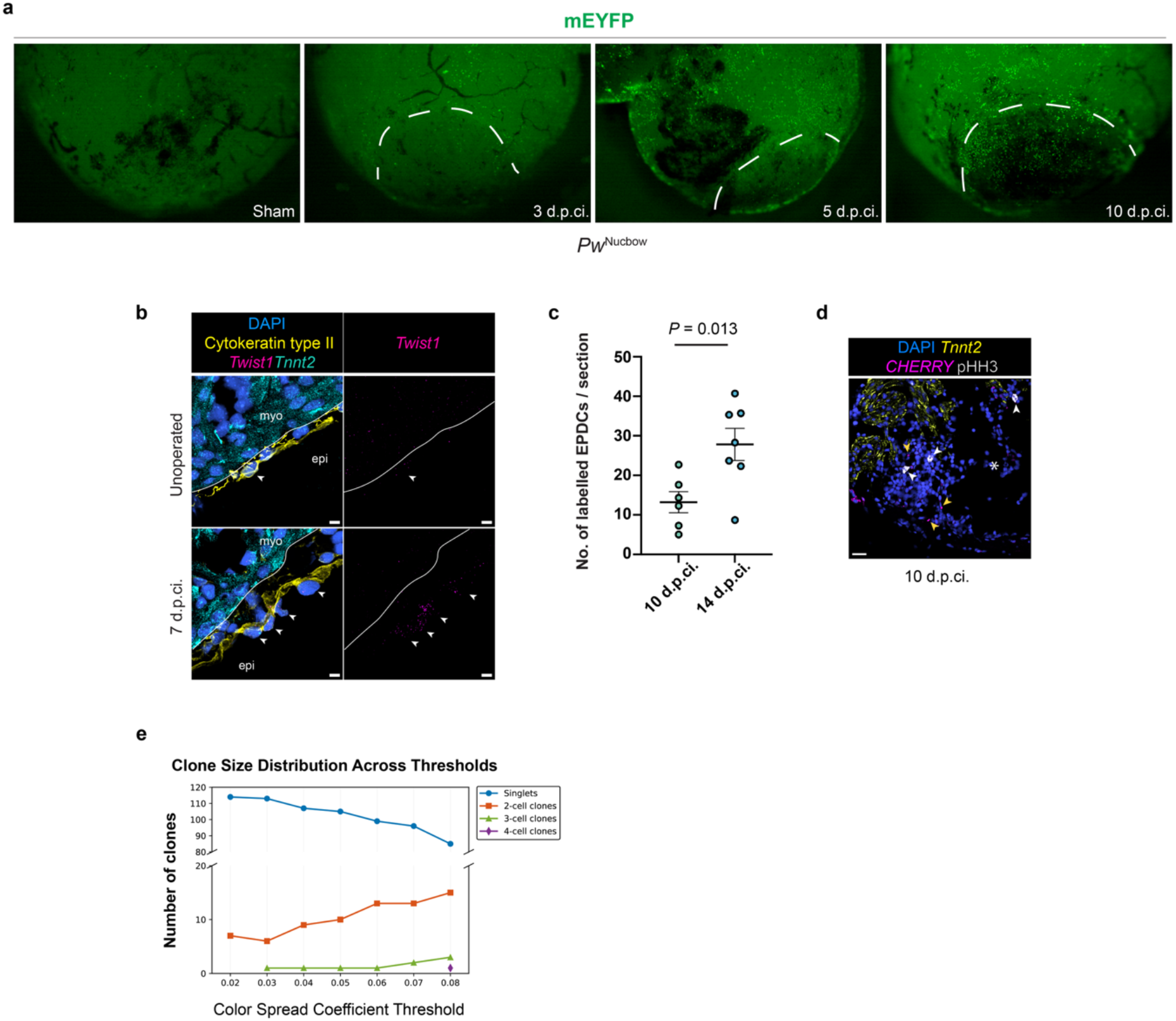
**a,** Representative images of sham-operated or cryoinjured TAT-Cre injected *Pw^Nucbow^* hearts at 3, 5 and 10 d.p.ci. Endogenous mEYFP fluorescence is shown. Approximated injury area is demarcated by dashed lines. **b,** Representative images of RNAscope *in situ* hybridization against the epithelial-to-mesenchymal transition marker *Twist1* and the cardiomyocyte marker *Tnnt2* in unoperated and cryoinjured hearts at 7 d.p.ci. Cytokeratin type II labels the epicardial layer, and DAPI labels nuclei. Solid lines indicate the boundary between the epicardium (epi) and myocardium (myo). Arrowheads indicate epicardial cells. Scale bars, 10 μm. **c,** Number of TAT-Cre labelled EPDCs at the injury site at 10 and 14 d.p.ci. Data represent individual animals. Welch’s t-test. **d,** Representative image of the injury site in TAT-Cre-injected GFP-loxP-Cherry reporter newts at 10 d.p.ci. RNAscope *in situ* hybridization against *Tnnt2* labels cardiomyocytes and CHERRY labels lineage-traced epicardial cells. White arrowheads indicate pHH3⁺ cells, and yellow arrowheads indicate Cherry⁺ EPDCs. Asterisk marks the injury site. Scale bar, 50 μm. **e,** Distributions of clone size across colour spread coefficient thresholds ranging from 0.02 to 0.08 in sham-operated hearts.

**Extended Data Fig. 2, related to Figure 3.**
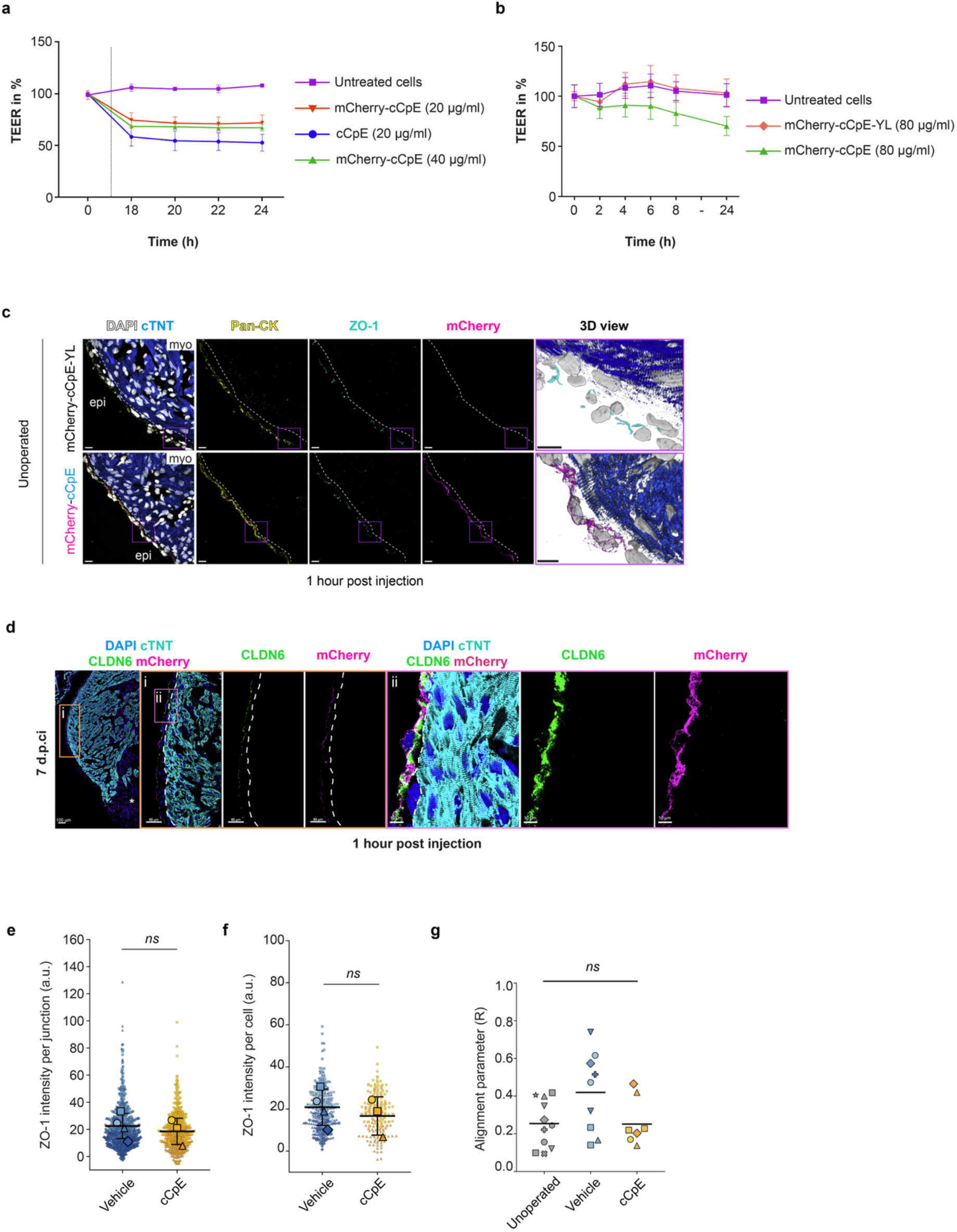
**a,** Transepithelial electrical resistance (TEER) measurements of untreated, cCpE- or mCherry-cCpE–treated Caco-2 monolayers. **b,** TEER measurements of Caco-2 monolayers treated with mCherry-cCpE or CLDN binding deficient mutant mCherry-cCpE-YL (Takahashi et al. 2008). **c,** mCherry-cCpE targets *P. waltl* epicardium. Representative immunostaining images of DAPI, cTnT, Pan-CK, ZO-1 and mCherry in animals intraperitoneally injected with mCherry-cCpE-YL and mCherry-cCpE and analyzed 1 h post-injection. Overview images (left) are shown as maximum intensity projections. Higher-magnification views of the region outlined by the magenta box are displayed as 3D views (right). mCherry-cCpE-YL: *n* = 3 animals; mCherry-cCpE: *n* = 8 animals. Scale bars, 20 μm (max intensity projections), 10 μm (3D views). Dashed lines indicate the myocardium (myo)-epicardium (epi) boundary. **d,** Representative immunostaining images of DAPI, cTnT, CLDN6, and mCherry in animals injected intraperitoneally with mCherry-cCpE at 7 d.p.ci, 1 h post-injection. Higher magnification views of the regions outlined by the orange (i) and magenta (ii) boxes in the overview images (left) show epicardial cells in which CLDN6 and mCherry-cCpE colocalize. Scale bars, 100 μm (overview), 50 μm (i), 10 μm (ii). **e, f,** Quantification of ZO-1 intensity per cell–cell junction (**e**) and per cell (**f**) in vehicle- and cCpE-treated hearts at 10 d.p.ci. Each point represents one junction (**e**) or one cell (**f**). Vehicle: *n* = 890 junctions and 308 cells; cCpE: *n* = 470 junctions and 162 cells. Data were collected from 3-4 animals per condition. Large symbols denote animal means. Values are background-subtracted. Data are the mean ± S.D. of animal means. Student’s t-test. Per cell-cell junction **(e)** *P* = 0.606; per cell **(f)** *P* = 0.565. **g,** Quantification of the mean resultant length R, the alignment parameter, by condition. Each point is one field of view (FOV) and the point shape identifies the animal. Unoperated, *n* = 11 FOV from 6 animals and; vehicle, *n* = 11 FOV from 6 animals and; cCpE, *n* = 7 FOV from 4 animals. Horizontal bars are group means. One-way ANOVA, *P* = 0.063.

**Extended Data Fig. 3, related to Figure 4.**
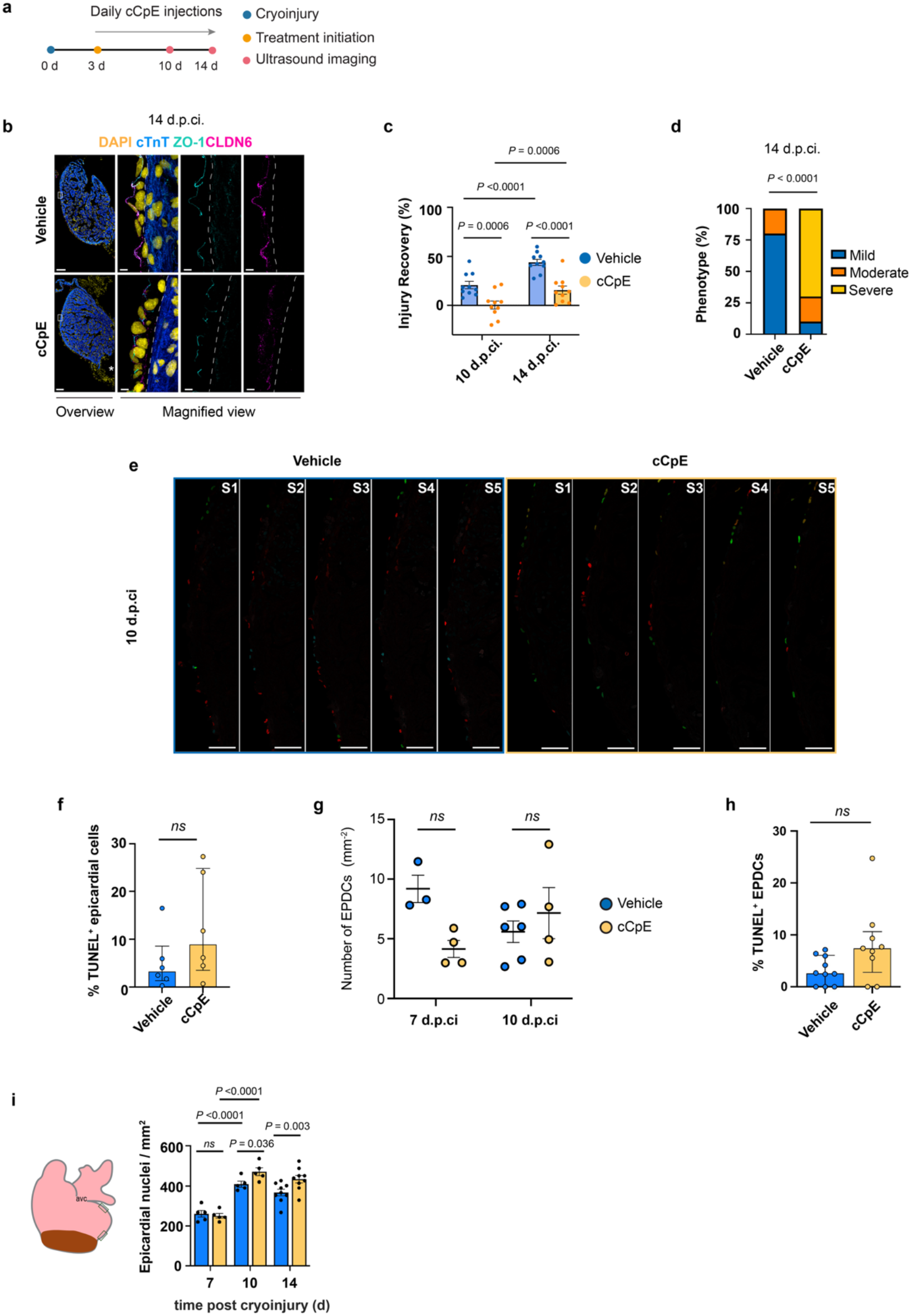
Prolonged cCpE treatment disrupts junctional localization of CLDN6. **a,** Schematic of the prolonged *in vivo* cCpE treatment. **b,** Representative images of DAPI, cTnT, ZO-1, and CLDN6 immunostaining in frontal heart sections from vehicle- or cCpE-treated hearts at 14 d.p.ci. Higher-magnification views of the region outlined by the white box show side views of epicardial tight junctions marked by ZO-1 and CLDN6. In cCpE-treated hearts, CLDN6 appears as cytoplasmic puncta. Scale bars, 200 μm (overview), 10 μm (magnified view). **c,** Recovery from injury was monitored in vehicle- and cCpE-treated animals. The injury area was calculated for each animal by echocardiography at the indicated time points and normalized to the pre-treatment injury size at 3 d.p.ci. Data are the mean ± s.e.m. Two-way ANOVA. **d,** Stacked bar plot showing the proportion of regeneration phenotypes classified as mild (<60%), moderate (60-80%) or severe (>80%) at 14 d.p.ci. in vehicle- and cCpE-treated hearts. Fisher’s exact test. **e,** Representative serial sections (S1-S5) of TAT-Cre labelled hearts from *Nucbow* reporter salamanders at 10 d.p.ci, treated with vehicle and cCpE. Scale bar, 100 μm. **f,** Quantification of TUNEL⁺ epicardial cells in vehicle- and cCpE-treated hearts at 10 d.p.ci. Points represent individual animals. Mann-Whitney test, *P* = 0.180. **g,** Quantification of number of EPDC at the injury site at 7 and 10 d.p.ci. in vehicle- and cCpE-treated hearts, normalized to the size of the injury. Data represent individual animals. Data are the mean ± s.e.m. Šídák’s multiple comparisons test. 7 d.p.ci; *P* = 0.511, 10 d.p.ci; *P* = 0.718. **h,** Proportion of TUNEL⁺ EPDCs at the injury site in vehicle- and cCpE-treated hearts at 10 d.p.ci. Data represent individual animals; lines indicate the median and interquartile range. 586 EPDCs in *n* = 10 animals, and 465 EPDCs in *n* = 6 animals were analysed from vehicle and cCpE-treated hearts respectively. Mann-Whitney U-test, *P* = 0.0514. **i,** Left, schematic of a ventricular section indicating the epicardial regions in which cell density was quantified, adjacent to the atrioventricular canal (avc) and at the injury border. Grey, epicardium; brown, injured myocardium. Right, density of epicardial nuclei within the Pan-CK⁺ layer at 7, 10 and 14 d.p.ci. in vehicle- and cCpE-treated hearts. Nuclei were counted in two rectangular regions of interest per section and pooled. Bars show mean ± s.e.m.; points represent individual animals. *n* = 5, 5, 6 animals per group at 7, 10 and 14 d.p.ci. respectively. Mixed effect analysis (REML) with Šídák’s multiple comparisons test.

**Extended Data Fig. 4, related to Figure 5.**
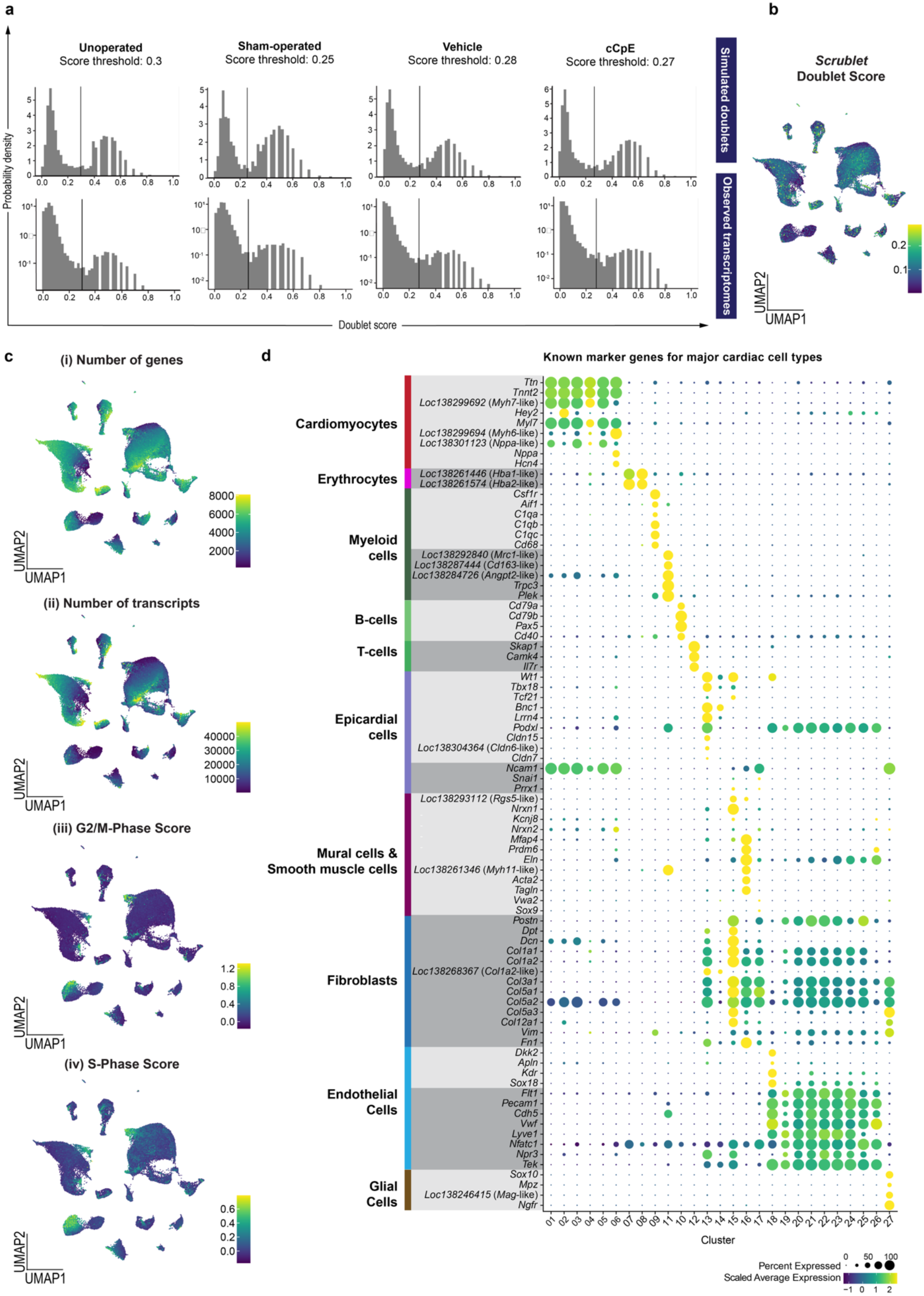
**a**, Histograms showing *Scrublet* doublet score distributions for simulated doublets and observed transcriptomes. Vertical line shows the score threshold used to filter out putative doublets. **b,** UMAP representation of cells from unoperated, sham-operated, vehicle- and cCpE-treated conditions, coloured by Scrublet score. **c**, UMAP representation of all cells coloured by (i) number of genes, (ii) number of transcripts, (iii) G2-M phase score, and (iv) S-phase score **d,** Dotplot showing canonical marker genes for major cardiac cell types and their expression across identified cell clusters.

**Extended Data Fig. 5, related to Figure 5.**
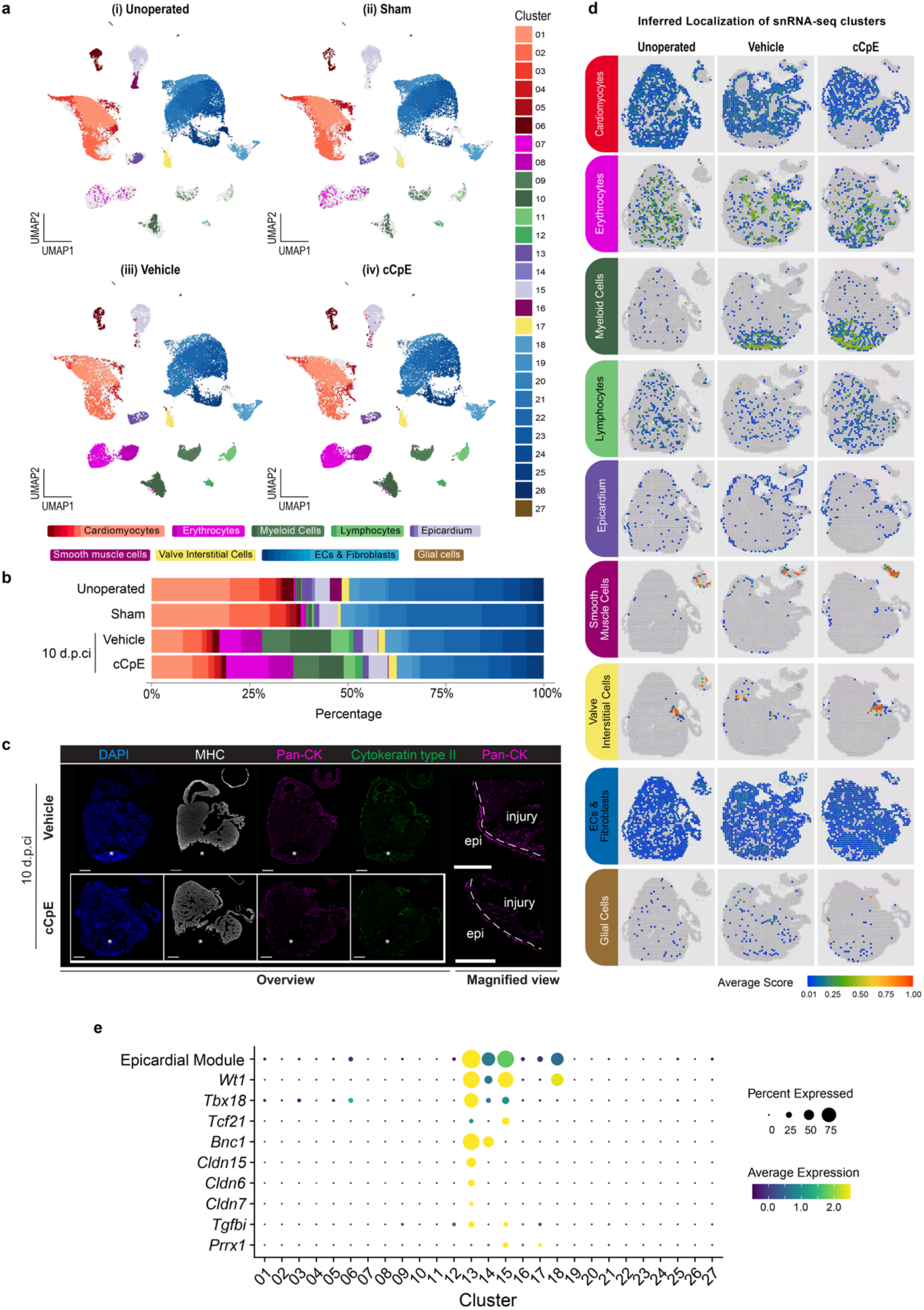
**a,** UMAP representation of nuclei from (i) unoperated, (ii) sham-operated, (iii) vehicle- and (iv) cCpE-treated hearts, coloured by cluster. Shades represent clusters belonging to the same major cardiac cell type. **b,** Bar plot showing the proportions of the identified clusters within each condition. **c,** Representative immunofluorescence images of fresh-frozen vehicle- and cCpE-treated hearts at 10 d.p.ci., stained for DAPI, MHC, Pan-CK and Cytokeratin type II. Magnified views of the apical injury region show a continuous epicardial layer (outside the dashed line) covering the injury site in both conditions. Sections analysed by immunofluorescence were adjacent to those used for spatial transcriptomics. Asterisks indicate the injury site. Scale bars, 500 μm. **d,** Inferred localization of major cardiac cell types from snRNA-seq based on integration with spatial transcriptomics. Spot intensity shows predicted score averaged for clusters belonging to the same major cardiac cell type. Spots with score < 0.01 are coloured in grey. **e,** Dot plot showing the expression of the epicardial gene module across all clusters, as well as the individual genes comprising the module.

**Extended Data Fig. 6, related to Figure 5, 6.**
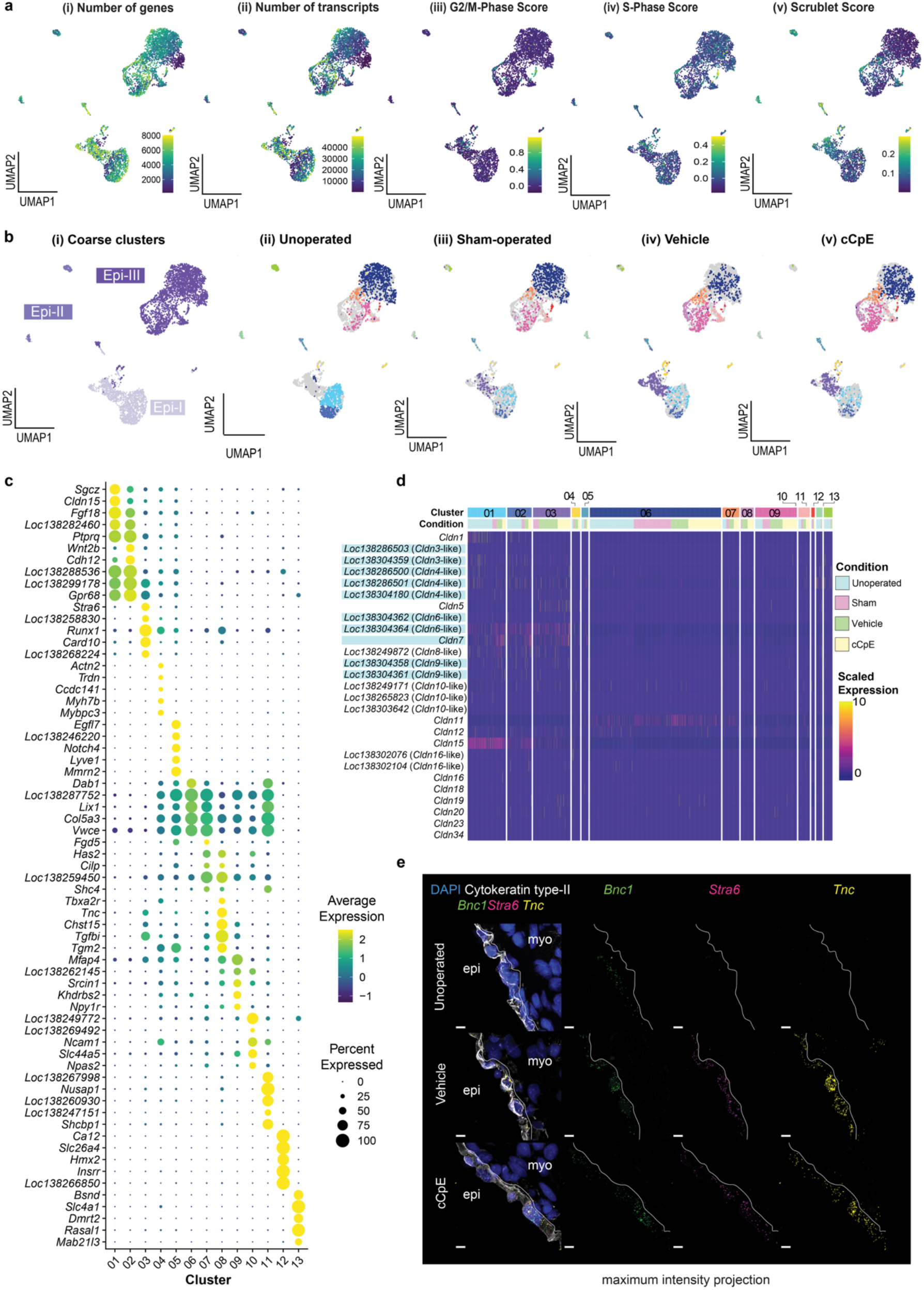
**a,** UMAP representation of subclustered epicardial cells coloured by (i) number of genes, (ii) number of transcripts, (iii) G2-M phase score, (iv) S-phase score, and (v) *Scrublet* doublet score. **b,** UMAP representation of nuclei from epicardial lineage cells (i) coloured based on cluster identity from the full dataset. (ii-v) show the nuclei from unoperated, sham-operated, vehicle-treated, and cCpE-treated animals, coloured by cluster identity. **c,** Dotplot showing top 5 marker genes for each cluster arranged by a combination of log2 fold change and adjusted *P*-value. **d,** Heatmap showing expression of Cldn family genes present in the dataset across clusters, stratified by condition. Reported cCpE-responsive Cldns are highlighted ^36^. **e,** Representative maximum intensity projection images of RNAscope *in situ* hybridization and immunostaining against the epicardial marker *Bnc1*, activated epicardium marker *Stra6*, DAPI, Cytokeratin type II, and *Tnc* in unoperated, vehicle-treated, and cCpE-treated hearts at 10 d.p.ci. Dashed lines indicate the epicardial–myocardial boundary. Scale bars, 10 μm.

**Extended Data Fig. 7, related to Fig. 6.**
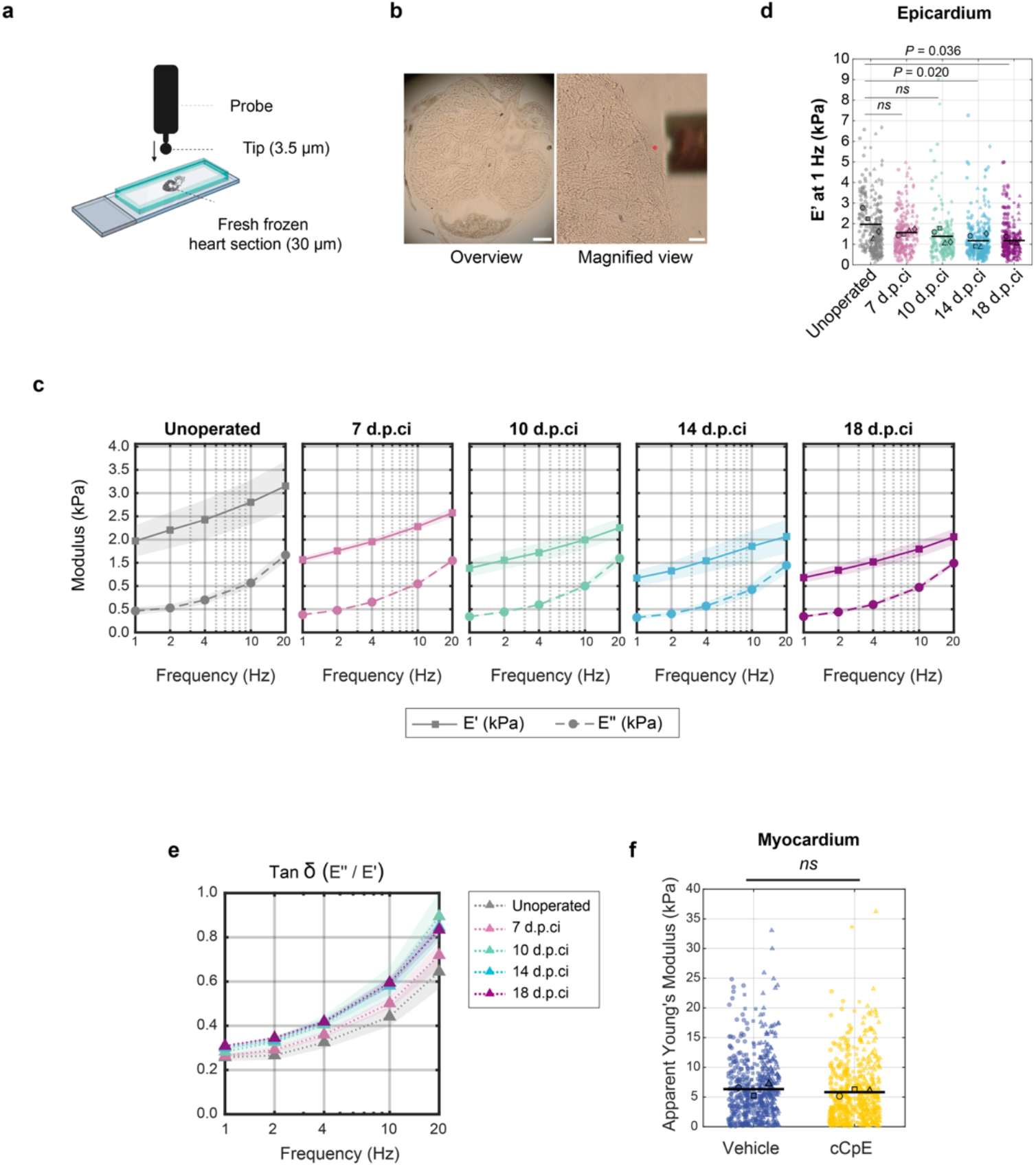
**a**, Schematic showing the nanoindentation setup. **b**, Representative brightfield images of a heart section selected for indentation (left) and a magnified view with the epicardium and the indentation probe (right). Scale bars, overview 500 μm, magnified view 100 μm. **c**, Time-dependent viscoelastic behaviour of the regenerating epicardium characterized over a frequency sweep (1,2,4,10,20 Hz). Storage modulus (E’, solid lines) and loss modulus (E’’, dashed lines); mean ± s.e.m. **d,** Scatter plot of storage modulus (E’) measured at 1Hz (kPa) in the epicardial layer across unoperated, 7, 10, 14 and 18 d.p.ci hearts. Each small point represents an individual measurement; each animal is denoted by a distinct symbol, with horizontal black lines indicating the group mean. Comparisons to the unoperated control were performed using linear hypothesis tests on the fixed-effects coefficients of a linear mixed-effects model, with animal as a random-effect and condition as a fixed-effect. (Satterthwaite F-test, Bonferroni-corrected for four comparisons; unoperated vs. 14 d.p.ci., *P* = 0.020; unoperated vs. 18 d.p.ci., *P* = 0.036; unoperated vs. 7 and 10 d.p.ci., *P* = 0.526 and *P* = 0.130. **e,** Variation of loss tangent (tanδ dotted lines) with frequency overlaid for all timepoints on a single plot. **f,** Graph showing the apparent Young’s modulus of vehicle- and cCpE-treated heart myocardium at 10 d.p.ci. Each data point represents an individual measurement, and each animal is denoted with a different symbol. Animal averages are shown as bigger symbols. *n* = 3 animals per group. Mann-Whitney U-test, *P* = 0.400.

**Extended Data Fig. 8, related to Figure 7.**
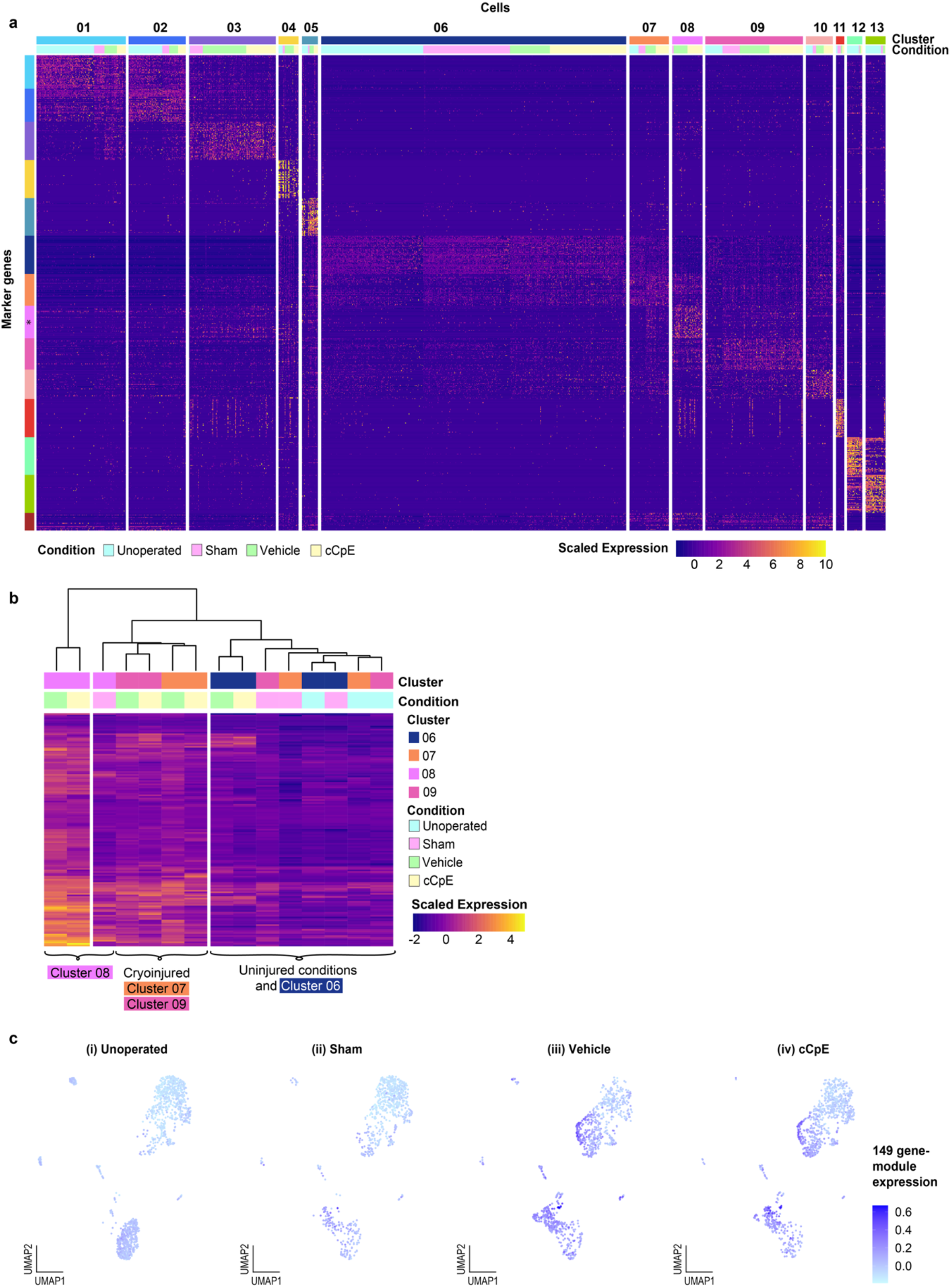
**a,** Heatmap showing the scaled expression of each cluster’s top 50 markers based on log2 fold change. Markers identified among the top markers in more than one cluster are consolidated at the bottom of the heatmap (dark brown). **b**, Heatmap showing aggregated expression of genes in the putative regenerative gene module across clusters 06–06 and experimental conditions. **c,** UMAP representation of the Seurat module score for the putative regenerative gene module, shown separately for cells of epicardial lineage in (i) unoperated, (ii) sham-operated, (iii) vehicle-treated, and (iv) cCpE-treated hearts.

**Extended Data Fig. 9, related to Figure 7.**
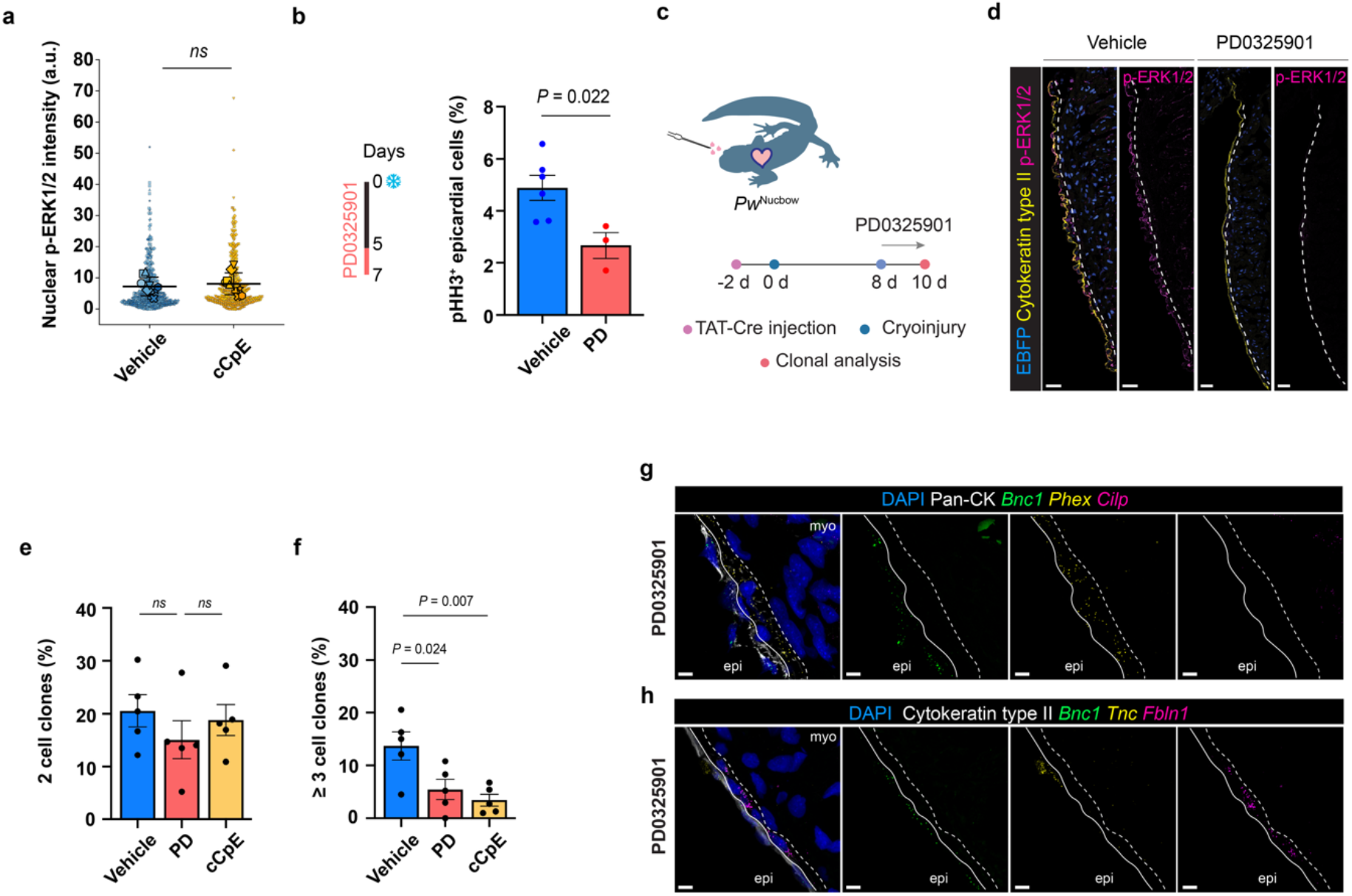
Inhibition of ERK1/2 phosphorylation impairs cell proliferation and clonal expansion. **a,** Quantification of nuclear phospho-ERK (p-ERK) intensity in 465 epicardial nuclei in vehicle- and 425 nuclei in cCpE-treated hearts across *n* = 6 animals in each group. Small symbols denote individual nuclei; large symbols are animal means; bars, mean ± S.D. of animal means. Student’s t-test, *P* = 0.588. **b,** Quantification of pHH3^+^ epicardial cells on vehicle- and PD0325901 (PD)-treated animals at 7 d.p.ci. Welch’s t-test. **c,** Schematic showing the experimental design for assessing epicardial clonal dynamics upon inhibition of ERK1/2 phosphorylation. **d,** Representative images of DAPI, Cytokeratin type II and p-ERK1/2 immunostaining in vehicle- or PD-treated hearts at 10 d.p.ci. Dashed lines indicate the epicardium-myocardium border. Scale bars, 50 μm. **e,** Proportion of 2 cell clones in vehicle-, PD- and cCpE-treated hearts at 10 d.p.ci. **f,** Proportion of clones containing more than 3 cells in vehicle-, PD- and cCpE-treated hearts at 10 d.p.ci. **(e, f)** Data represent individual animals, *n* = 5 animals for all conditions. One-way ANOVA with Dunnett’s multiple comparison test. Vehicle vs. PD: *P* = 0.358, vehicle vs. cCpE: >0.66 **(e). g-h,** Representative images of RNAscope *in situ* hybridization against *Bnc1*, *Phex* and *Cilp*. **(g)** and *Bnc1*, *Tnc* and *Fbln1* **(h)** in PD0325901-treated hearts at 10 d.p.ci. Pan-CK and Cytokeratin type II delineate the epicardial and subepicardial regions. Solid lines indicate the boundary between the epicardial and subepicardial compartments, whereas dashed lines indicate the boundary between the sub-epicardium and myocardium. Maximum-intensity projections are shown. Scale bars, 10 μm.

**Extended Data Fig. 10, related to Figure 5, 6, 7.**
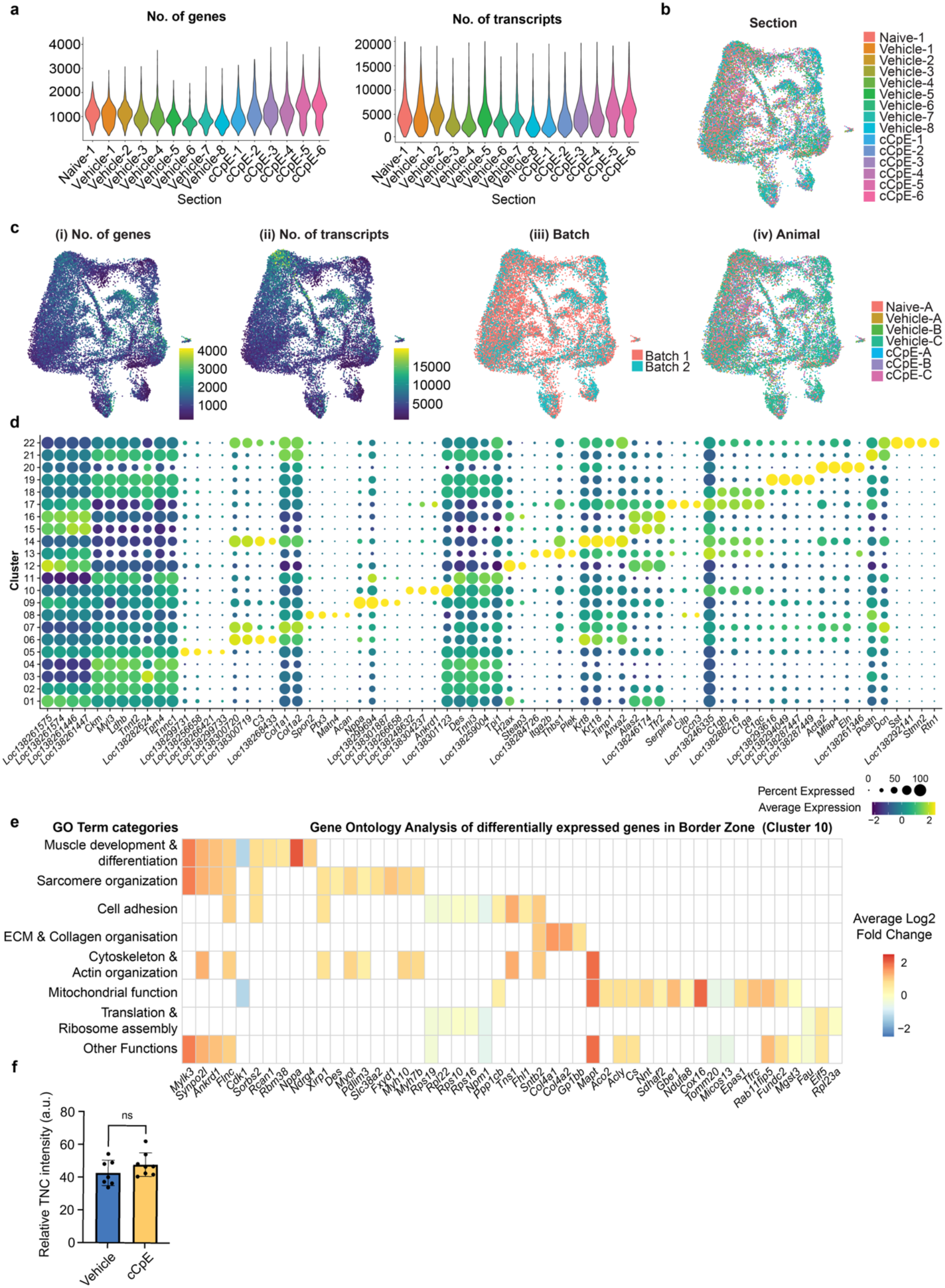
**a,** Violin plots showing the distribution of the number of genes (nfeature) and number of transcripts (nCount) for each spatial section. **b,** UMAP representation of 10x Visium spots coloured by section of origin **c,** (i) number of genes, (ii) number of transcripts, (iii) batch, and (iv) animal of origin. **d,** Dotplot showing expression of top 5 marker genes per cluster. **e,** Heatmap showing all categories summarizing significantly overrepresented GO terms for genes differentially expressed between vehicle- and cCpE-treated hearts in the border zone (Cluster 10) (adjusted *P* < 0.05, q < 0.05). Genes comprising these GO terms are coloured by their log2 fold changes. **f,** Graph showing the quantification of relative TNC intensity in vehicle- vs. cCpE-treated hearts at 14 d.p.ci. Points represent individual animals, *n* = 7 for vehicle-, and *n* = 8 for cCpE-treated hearts. Statistical significance was determined using an unpaired t-test with Welch’s correction. *P* = 0.206.

## Methods

### Animals

All the procedures in this study related to animal handling, care and treatment were performed according to the guidelines approved by the Jordbruksverket/Sweden under the ethical permit numbers 18160-18 and 6857-2024. *P. waltl* were bred and maintained in our aquatic animal facility. The transgenic lines tgTol2(*CAG:loxP-GFP-loxP-Cherry*)^Sinon^ and tgTol2(*CAG:Nucbow*)^Sinon^ were described previously ^1^.

### Cryoinjury

Post-metamorphic salamanders, 7-12 centimeters, were anesthetized by immersion in 0.1% MS222 (A5040, Sigma) and placed in a supine position. A skin incision was made and pericardium was cut open. The heart was maneuvered out of the pericardial cavity. Excess moisture was removed with a tissue. A liquid nitrogen-cooled copper filament (1.2 mm) was applied to the ventricle apex for 10 seconds. Afterwards, the heart was placed back into the pericardial cavity. Pericardium and the outer skin were sealed with Ethilon 8-0 black monofilament sutures (2808G, Ethicon). Animals were placed in 0.5% sulfamerazine solution (S0800, Sigma) on ice overnight and transferred to regular water the next day. 10-days post cryoinjury (d.p.ci) water temperature was raised to 25°C.

### Pericardial injections of the TAT-Cre recombinase

Animals were anesthetized in 0.1% MS222 (A5040, Sigma). 100 µM TAT-Cre recombinase (SCR508, Millipore) diluted in 10 µl aHBSS (80% HBSS, 14175095, Gibco) was microinjected (WPI PV850) into the pericardial sac of the reporter salamanders. Animals were kept in 25°C water for two days after the microinjection. For labelling combined with the injury, animals were subjected to cryoinjury 40-48h post injection (hpi).

### Treatment of salamanders with cCpE and its variants

Salamanders were injected intraperitoneally with 2.5 μg/g Strep-Tag®II-tagged cCpE daily according to experimental scheme (Day 3-6 when harvesting at 7 d.p.ci, Day 3-6 when harvesting at 10 d.p.ci and Day 3-13 when harvesting at 14 d.p.ci). To confirm that cCpE targets epicardium, Strep-Tag®II-tagged mCherry-cCpE and Strep-Tag®II-tagged mCherry-cCpE-Y306A/L315A were injected intraperitoneally at a concentration of 2.5 μg/g 1h before harvesting the heart.

### Treatment of salamanders with PD0325901

PD0325601 was injected intraperitoneally at a concentration of 5 μg/g daily according to experimental scheme (Day 5-7 when harvesting at 7 d.p.ci, Day 8-10 when harvesting at 10 d.p.ci, last injection 6 hours before harvesting the heart).

### Tissue processing for histological analysis and nanoindentation

Animals were anesthetized in 0.1% MS222. For immunostaining, hearts were excised, washed in 70% PBS with heparin (100 units/ml, H4784, Sigma) and fixed overnight at 4°C with 4% paraformaldehyde (J16643.K2, Thermo), then rinsed three times before being exposed to 15% and 30% sucrose (S0386, Sigma) solutions at 4°C. Afterwards, they were equilibrated with O.C.T. (4583, Tissue-Tek®) and frozen. Tissues were sectioned in the frontal plane at 14 µm using a cryostat (Cryostar NX70, Thermo) and stored at -80°C.

For nanoindentation, hearts were excised, rinsed in 70% PBS with heparin and placed in a cryomold with O.C.T. Then, the cryomold was gently placed in an isopentane bath (78-78-4, VWR) cooled in liquid nitrogen for fast and uniform freezing. Tissues were sectioned in the frontal plane at 30 µm using a cryostat and stored at -80°C.

### Staining Procedures

For immunofluorescent staining, tissue sections were fixed in 4% paraformaldehyde for 5 min at RT and washed three times with PBS, 5 min each. Tissues were permeabilized with PBS containing 0.25 % Triton X-100 (T8787, Sigma) for 10 min, rinsed in PBS and blocked in 1% bovine serum albumin (A7906, Sigma), 5% normal goat serum (31873, Invitrogen), 0.1% Triton X-100 in PBS (PBS-T) for 1 hour at RT. Primary antibodies in blocking solution were incubated overnight at 4°C. Next day, sections were washed three times with 0.1% PBS-T for 10 min each. Secondary antibodies were diluted in blocking solution and incubated for 2 hours at RT. For nuclear staining, the sections were incubated with DAPI (2 µg ml^-1^, D6542, Sigma) for 10 min at RT, followed by five washes with PBS-T and one wash with PBS, 5 min each. Slides were rinsed in Milli-Q water and mounted with DAKO fluorescent mounting medium (S3023, Agilent). For pHH3 and CLDN6 staining, heat-induced epitope retrieval was performed in 1x citrate buffer (C6666, Sigma) at 60-65 °C for 10 min before permeabilization and subsequent immunostaining. Apoptotic cells were detected using Click-iT™ Plus TUNEL Assay Kits for In Situ Apoptosis Detection (C10619, Invitrogen). For picrosirius red staining, tissue sections were fixed in 4% paraformaldehyde for 5min at RT, washed three times in Milli-Q water (5min each), followed by incubations in ethanol (100%, 100%, 70%, 50%) for 2min each. Then, they were rinsed in Milli-Q water and incubated with Picrosirius Red (ab246832, Abcam) for 1h at RT, followed by three washes in 0.5% glacial acetic acid (A6283, Sigma) for 2min each. Finally, tissues were dehydrated in an ethanol gradient (70%, 100%, 100%, 2min each) before being cleared in xylene (214736, Sigma) and mounted in VectaMount™ Permanent Mounting Medium (H-5000, VectorLabs). For whole mount immunostaining, hearts were harvested and fixed overnight in 4% paraformaldehyde at 4°C, rinsed three times and bleached for 10 min at RT in 6% H_2_O_2_ (H109, Sigma), 0.5% KOH (P1767, Sigma) in PBS, followed by three PBS washes. Then, hearts were cut sagitally and permeabilized in 0.2M glycine (G7126, Sigma), 2% Tx100, 20% DMSO (472301, Sigma) in PBS for 4 days at 37°C on a shaker, washed in PBS and blocked in 1% BSA, 5% NGS, 0.1% Tx100 in PBS. Primary antibodies were incubated in 2% Tween-20 (P1379, Sigma) supplemented with 0.01% sodium azide (S8032, Sigma) at 37°C for 4 days, washed overnight and incubated in secondary antibodies for 4h at 37°C, followed by a 30min DAPI incubation. Tissues were washed thoroughly and dehydrated in an ethanol gradient (30%, 50%, 70%, 100%), rinsed in fresh 100% ethanol and cleared in ethyl cinnamate (8.00238, Sigma). Stained tissues were stored in ethyl cinnamate until imaging.

### Antibodies used

Rabbit anti-pan Cytokeratin (1:250, ab9377, Abcam), rat anti-mCherry (1:200, M11217, Invitrogen), mouse anti-Tenascin-C (1:25, MT1, DSHB), rabbit anti-Tenascin (1:300, AB19013, Chemicon), mouse anti-MHC (1:200, MF-20, DSHB), mouse anti-Troponin-T (1:80, CT3, DSHB), mouse anti-Cytokeratin type-II (1:100, 1h5, DSHB), rabbit anti-phospho-Histone H3 (1:1000, 06-570, Millipore), rabbit anti-CLDN6 (1:50, ab314134, Abcam), mouse anti-ZO-1 (1:100, 33-6100, Invitrogen), mouse anti-Phospho-Myosin Light Chain 2 Ser16 (1:200, #3675, Cell Signaling Technology), mouse anti-α-actinin (1:800, A7811, Sigma), rabbit anti-phospho-ERK (1:150, #9101, Cell Signaling Technology), BD Pharmingen™ Alexa Fluor® 647 mouse anti-E-Cadherin (1:70, 560062, BD Biosciences). Cross-adsorbed Alexa Fluor conjugated secondary antibodies from Invitrogen were used in 1:1000 dilution. Specifically, anti-mouse IgG1 488 (A21121), anti-mouse IgG2a 488 (A21131), anti-mouse IgG2b 488 (A21141), anti-mouse IgM 488 (A21042), anti-mouse IgM 568 (A21043), anti-rat 568 (A11077), anti-rabbit 568 (A11011), anti-mouse IgG1 647 (A21240), anti-mouse IgM 647 (A21238), anti-rabbit 647 (A21245). Cross-adsorbed CF770 conjugated secondary antibodies from Biotium were used at 1:500 dilution. Specifically, anti-mouse IgG1 (#20254), anti-mouse IgG2a (#20264), anti-mouse IgG2b (#20274) and anti-rabbit (#20078).

### RNAscope in situ hybridization

Custom RNAscope probes against *Twist1*, *Tnnt2e, Phex*, *Cilp1*, *Bnc1*, *Fbln1*, *Tnc,* and *Stra6*, were designed and manufactured by Advanced Cell Diagnostics (ACD). RNAscope Multiplex Fluorescent V2 Assay (323110, ACD) was performed on fixed frozen tissue according to manufacturer’s recommendations with the following optimizations: antigen retrieval at 95°C for 10 min and Protease III treatment for 8-10min. After the RNAscope protocol, slides were washed in PBS and incubated in blocking buffer, followed by immunofluorescence protocol as described in the previous section.

### Molecular Cloning of mCherry-cCpE and mCherry-cCpE-YALA

The coding sequence for mCherry was synthesized by IDT, amplified by PCR and cloned into Strep-Tag®II-c-CpE_pET22b(+) (Genscript) and Strep-Tag®II-c-CpE-Y306A/L315A_pET22b(+) (GenScript) using the In-Fusion Snap Assembly kit (Takara Bio). All plasmids were verified by Sanger sequencing.

### Expression and Purification of Strep-Tag®II-tagged mCherry-cCpE, mCherry-cCpE-Y306A/L315A, and c-CPE

BL21-Codon Plus (DE3)-RIL competent cells (230245, Agilent Technologies) were transformed with Strep-Tag®II-mCherry-c-CPE_pET22b(+), Strep-Tag®II - mCherry-c-CPE-Y306A/L315A_pET22b(+) or Strep-Tag®II-c-CPE_pET22b(+) according to manufacturer’s instructions. A 10 ml overnight culture was used to inoculate 1 liter of LB medium (10855001, Gibco) containing 100 µg/ml carbenicillin (C1613, Sigma-Aldrich) and 25 µg/ml chloramphenicol (C0378, Sigma-Aldrich). Protein expression was induced with 1mM IPTG (R1171, Thermo Fisher) when cultures reached an OD_600_ of 0.5-0.7 following incubation at 37°C with vigorous shaking. After 3 hours, cells were harvested by centrifugation at 4000g for 30min at 4°C. Pellets were resuspended in lysis buffer containing 100 mM Tris/HCl, 150 mM NaCl, 1 mM EDTA (2-1003-100, IBA), rLysozyme (60 KU/g cell paste, 71110, Millipore), benzonase nuclease (250 Units/ml, E1014-25KU, Sigma) EDTA-free protease inhibitor (A32655, Thermo), pH 8.0 and kept on ice for 30 min, followed by sonication with Bandelin Sonopuls (Cycle 2, 6 minutes, 50% power output). After centrifugation of lysed cells at 13000g for 30min at 4°C, the supernatant was passed through a Strep-Tactin® 4Flow® high-capacity column (2-1253-001, IBA). Unbound proteins were washed away with 100 mM Tris/HCl, 150 mM NaCl, 1 mM EDTA, pH 8.0 (2-1003-100, IBA), and StrepII-tagged proteins were eluted with 2.5 mM Desthiobiotin (2-1000-025, IBA). The eluate was concentrated using the Pierce Protein Concentrator (10K MWCO, 88517 Pierce) and endotoxins were removed by incubation with TritonX-114 (X114, Sigma). TritonX-114 residuals were removed using high affinity triton binding beads (152-8620, Bio-Rad) and salt removal was performed with Zeba spin desalting columns (86860, Thermo Fisher). Purified Strep-Tag®II-tagged mCherry-cCpE, mCherry-cCpE-Y306A/L315A and c-CPE proteins were kept at -80°C.

### Echocardiography

Serial echocardiography was performed using the Vevo 3100 Imaging System (VisualSonics, Amsterdam, Netherlands) equipped with a high frequency transducer (MX700, 26-71 MHz). Animals were anesthetized with 0.1% MS-222 and placed in supine position (VisualSonics, Amsterdam, Netherlands). B-mode parasternal long axis (PLAX) images were acquired by placing the transducer directly above the ventral side of the animal, parallel to the midline of the chest. For each animal and timepoint, we first obtained B-mode images corresponding to the PLAX plane containing both the ventricle and outflow tract. Within the context of regeneration, additional planes containing the largest cross-sectional area of the injury were obtained for injury size calculations, when necessary. This was achieved by scanning the transducer along the left-right axis using the in-built micromanipulator. Focus depth, 2D gain, image dimensions were optimized according to manufacturer’s recommendations, and at least 4 cardiac cycles were recorded per imaging sequence. After the procedure animals were returned to water to resuscitate. Injury size was calculated by measuring the area of the whole ventricle (excluding outflow tract but including injury area) and the injury itself in Vevo Lab 3.2.0 using the 2D area function. The following formula was used: (injury size / ventricle size) * 100.

### Mathematical Modelling

#### Model of clonal expansion in the regenerating epicardium: cell proliferation and delamination

To explain the temporal change in clone sizes and distribution in the epicardial clonal expansion we first proposed a proliferation and delamination mathematical model (Fig. 4l). Cells form clones which can lose or gain cells as time passes. We will follow a clone of size *i* with *i* being a generic constant which can take the value of any positive integer. We consider that there are two mechanisms which change the clone size: cell division/proliferation and delamination. We assume that cells proliferate or delaminate at rates r and d, respectively. When a cell from one clone of size *i* divides, the two daughter cells remain in the clone, and the clone ends with a size of *i +1*. When a cell delaminates from a clone of size *i*, it will leave the clone (and the tissue) and the clone will obtain a size of *i -1*. We assume that delaminated cells instantly leave the epicardium and reach the injury site. The total rate of proliferation in a clone of size *i* is the sum of the rates of its individual cells, *r* i*. The same argument is valid for delamination, being *d* i* the total rate of delamination. In consequence, the change in number of clones of size *i* will be given by the following equation:

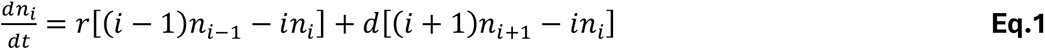

Where *n_i_* is the number of clones of size *i*.

The first term on the right-hand side (rhs) corresponds to the increase in number of clones of size *i* due to divisions of clones of size *i -1*. The second term on the rhs describes the decrease in number of clones of size *i* due to divisions of clones of size *i*, forming clones of size *i+1*. The third term on the rhs corresponds to the increase in the number of clones of size *i* due to the delamination of size *i+1* which lose one cell. The last term on the rhs is the decrease in number of clones of size *i* due to delamination of clones of size *i*, forming clones of size *i-1*.

The mathematical model is finally formed by a system of infinite coupled ordinary differential equations for all possible positive integer values of *i* starting from one. With this model, we define the total number of cells *N* as the sum of the number of cells in each clone multiplied by the number of clones:

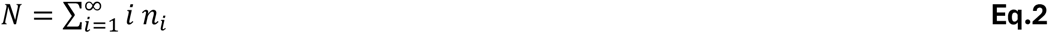

We calculate its time derivative as follows:

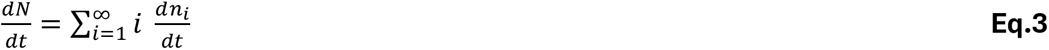

We replace Eq. 1into Eq. 3 to obtain:

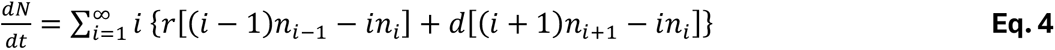

After rearranging the summatory indexes, some of the terms cancel each other and we arrive at:

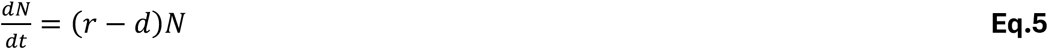

This model predicts that the number of cells increases exponentially in time with exponential rate r -d. As mentioned in the Results section, this is what we observed in Fig. 2h.

We also calculate the total number of clones (*N_c_*):

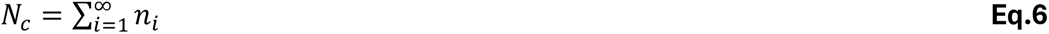

We calculate its time derivative as follows:

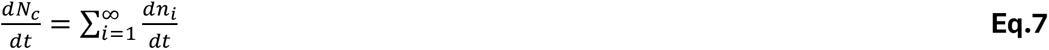

We replace Eq. 1 into Eq. 7 to obtain:

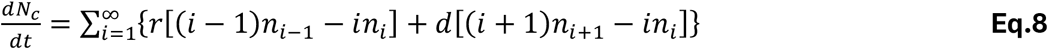

Again, after rearrangement of the summatory indexes, some of the terms cancel each other and we arrive at:

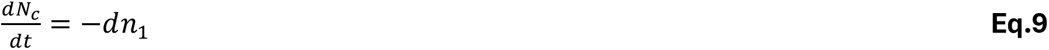

Since the delamination rate d and the number of clones containing only one cell n1 are either positive or, in the trivial case, null, Eq. 6 predicts that the total number of clones either decrease or remain constant in time. This contradicts what it seen experimentally where the number of clones increase in time (see Fig. 4m in the Results section). We deduce that a model assuming proliferation and delamination alone cannot explain the phenomenology observed.

#### Model of clonal expansion in the regenerating epicardium: clone fragmentation, cell proliferation and –delamination

Since the previous model assuming proliferation and delamination alone predicts that the clone number cannot increase, we considered adding a mechanism of clone fragmentation to increase the number of clones over time (Fig. 4l). This model variant works similarly as the previous one in which cells proliferate and delaminate, but now we incorporated the capacity for clone fragmentation. When a clone fragmentates, it divides into two clones of smaller size. We will assume that a clone of size *i* can fragmentate into two clones, one of size 1 and one of size *i -1*. We introduce the rate of fragmentation *f*. The equations for the change in time of clone of size *i* are:

If *i* ≠ 1:

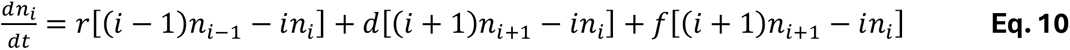

If *i* = 1:

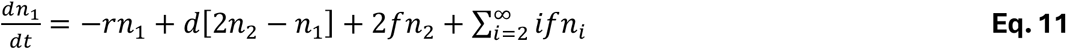

As before, we define the total number of cells N as the sum of the number of cells in each clone multiplied by the number of clones:

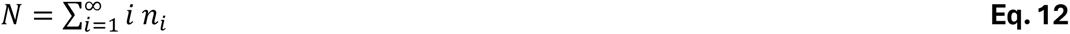

We calculate its time derivative:

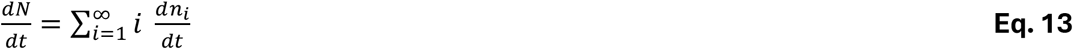

If we combine Eq. 10, Eq. 11 and Eq. 13 we obtain:

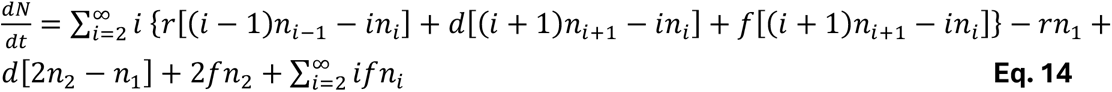

After some rearrangement in the summatory indexes, we arrive at:

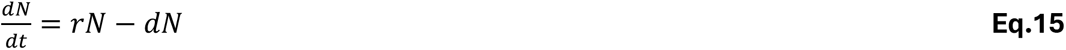

As expected, the fragmentation part of the model, does not change the dynamics of the total number of cells and thus, Eq. 15 is equal to Eq. 5. From the delamination term we see that the source of delaminated cells is *dN*, which we use to obtain the change in the number of delaminated cells *N_d_*:

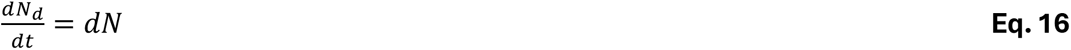

We also calculate the total number of clones (*N_c_*):

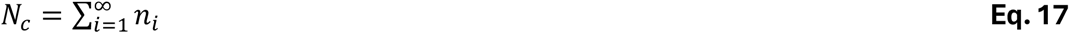

We calculate its time derivative:

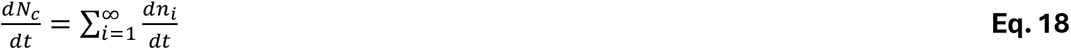

If we combine Eq. 10, Eq. 11 and Eq. 17 we obtain:

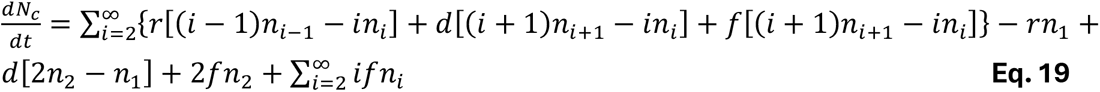

After some rearrangement in the summatory indexes, we arrive at:

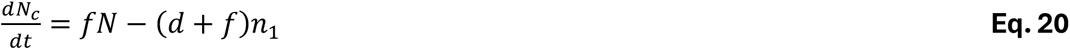

Therefore, this model variant predicts that the number of clones can increase over time, depending on the parameter values.

For the sake of simplicity, we assumed that the delamination (*d*) and fragmentation (*f*) rates are constant. In contrast, we modelled the proliferation rate as time-dependent, *r*(*t*), based on our quantification of pHH3^+^ epicardial cells, which revealed a clear two-phase kinetic profile characterised by an initial increase followed by an abrupt decrease that subsequently remains constant (Fig. 1c). Hence, we modelled the proliferation rate as follows:

If *t* < *t_lim_*:

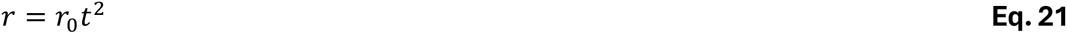

And if *t* ≥ *t_lim_*:

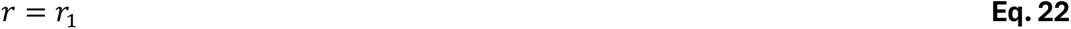

Where *t_lim_* represents the time separating the two kinetic phases, *r*_0_ is a parameter controlling the magnitude of the quadratic increase in proliferation rate and *r*_1_ represents the proliferation rate after *t_lim_*.

### Model fitting and parameter estimation

In this section, we describe the fitting of the model assuming clone fragmentation, cell proliferation and delamination to the experimental data. To reduce the number of the free parameters we focused on those controlling proliferation (Eq. 21 and 22). The parameters *t_lim_* and *r_1_* were estimated from the experimental time course of the percentage of pHH3+ epicardial cells (see Fig. 1C). Since the percentage of pHH3^+^ epicardial cells peaked at day 7 post injury and decreased abruptly by day 10, we set *t_lim_* = 9 *days* (see Eq. 21 and Eq. 22). By combining Eq. 21 and 22 and using the percentages of pHH3^+^ epicardial cells immediately before and after *t_lim_*, we obtain: 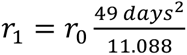 Thus, *r* is determined through this expression by the parameter *r*_0_, which is one of the free parameters estimated by fitting the model to the experimental data (see below).

We simultaneously fitted the model to three experimental datasets: the time course of the number of labelled epicardial cells (*N*), the number of clones (*N_c_*), and the number of EPDCs, which was used as a proxy for the cumulative number of delaminated cells (*N_d_*). The model contains three free parameters: *r*_0_, *d* and *f*, corresponding to cell proliferation, cell delamination and clone fragmentation, respectively. Parameter estimation was performed in Python using the differential_evolution function from the Scipy library ^2^ and the following custom objective function:

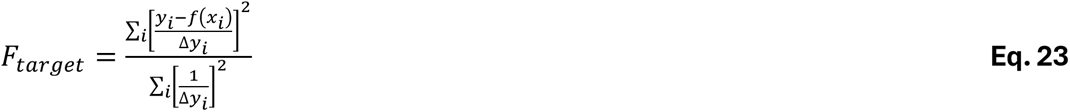

Where *y_i_* is an experimental measurement, Δ*y_i_* its uncertainty, and*f*(*x_i_*) the corresponding model prediction. The summation was performed over all experimental measurements of the number of labelled epicardial cells, the number of clones, and the number of EPDCs. Dividing by Δ*y_i_* weighted the fit towards experimental data points with lower relative uncertainty.

The total number of labelled epicardial cells serves as the source population for the delaminated cells at the lesion site (Eq. 16). In our clonal expansion experiments, we measured approximately one quarter of the total labelled epicardial cells contributing to the pool of delaminated cells. To account for this difference, we introduced a correction factor of 4 into Eq. 16. The modified equation used for the model fitting is:

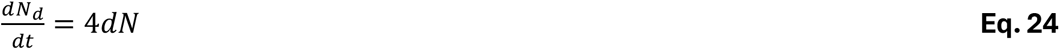

To quantify parameter uncertainty and propagate variance into model predictions, we implemented a custom bootstrap procedure. Experimental measurements were grouped into two independent datasets: one containing EPDC cell counts (*N_d_*) and another containing the coupled measurements of *N* and *N_c_* obtained from the same animal. At each iteration, one of the two datasets was selected at random, and a single experimental record at a specific time point was removed and replaced by a randomly selected record from the same dataset. Thus, each iteration introduced a single-sample perturbation to the experimental dataset, after which the model was refitted. This procedure was repeated iteratively until convergence was reached. Convergence was assessed from the percentile bootstrap 95% confidence intervals for the three fitted parameters (*d*, *r*_0_, and *f*). The confidence interval width was recalculated every 50 iterations, and convergence was declared when the relative change in interval width between consecutive blocks was below 2% for three consecutive blocks simultaneously for all three parameters.

To compare parameter estimates between the control and cCpE-treated conditions, we calculated the empirical distribution of the differences between parameter estimates obtained under the two conditions. The 2.5^th^ and the 67.5^th^ percentiles of this distribution were used to obtain the 65% confidence interval (CI) for the difference using the percentile bootstrap method. A difference was considered statistically significant if the resulting 95% CI did not contain zero, corresponding to rejection of the null hypothesis of no difference between conditions at the α = 0.05 significance level.

### Mechanical stress inference

We used ForSys ^3,4^in its static modality to infer mechanical stress in the epicardium. At each triple junction (union of at least three cells) a pivot can be defined. Here it is assumed that there exist forces at the intersection of cell membranes of two different cells. The membranes are contracting and in consequence pulling the pivot in the direction tangential to the membrane interface. It is assumed that the force can be written as 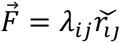 where *λ_ij_* is an unknown constant which represent the strength of the force between triple junctions *i* and *j* and 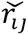 is a versor going from *i* to *j*. Each triple junction of the tissue is assumed to be in mechanical equilibrium. The force equilibrium at pivot *i* which is linked to pivots *j*, *k* and *l* is:

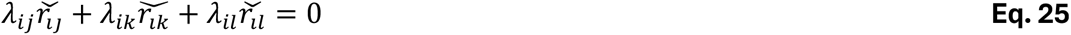

This defines two equilibrium equations per triple junction, one for the force components in the horizontal direction and one for the vertical one. To ensure that the trivial solution is avoided (where all forces are zero) one extra equation is added. The sum of all *λ_i_*_j_ is equal to the number vertices in the tissue. This essentially corresponds to asking that the mean value of *λ_i_*_j_ is equal to one. ForSys creates a matrix from a segmented image where each row is one of these equations. We used the non-negative least squares solver within ForSys to solve this matrix and estimate the values of the forces.

To evaluate differences in the mechanical stress and pressures of the tissue between the two experimental groups (analyzed using ForSys), we first evaluated their distribution, analyzing normality using Shapiro-Wilk test ^5^. Because data was not normal, non-parametric statistical tests were used. A two-sample two-tailed Kolmogorov Smirnov ^6^test was used to compare overall shape of the continuous distributions between the two groups, while a two-tailed Mann-Whitney U test ^7^ was conducted to compare median values. Statistical significance was set at *α* = 0.05.

### Transepithelial Electrical Resistance (TEER) measurements

Caco-2 cells were seeded at 2.68 × 10^5^ cells/cm^2^ on 12-well Corning Transwell® plates with 0.4 µm polycarbonate inserts (CLS3401, Corning) and cultured in high-glucose DMEM containing phenol red (D0816, Sigma-Aldrich) supplemented with 10% FBS and 1% penicillin-streptomycin. Cells were grown until stable TEER values were reached and subsequently treated with cCpE, mCherry-cCpE or mCherry-cCpE-Y306A/L315A diluted in cell culture medium. TEER was measured using a Millicell ERS 3.0 Voltohmmeter (MERS03000, Merck) every 2 h for 24 h. Measurement temperature was kept consistent throughout the experiment. Resistance values were normalized to the corresponding baseline (0 h) value for each well.

### Nanoindentation

Prior to measurement, sections were thawed and washed five times in 80% PBS (aPBS). A polydimethylsiloxane (PDMS; Sylgard 184, Dow Corning) chamber was placed on the slide to ensure hydration over the section. Mechanical properties were measured using the Chiaro nanoindenter (Optics11 Life, Amsterdam, The Netherlands). A spherical probe of radius 3.5 µm and spring constant 0.44N/m was used to indent the epicardium and another probe of radius 11 µm and spring constant 0.45N/m was used to indent the myocardium. The probe was calibrated before each session by indenting a glass dish filled with aPBS (80%) and was verified against the manufacturer’s factory calibration value.

Viscoelastic properties of the epicardium were assessed by dynamic measurement analysis (DMA), in which the probe was held at a 0.5 µm static offset and oscillated at 80 nm amplitude across frequencies of 1, 2, 4, 10, and 20 Hz (5 periods each) yielding storage and loss moduli, and loss tangent. At least 80 indentations were performed per section; one section was indented per animal and four animals per condition. Force-displacement curves without a clear contact point were discarded. For DMA as well, each force-displacement curve was inspected for contact quality, and those without a well-defined contact point were discarded ^8^.

The apparent Young’s modulus of the myocardium was measured by indenting 1 µm deep at 1 µm/s. Force-displacement curves were fitted with the Hertz contact model assuming a Poisson’s ratio of 0.5. Mean fit residuals were high (0.92 ± 0.08 SD for vehicle-treated, and 0.64 ± 0.05 SD for cCpE-treated). At least 130 readings were taken across the uninjured part of the myocardium. One section was indented per animal, three animals per condition. Atrioventricular canal was used as an anatomical landmark to ensure comparable regions of the ventricle were indented.

The apparent Young’s Modulus calculations and DMA analysis were done using Optics11 Prova software (v0.6.4).

### Image acquisition and analysis

Immunofluorescence images were acquired using a Leica Stellaris 5x confocal microscope. Zeiss AxioScan.Z1 Slide Scanner and/or Evident Slideview VS200 was used for high content immunofluorescence imaging and for collagen birefringence visualization using circularly polarized light. LAS X v1.4.4 (Leica Microsystems), OlyVIA v4.2 (Evident), Fiji (ImageJ) v1.54f, and Imaris v11.0.1 (Oxford Instruments) were used for image analysis.

### Single-nuclei Isolation, library preparation, and sequencing

Hearts were harvested from wild-type animals for unoperated and sham-operated conditions, and from tgTol2(*CAG:Nucbow*)^Sinon^ for vehicle- and cCpE-treated conditions. *Nucbow* reporter animals were injected with TAT-Cre recombinase prior to injury as described in Fig.2a. After harvesting, hearts were submerged in ice cold heparin-aPBS (1U/mL) and bisected into two hemispheres. The whole heart (including outflow tract and atria) was kept for post-metamorphic homeostasis samples whereas only the ventricle was kept for sham-operated and cryoinjured samples. The inside surface of each hemisphere was gently washed using a pasteur pipette to remove as much blood as possible without disrupting the tissue. Samples were then rinsed 3 times with ice-cold heparin-aPBS and 3 times with ice-cold aPBS. After removing excess buffer with tissue paper, each heart was frozen separately in liquid nitrogen. Nuclei isolation was performed using a commercial kit (10x Genomics, 1000493) following the manufacturer’s protocol with minor adjustments. Briefly, samples were thawed on ice for 15 min then transferred to the sample dissociation tube containing 50 µL of 1x lysis buffer. Samples were then homogenized on ice using surgical scissors for 1 min, followed by 1 min grinding with the provided pestle. Thereafter, 650 µL of lysis buffer was added and the whole solution was pipetted 10 times with a wide-bore pipette followed by 10 times with a narrow-bore pipette and allowed to incubate on ice for 2 min. Debris removal was then performed using the supplied Nuclei Isolation Column and 1x Debris Removal Buffer following the manufacturer’s protocol. All subsequent centrifugation and wash steps were also performed according to the manufacturer’s protocol. 1x Lysis Buffer and 1x Density Centrifugation Buffer were supplemented with 0.5 U/ µL of Protector RNAse inhibitor (Roche, 3335402001) whereas 1x Wash Buffer was supplemented with 1U/ µL Protector RNAse inhibitor. After two washes, the nuclei suspension was filtered using a 30 µm MACS SmartStrainer (Miltenyi Biotec, 130-068-458) and stained with 7-aminoactinomycin D (Miltenyi Biotec, 170-081-088) at a 1:10 dilution prior to fluorescence activated cell sorting (FACS). FACS was performed at the Biomedicum Flow Cytometry Core Facility (BFC) using a FACSAria Fusion Sorter (BD). Each sample was sorted for a maximum of 1 h and kept on ice until library preparation for single nuclei RNA sequencing. Library preparation was performed by the National Genomics Infrastructure (NGI) using the Chromium GEM-X Single Cell 3’ v4 Gene Expression kit (10x) with a target input of 20,000 nuclei per lane. Libraries were sequenced using a NovaSeq X Plus system (Illumina) with a target of 50,000-100,000 reads per nucleus.

### snRNA-seq alignment and quality control

Raw FASTQ files from snRNA-seq were mapped to the *P. waltl* NCBI genome (NCBI RefSeq: GCF_031143425.1) using *CellRanger* (v6.0.1). *CellBender* (v0.3.0) was used to remove ambient RNA. Nuclei with < 200 genes and > 50,000 transcripts were removed ^6^. Genes expressed in < 3 nuclei were filtered out. Doublet scores (ranging 0 to 1) were computed using the Python package *Scrublet*, and nuclei with doublet scores above the automatically determined threshold were discarded (Unoperated: 0.3, Sham-operated: 0.25, Vehicle-treated: 0.28, cCpE-treated: 0.27) ^10^. The remaining nuclei were analysed using *Seurat v5.3.0*. ^11^.

### snRNA-seq data analysis

Samples were merged, normalised and scaled, and the variables nCount (number of transcripts), nFeature (number of genes), S.Score (S-phase cell cycle score), and G2M.Score (G2/M-Phase cell cycle score) were regressed out. The closest *P. waltl* orthologues of known cell cycle genes were identified and used for regression ^12^. Samples were integrated using Harmony, and 25 dimensions were retained. Coarse-grain clustering using the Leiden algorithm was performed at resolution 0.6. Clusters were annotated based on canonical marker genes from literature and inferred localization on spatial transcriptomics sections. The epicardial gene module was quantified using the *AddModuleScore()* function in Seurat.

### Epicardium and putative EPDC subset analysis

Three clusters corresponding to epicardium and putative EPDCs were subsetted for further analysis. Cells belonging to these clusters were reanalysed, including normalization, scaling, variable regression, and Harmony-based integration. 15 dimensions were retained for downstream analysis, and Leiden clustering was performed at a resolution of 1.0, resulting in 16 clusters. Clusters sharing top marker genes were subsequently merged, yielding 13 final clusters. Top markers for the clusters were calculated using *FindAllMarkers()*

### Differential gene expression analysis in snRNA-seq

Differential gene expression analysis between conditions was performed using the MAST test implemented in the Seurat function *FindMarkers()*. Covariates defined were nFeature (number of genes) and nCount (number of transcripts). All reported differentially expressed genes have adjusted *P*-value < 0.05.

### Gene Ontology (GO) overrepresentation analysis in snRNA-seq and spatial transcriptomics

GO overrepresentation analysis was performed using *enrichGO* from the R package *clusterProfiler* (adjusted *P*-value < 0.05, q-value < 0.05, minimum gene set size = 3) ^13^.

### Integration of spatial transcriptomics and snRNAseq data

To infer the spatial localization of clusters identified by snRNA-seq, we used the anchor-based transfer learning method implemented in *Seurat v5.3.1* ^14^. CCA was used to identify transfer anchors between the log-normalized snRNA-seq dataset (reference dataset) and the spatial transcriptomics dataset (query dataset). This approach assigns a prediction score ranging from 0 to 1 for each snRNAseq cluster to every spatial transcriptomics spot. Prediction scores for snRNA-seq clusters (epicardial lineage clusters 01-13 and main dataset clusters 01-27) were grouped and averaged for the major cardiac cell types resulting in prediction scores for cardiomyocytes, erythrocytes, myeloid cells, lymphocytes, epicardium, smooth muscle cells, valve interstitial cells, endothelial cells and fibroblasts, and glial cells. Prediction scores > 0.01 were visualized on representative spatial transcriptomic sections.

### Regenerative gene expression profile

Gene expression was aggregated, log-normalized, and scaled for each cluster-condition combination using the *AggregateExpression()* function in Seurat after excluding cluster-condition combinations with < 3 cells. The putative regenerative profile was defined as the average expression of genes *Tnc*, *Chst15*, *Tgm2*, *Oxtr*, *Tgfbi*, *Cilp*, *Loc1382C175S*, *Slc2a1*, *Slc1a5*, *Cysltr1*, and *Cpm* across cluster-condition groups corresponding to Clusters 07-06 in unoperated, sham-operated, and vehicle-treated conditions. Spearman’s correlation was calculated between the putative regenerative expression profile and all genes in the dataset. Genes with correlation coefficients > 0.6 were designated members of the putative regenerative gene module. Module enrichment for vehicle-treated versus cCpE-treated conditions was computed for each cluster of interest, wherein all genes were ranked by the log2 fold changes between vehicle- and cCpE-treated conditions, and the enrichment of the putative regenerative module was computed using a competitive gene test implemented in *cameraPR()* function from the *limma* package ^15^. Gene Ontology (GO) overrepresentation analysis was performed, and selected regenerative module genes and GO terms were visualized using a network made in Cytoscape (v3.10.4).

### Spatial transcriptomics library preparation and sequencing

Spatial transcriptomic profiling was performed using Visium v1 3’ Gene Expression technology (10x Genomics). Fresh-frozen tissues were cryosectioned at 10 µm and positioned onto Visium slides (PN-1000184, 10x Genomics), followed by methanol fixation and hematoxylin and eosin (HGE) staining according to manufacturer’s instructions (CG000239 Rev A). Imaging was performed using a Zeiss AxioScan.Z1 slide scanner. The optimal tissue permeabilization time was determined using the Visium Spatial Tissue Optimization Kit (PN-1000163, 10x Genomics) and set to 6 min. Libraries were prepared according to the manufacturer’s instructions and sequenced on an Illumina NovaSeq X using one lane of a 10B-3000 flow cell with a custom read configuration of 28-10-10-60cycles.

### Spatial transcriptomics alignment, quality control, and clustering

Raw FASTQ files from spatial transcriptomics were mapped to the *P. waltl* NCBI genome (NCBI RefSeq: GCF_031143425.1) using *SpaceRanger* (v2.0.1). Spots with < 200 detected genes or > 20,000 transcripts were excluded. Genes detected in ≥ 3 spots per section and in ≥ 6 distinct sections were retained for downstream analysis. After filtering, the top 2,000 variable genes were used to perform principal component analysis (PCA). Animal-specific effects were corrected using canonical correlation analysis (CCA), and 35 dimensions were retained. Leiden clustering at a resolution of 1.6 identified 22 clusters, which were annotated based on their spatial localization and expression of cluster-specific marker genes.

### Differential gene expression analysis

Differential gene expression (DGE) analysis was performed using the MAST test implemented in the Seurat function *FindMarkers()*. Animal, nFeature (number of detected genes), and nCount (number of detected transcripts) were included as covariates in the model. Despite rigorous standardization of section selection for spatial transcriptomic analysis, variation in the plane of sectioning resulted in differences in the size of the captured injury area, particularly among vehicle-treated hearts. As injury size may reflect differences in regenerative progression and consequently influence gene expression profiles, cluster-wise DGE analyses were performed comparing (1) vehicle-treated and cCPE-treated hearts and (2) sections from vehicle-treated hearts with small versus large injury areas, representing more advanced and ongoing regeneration, respectively. Genes that were significantly differentially expressed in the vehicle-treated versus cCpE-treated comparison, but not in the small versus large injury area comparison, were retained as candidate treatment-specific transcriptional changes independent of regenerative stage. Significantly overrepresented GO terms were grouped into broader biological categories by manual curation.

### Identifying *P. waltl* genes with gene descriptions in addition to gene symbols

The NCBI-provided gene symbols for the *P. waltl* genome reference (GCF_031143425.1, downloaded 2022-11-26) are used unless otherwise stated. Genes of uncertain function or undetermined orthology are annotated by NCBI with their locus number (e.g. Loc138304364), but a description may be provided (e.g. Cldn6-like, https://www.ncbi.nlm.nih.gov/books/NBK3840/). To aid identification of canonical cell type markers, cell cycle genes, and Claudin genes, these NCBI-provided gene descriptions were used in addition to gene symbols. Gene descriptions were mapped to putative gene symbols as follows: first, the GTF file was used to extract information on gene ID, description, and gene biotype. Protein-coding genes were processed in two categories: those with existing gene symbols, and others annotated as a locus, i.e. starting with “LOC”. Existing gene symbols were converted according to the human gene symbol convention and used where available. For genes starting with “LOC”, human or mouse gene symbols were retrieved from UniProt using the gene description. Among the genes missing a human symbol but successfully annotated with a mouse symbol, Ensembl gene IDs and the getHomologs() function were used to map gene symbols from mouse to human. The remaining genes were annotated if their human gene symbols could be identified through manual curation based on the gene description. Duplicate human gene symbols were made unique by appending their “LOC” identifiers or “PW”.

### Quantification of apoptotic epicardial cells

Apoptotic epicardial cells were quantified by enumerating the TUNEL^+^ Pan-CK^+^ cells and normalizing this value to the total number of Pan-CK^+^ epicardial cells within the analyzed region. Analysis was limited to the ventricle. 4-5 sections were analyzed per animal.

### Quantification of proliferating epicardial cells

The number of pHH3^+^ Pan-CK^+^ mitotic epicardial cells was enumerated and divided by the total number of epicardial cells. Analysis was limited to the ventricle. 4-5 sections were analyzed per animal. The proportion of apoptotic/proliferating epicardial cells was expressed as a percentage.

### Quantification of EPDCs in the injury area

The number of TAT-Cre labelled EPDCs within the injury area was enumerated and normalized to the injury area of each section. Injury area was measured using the measurement tool in Leica LAS X software. 3-4 sections were analyzed per animal.

### Quantification of apoptotic EPDCs in the injury area

The number of apoptotic (TUNEL^+^) TAT-Cre labelled EPDCs was enumerated and divided by the total number of EPDCs at the injury site, then expressed as a percentage.

### Quantification of proliferating EPDCs in the injury area

The number of proliferating (pHH3^+^) TAT-Cre labelled EPDCs were enumerated and divided by the total number of EPDCs at the injury site, then expressed as a percentage.

### Quantification of epicardial nuclei density

Number of pan-CK^+^ cell nuclei was enumerated on 14 μm cryosections. For each animal three sections were analysed. Two rectangular regions of interest of fixed area (0.01366 mm²) were placed over the pan-CK⁺ epicardial layer in each section: one adjacent to the atrioventricular canal and one at the injury border. Regions were positioned so that the epicardial layer ran along the long axis of the rectangle. Values from both regions and all sections were averaged to give a single value per animal.

### Quantification of nuclear p-ERK intensity

To assess nuclear p-ERK levels, DAPI^+^ nuclei in the pan-CK^+^ epicardial layer were manually selected as regions of interest (ROI) in Fiji. Then, these ROIs were applied on the corresponding p-ERK immunofluorescence channel and the mean fluorescence intensity within each ROI was measured.

### Clonality Analysis

To assess clonal relationships between epicardial cells, the images were acquired as separate single channel grey-scale images for each fluorophore mCherry, mEYFP and mCerulean and rendered in red, green and blue for visualization. TAT-Cre labelled nuclei in the Pan-CK^+^ epicardial layer were manually selected as regions of interest (ROIs) in Fiji. Using the Multi Measure tool, centroid coordinates and mean gray values were extracted for each nucleus from all three channels. The exported data were subsequently processed in MATLAB (MathWorks).

For each animal, the minimum fluorescence intensity in each channel was identified and used as the background value. The relative channel weight was calculated for each nucleus by normalizing the background-corrected intensity of each channel to the sum of the intensities across all three channels ^16^.

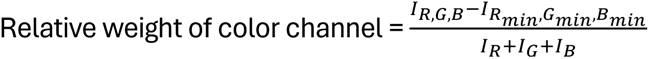

The centroid coordinates of each nucleus were used to organize nuclei according to their spatial positions, ensuring that physically neighboring nuclei were compared during color-distance calculations. Color distance and color spread coefficient values were then calculated using the formulas below to assess clonal relationships between neighboring nuclei ^17^.

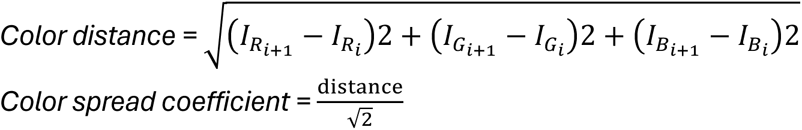

To determine the color spread coefficient threshold, values ranging from 0.02 to 0.08 were evaluated in the sham-operated dataset, where minimal clonal expansion is expected (Extended Data Fig.1d). A marked increase in doublet detection was observed between 0.05 and 0.06, suggesting that higher thresholds may increase the incidence of false-positive classifications. Consequently, neighboring nuclei with a color spread coefficient below 0.05 was considered clonally related. Three sections were imaged per animal, and 4-6 animals were analyzed for each condition.

### Imaging of whole-mount samples

For imaging the apical cell membrane of the epicardial cells, whole-mount sample was mounted with the cut surface facing up, away from the objective, and z-stacks were acquired starting at the apical surface with a 40x glycerol objective at 0.75x zoom, using 2 µm steps (10-20 steps per stack depending on sample). For imaging the TNC-rich scaffold, whole-mount sample was mounted with the cut surface down, facing the 10x air objective. Z-stacks were acquired at 1x objective zoom in 10 µm steps (23-26 sections per stack).

### Epicardial cell segmentation

Max projected images of the Z-stack for the ZO-1or E-cadherin were used for cell segmentation. Images were segmented in Ilastik (Version 1.4.1) using the Pixel Classification and Multicut workflows and exported as a label image. Segmented images were manually inspected and corrected as needed.

### Quantification of cell shape and orientation

From the segmented images described above, apical cell area, aspect ratio and orientation were quantified with a custom Python script using the io and measure modules from the scikit-image together with NumPy ^18,16^. Cell area and aspect ratio were obtained from region properties, with areas reported in μm ² using the pixel size of the corresponding acquisition and aspect ratio defined as the ratio of major to minor axis of the equivalent ellipse. Orientation was calculated using the two-fold (nematic, p=2) shape function introduced for epithelial cells by ^20^evaluated over the cell outline, extracted by the marching-squares algorithm:

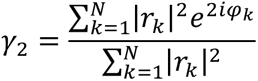

where r_k_ and *φ_k_* denote the position and polar angle of the k-th outline point relative to the cell centroid. The orientation of each cell is given by:

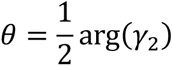

reported in degrees in the range (–90°, 90°).

For each field-of-view (FOV), the alignment parameter R was computed as the resultant length of the doubled angles,

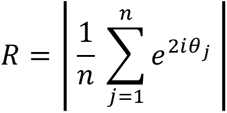

where n is the number of cells in that FOV. R=0 indicates that orientations are spread evenly and R=1 indicated perfect alignment. To test whether orientations within a FOV departed from a uniform distribution, a Rayleigh test was used with the p value obtained from the large-sample approximation.

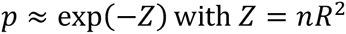

A FOV was classified as significantly aligned when p<0.05. The test statistic was the difference in the percentage of significantly aligned FOVs between conditions, calculated across all FOVs. Because multiple FOVs were imaged per animal, FOVs are not independent observations; permutations were therefore performed at the level of animals, with all FOVs from an animal reassigned together. For each pairwise comparison, all possible reassignments of animals into two groups were enumerated, giving exact p values. Pairwise p values were Holm-corrected and compared against α = 0.05. Statistical tests were performed in Python using Numpy, pandas and SciPy ^21^.

### Quantification of CLDN6 intensity in whole-mount images

CLDN6 intensity at cell-cell junctions was quantified using a custom Python script applied to CLDN6 IF images and corresponding segmented images described above. The script uses scikit-image, SciPy, Numpy and pandas as above, with tifffile for image reading. The analysis was restricted to images from the same staining and imaging session, acquired with identical settings. A 1-pixel (px) skeleton was generated from the segmented image to provide a topological representation of cell-cell interfaces. Skeleton pixels with three or more neighbours were classified as vertices and removed so that each remaining connected segment corresponded to a single bicellular junction. A measurement band was defined by expanding 1-px on either side of the skeleton, resulting in a 3-px-wide region spanning the length of each junction, and mean CLDN6 fluorescence intensity was calculated within this region for each cell-cell junction. A background value measured for each image as the mean of five square ROIs placed in junction-free regions, was subtracted from every junction mean. For each cell, junctional CLDN6 (or ZO1) intensity was expressed as the mean of its bordering junctions weighted by junction length. Heterogeneity of CLDN6 between the junctions of a single cell was expressed as the coefficient of variation, calculated on raw mean junctional intensities.

### Quantification of ZO-1 intensity in whole-mount images

Junctional ZO-1 intensity was quantified using the same python script described above for CLDN6.

### Border muscle protrusions

The length and number of cardiomyocyte protrusions extending into the TNC-rich extracellular matrix scaffold were quantified using the measurement tool in Leica Las X software. For each whole-mount sample, ten z-stack images spanning the injury site were acquired from the center of the injury at 10 µm intervals and used to generate a maximum intensity projection. Only protrusions that were fully contained within the acquired imaging volume were included in the analysis.

### TNC intensity quantification

Whole-mount images of the ventricular apex were cropped to isolate the TNC-rich scaffold region and imported into Imaris (11.0.1, Oxford Instruments). Scaffold surfaces were rendered using the surface module, and mean TNC fluorescence intensity was quantified for each surface. Extracted intensity values were compiled for downstream analysis.

### Quantification of TNC-rich bridge orientation

Quantification of TNC-rich bridge orientation was performed in a blinded manner using Imaris (11.0.1, Oxford Instruments). Whole-mount images were imported and divided into three regions spanning the curvature of the injury. Within each region, five spots were placed along the TNC signal at the injury apex to define a local reference plane that accounted for tissue curvature. In addition, two spots were placed at the start and end points of each TNC-rich bridge connecting the injury apex to the base. The xyz coordinates of all spots were exported and compiled for analysis. Bridge angles relative to the corresponding local reference plane were subsequently calculated using a custom script adapted from ^22^.

To evaluate TNC bridge orientation, a second custom script also adapted from ^22^ was used. The calculated angles and subsequently their cumulative distribution were represented as empirical cumulative distribution functions (ECDFs). A parameterised model of 3D orientation was then fitted separately to each treatment group by optimising the model parameters, D, to minimize the squared differences between the modelled and experimental ECDFs. Fitted model distributions were generated from optimised parameters, and 65% confidence intervals were estimated by bootstrap resampling. The observed angle distributions were additionally compared with the theoretical isotropic orientation distribution (F(alpha)=sin(alpha)), using a two-sided Kolmogorov–Smirnov (KS) test.

### Sarcomere organization assessment

Sarcomere organization was assessed semi-quantitatively in tissue sections by determining the presence of organized sarcomeres within a distance of 30 µm from the tip of the muscle protrusion. Protrusions were assigned a binary score, with 1 or 0 indicating the presence or absence of organized sarcomeres, respectively.

## Notes

https://github.com/delfiperez/Claudin-dependent_clonal_expansion_couples_epicardial_activation_to_regenerative_fate_acquisition

https://github.com/Elif-Eroglu-Lab/cCpE_paper_2026.git

https://doi.org/10.5281/zenodo.23068479

https://github.com/borgesaugusto/forsys

https://doi.org/10.5281/zenodo.11282554

### Competing Interest Statement

The authors have declared no competing interest.

## References

1. Lepilina, A. et al. A dynamic epicardial injury response supports progenitor cell activity during zebrafish heart regeneration. Cell 127, 607–616 (2006).

2. Wang, J., Karra, R., Dickson, A. L. & Poss, K. D. Fibronectin is deposited by injury-activated epicardial cells and is necessary for zebrafish heart regeneration. Dev. Biol. 382, 427–435 (2013).

3. Mercer, S. E., Odelberg, S. J. & Simon, H.-G. A dynamic spatiotemporal extracellular matrix facilitates epicardial-mediated vertebrate heart regeneration. Dev. Biol. 382, 457–466 (2013).

4. Marro, J., Pfefferli, C., de Preux Charles, A.-S., Bise, T. & Jaźwińska, A. Collagen XII Contributes to Epicardial and Connective Tissues in the Zebrafish Heart during Ontogenesis and Regeneration. PLoS ONE 11, e0165467 (2016).

5. González-Rosa, J. M., Peralta, M. & Mercader, N. Pan-epicardial lineage tracing reveals that epicardium derived cells give rise to myofibroblasts and perivascular cells during zebrafish heart regeneration. Dev. Biol. 370, 173–186 (2012).

6. Xia, Y. et al. Activation of a transient progenitor state in the epicardium is required for zebrafish heart regeneration. Nat. Commun. 13, 7704 (2022).

7. Sun, J. et al. hapln1 Defines an Epicardial Cell Subpopulation Required for Cardiomyocyte Expansion During Heart Morphogenesis and Regeneration. Circulation 146, 48–63 (2022).

8. Eroglu, E. et al. Epicardium-derived cells organize through tight junctions to replenish cardiac muscle in salamanders. Nat. Cell Biol. 24, 645–658 (2022).

9. Zhou, B. et al. Adult mouse epicardium modulates myocardial injury by secreting paracrine factors. J. Clin. Invest. 121, 1864–1604 (2011).

10. Quijada, P., Trembley, M. A. & Small, E. M. The role of the epicardium during heart development and repair. Circ. Res. 126, 377–364 (2020).

11. Beutel, O., Maraspini, R., Pombo-García, K., Martin-Lemaitre, C. & Honigmann, A. Phase separation of zonula occludens proteins drives formation of tight junctions. Cell 179, 623–636.e11 (2016).

12. Schwayer, C. et al. Mechanosensation of Tight Junctions Depends on ZO-1 Phase Separation and Flow. Cell 179, 637–652.e18 (2016).

13. Citi, S. et al. A short guide to the tight junction. J. Cell Sci. 137, (2024).

14. Günzel, D., Lehmann, M. & Honigmann, A. Tight junction structure, assembly and (dys)function. Nat. Rev. Mol. Cell Biol. (2026) doi:10.1038/s41580-026-00678-w.

15. Mineta, K. et al. Predicted expansion of the claudin multigene family. FEBS Lett. 585, 606–612 (2011).

16. Günzel, D. & Yu, A. S. L. Claudins and the modulation of tight junction permeability. Physiol. Rev. 93, 525–566 (2013).

17. Kashihara, H. et al. Functional landscape of mechanistic diversity in 27 claudin family members at tight junctions. Sci. Adv. 11, eadx7431 (2025).

18. Shen, L., Weber, C. R. & Turner, J. R. The tight junction protein complex undergoes rapid and continuous molecular remodeling at steady state. J. Cell Biol. 181, 683–665 (2008).

19. Higashi, T., Arnold, T. R., Stephenson, R. E., Dinshaw, K. M. & Miller, A. L. Maintenance of the Epithelial Barrier and Remodeling of Cell-Cell Junctions during Cytokinesis. Curr. Biol. 26, 1826–1842 (2016).

20. Stephenson, R. E. et al. Rho flares repair local tight junction leaks. Dev. Cell 48, 445–456.e5 (2016).

21. Varadarajan, S., Stephenson, R. E. & Miller, A. L. Multiscale dynamics of tight junction remodeling. J. Cell Sci. 132, (2016).

22. Vermij, S. H., Abriel, H. & van Veen, T. A. B. Refining the molecular organization of the cardiac intercalated disc. Cardiovasc. Res. 113, 256–275 (2017).

23. Zhang, J. et al. Cardiomyocyte Expression of ZO-1 Is Essential for Normal Atrioventricular Conduction but Does Not Alter Ventricular Function. Circ. Res. 127, 284–267 (2020).

24. Horowitz, A., Chanez-Paredes, S. D., Haest, X. & Turner, J. R. Paracellular permeability and tight junction regulation in gut health and disease. Nat. Rev. Gastroenterol. Hepatol. 20, 417–432 (2023).

25. Meoli, L. & Günzel, D. The role of claudins in homeostasis. Nat. Rev. Nephrol. 19, 587–603 (2023).

26. Yen, C. Y. T., Kagioglou, S. & Eroglu, E. European Research Council-funded grant: tight junctions and heart regeneration. Eur. Heart J. 46, 2841–2843 (2025).

27. Hanna, P. C., Wieckowski, E. U., Mietzner, T. A. & McClane, B. A. Mapping of functional regions of Clostridium perfringens type A enterotoxin. Infect. Immun. 60, 2110–2114 (1662).

28. Sonoda, N. et al. Clostridium perfringens enterotoxin fragment removes specific claudins from tight junction strands: Evidence for direct involvement of claudins in tight junction barrier. J. Cell Biol. 147, 165–204 (1666).

29. Ogbu, C. P., Roy, S. & Vecchio, A. J. Disruption of Claudin-Made Tight Junction Barriers by Clostridium perfringens Enterotoxin: Insights from Structural Biology. Cells 11, (2022).

30. Loulier, K. et al. Multiplex cell and lineage tracking with combinatorial labels. Neuron 81, 505–520 (2014).

31. Joven, A. et al. Cellular basis of brain maturation and acquisition of complex behaviors in salamanders. Development 145, (2018).

32. Brockway, N. L. et al. Multicolor lineage tracing using in vivo time-lapse imaging reveals coordinated death of clonally related cells in the developing vertebrate brain. Dev. Biol. 453, 130–140 (2016).

33. Choi, W. et al. Remodeling the zonula adherens in response to tension and the role of afadin in this response. J. Cell Biol. 213, 243–260 (2016).

34. Wada, M., Tamura, A., Takahashi, N. & Tsukita, S. Loss of claudins 2 and 15 from mice causes defects in paracellular Na+ flow and nutrient transport in gut and leads to death from malnutrition. Gastroenterology 144, 366–380 (2013).

35. Waldow, A. et al. cCPE Fusion Proteins as Molecular Probes to Detect Claudins and Tight Junction Dysregulation in Gastrointestinal Cell Lines, Tissue Explants and Patient-Derived Organoids. Pharmaceutics 15, (2023).

36. Veshnyakova, A. et al. On the interaction of Clostridium perfringens enterotoxin with claudins. Toxins (Basel) 2, 1336–1356 (2010).

37. Marinari, E. et al. Live-cell delamination counterbalances epithelial growth to limit tissue overcrowding. Nature 484, 542–545 (2012).

38. Borges, A. et al. ForSys: Non-invasive stress inference from time-lapse microscopy. iScience 28, 113685 (2025).

39. Borges, A. et al. Protocol for inferring mechanical stresses in tissues using ForSys, an open-source Python tool. STAR Protocols 7, 104565 (2026).

40. Weinberger, M., Simões, F. C., Patient, R., Sauka-Spengler, T. & Riley, P. R. Functional Heterogeneity within the Developing Zebrafish Epicardium. Dev. Cell 52, 574–560.e6 (2020).

41. Knight-Schrijver, V. R. et al. A single-cell comparison of adult and fetal human epicardium defines the age-associated changes in epicardial activity. Nat. Cardiovasc. Res. 1, 1215–1226 (2022).

42. Meier, A. B. et al. Epicardioid single-cell genomics uncovers principles of human epicardium biology in heart development and disease. Nat. Biotechnol. 41, 1787–1800 (2023).

43. Runne, C. & Chen, S. PLEKHG2 promotes heterotrimeric & protein βγ-stimulated lymphocyte migration via Rac and Cdc42 activation and actin polymerization. Mol. Cell. Biol. 33, 4264–4307 (2013).

44. Kremerskothen, J., Plaas, C., Kindler, S., Frotscher, M. & Barnekow, A. Synaptopodin, a molecule involved in the formation of the dendritic spine apparatus, is a dual actin/alpha-actinin binding protein. J. Neurochem. 92, 567–606 (2005).

45. Liu, L. et al. Hairless is a histone H3K6 demethylase. FASEB J. 28, 1534–1542 (2014).

46. Liu, L. et al. Hairless regulates heterochromatin maintenance and muscle stem cell function as a histone demethylase antagonist. Proc Natl Acad Sci USA 118, (2021).

47. Kroes, R. A. et al. Overexpression of ST6GalNAcV, a ganglioside-specific alpha2,6-sialyltransferase, inhibits glioma growth in vivo. Proc Natl Acad Sci USA 107, 12646–12651 (2010).

48. Li, J. et al. Site-Specific N-Glycoproteomic Analysis Reveals Upregulated Sialylation and Core Fucosylation during Transient Regeneration Loss in Neonatal Mouse Hearts. J. Proteome Res. 19, 3161–3200 (2020).

49. Trepat, X. et al. Universal physical responses to stretch in the living cell. Nature 447, 562–565 (2007).

50. Streef, T. J. & Smits, A. M. Epicardial contribution to the developing and injured heart: exploring the cellular composition of the epicardium. Front. Cardiovasc. Med. 8, 750243 (2021).

51. Ma, X., Sung, D. C., Yang, Y., Wakabayashi, Y. & Adelstein, R. S. Nonmuscle myosin IIB regulates epicardial integrity and epicardium-derived mesenchymal cell maturation. J. Cell Sci. 130, 2666–2706 (2017).

52. Sánchez, N. S. & Barnett, J. V. TGFβ and BMP-2 regulate epicardial cell invasion via TGFβR3 activation of the Par6/Smurf1/RhoA pathway. Cell. Signal. 24, 536–548 (2012).

53. Chen, X., Zhong, X. & Huang, G. N. Heart regeneration from the whole-organism perspective to single-cell resolution. *npj Regen*. Med. 9, 34 (2024).

54. Jopling, C. et al. Zebrafish heart regeneration occurs by cardiomyocyte dedifferentiation and proliferation. Nature 464, 606–606 (2010).

55. Boskovic, S. et al. The stress responsive gene ankrd1a is dynamically regulated during skeletal muscle development and upregulated following cardiac injury in border zone cardiomyocytes in adult zebrafish. Gene 792, 145725 (2021).

56. van der Ven, P. F. M. et al. Characterization of muscle filamin isoforms suggests a possible role of γ-filamin/ABP-L in sarcomeric Z-disc formation. Cell Motility and the Cytoskeleton (2000).

57. Jung-Ching Lin, J., et al. Structure, Expression, and Function of a Novel Intercalated Disc Protein, Xin. J. Med. Sci. 25, 215–222 (2005).

58. Salmikangas, P. et al. Myotilin, a novel sarcomeric protein with two Ig-like domains, is encoded by a candidate gene for limb-girdle muscular dystrophy. Hum. Mol. Genet. 8, 1326–1336 (1666).

59. Xia, H., Winokur, S. T., Kuo, W. L., Altherr, M. R. & Bredt, D. S. Actinin-associated LIM protein: identification of a domain interaction between PDZ and spectrin-like repeat motifs. J. Cell Biol. 139, 507–515 (1667).

60. Granger, B. L. & Lazarides, E. Desmin and vimentin coexist at the periphery of the myofibril Z disc. Cell 18, 1053–1063 (1676).

61. Watanabe, T. et al. Localization of type IV collagen alpha chain in the myocardium of dilated and hypertrophic cardiomyopathy. Jpn Heart J 39, 753–762 (1668).

62. Beisaw, A. et al. AP-1 Contributes to Chromatin Accessibility to Promote Sarcomere Disassembly and Cardiomyocyte Protrusion During Zebrafish Heart Regeneration. Circ. Res. 126, 1760–1778 (2020).

63. Constanty, F. et al. Border-zone cardiomyocytes and macrophages regulate extracellular matrix remodeling to promote cardiomyocyte protrusion during cardiac regeneration. Nat. Commun. 16, 3823 (2025).

64. Wu, M. et al. Epicardial spindle orientation controls cell entry into the myocardium. Dev. Cell 19, 114–125 (2010).

65. Eisenhoffer, G. T. et al. Crowding induces live cell extrusion to maintain homeostatic cell numbers in epithelia. Nature 484, 546–546 (2012).

66. Matamoro-Vidal, A. & Levayer, R. Multiple influences of mechanical forces on cell competition. Curr. Biol. 29, R762–R774 (2016).

67. LeGoff, L. & Lecuit, T. Mechanical forces and growth in animal tissues. Cold Spring Harb. Perspect. Biol. 8, a016232 (2015).

68. Miroshnikova, Y. A. et al. Adhesion forces and cortical tension couple cell proliferation and differentiation to drive epidermal stratification. Nat. Cell Biol. 20, 66–80 (2018).

69. Gurdon, J. B. A community effect in animal development. Nature 336, 772–774 (1688).

70. Itoh, M. et al. Direct binding of three tight junction-associated MAGUKs, ZO-1, ZO-2, and ZO-3, with the COOH termini of claudins. J. Cell Biol. 147, 1351–1363 (1666).

71. Zihni, C., Mills, C., Matter, K. & Balda, M. S. Tight junctions: from simple barriers to multifunctional molecular gates. Nat. Rev. Mol. Cell Biol. 17, 564–580 (2016).

72. Wengerhoff, S. M., Weiss, A. R., Dwyer, K. L. & Dettman, R. W. A migratory role for EphrinB ligands in avian epicardial mesothelial cells. Dev. Dyn. 239, 568–606 (2010).

73. Xiao, Y. et al. Hippo Signaling Plays an Essential Role in Cell State Transitions during Cardiac Fibroblast Development. Dev. Cell 45, 153–166.e6 (2018).

74. Singh, A. et al. Hippo signaling mediators yap and taz are required in the epicardium for coronary vasculature development. Cell Rep. 15, 1384–1363 (2016).

75. Ramjee, V. et al. Epicardial YAP/TAZ orchestrate an immunosuppressive response following myocardial infarction. J. Clin. Invest. 127, 866–611 (2017).

76. Deng, S.-B. et al. Triiodothyronine promotes the proliferation of epicardial progenitor cells through the MAPK/ERK pathway. Biochem. Biophys. Res. Commun. 486, 372–377 (2017).

77. Gamba, L., Amin-Javaheri, A., Kim, J., Warburton, D. & Lien, C.-L. Collagenolytic Activity Is Associated with Scar Resolution in Zebrafish Hearts after Cryoinjury. J. Cardiovasc. Dev. Dis. 4, (2017).

78. Zlatanova, I. et al. An injury-responsive mmp14b enhancer is required for heart regeneration. Sci. Adv. 9, eadh5313 (2023).

79. Sun, J., Peterson, E. A., Chen, X. & Wang, J. hapln1a+ cells guide coronary growth during heart morphogenesis and regeneration. Nat. Commun. 14, 3505 (2023).

80. Cao, J. & Poss, K. D. The epicardium as a hub for heart regeneration. Nat. Rev. Cardiol. 15, 631–647 (2018).

81. Cheung, A. K. L. et al. Cysteine-rich intestinal protein 2 (CRIP2) acts as a repressor of NF-kappaB-mediated proangiogenic cytokine transcription to suppress tumorigenesis and angiogenesis. Proc Natl Acad Sci USA 108, 8360–8365 (2011).

82. Kedage, V., Strittmatter, B. G., Dausinas, P. B. & Hollenhorst, P. C. Phosphorylation of the oncogenic transcription factor ERG in prostate cells dissociates polycomb repressive complex 2, allowing target gene activation. J. Biol. Chem. 292, 17225–17235 (2017).

83. Zhang, S., et al. SRGN-mediated reactivation of the YAP/CRISPLD2 axis promotes aggressiveness of hepatocellular carcinoma. Int. J. Biol. Sci. 21, 3262–3285 (2025).

84. Wang, W. et al. PTPN14 is required for the density-dependent control of YAP1. Genes Dev. 26, 1656–1671 (2012).

85. Liu, X. et al. PTPN14 interacts with and negatively regulates the oncogenic function of YAP. Oncogene 32, 1266–1273 (2013).

86. Raha, S. et al. Molecular mechanism of augmentation of oncogenic activity of MYC by TAF2 in HCC. J. Biol. Chem. 113581 (2026) doi:10.1016/j.jbc.2026.113581.

87. Chen, Y. et al. PTGS2/GRP78 activation triggers endoplasmic reticulum stress leading to lipid metabolism disruption and cell apoptosis, exacerbating damage in bovine mastitis. Biomolecules 14, (2024).

## References

88. Joven, A. et al. Cellular basis of brain maturation and acquisition of complex behaviors in salamanders. Development 145, (2018).

89. Virtanen, P. et al. SciPy 1.0: fundamental algorithms for scientific computing in Python. Nat. Methods 17, 261–272 (2020).

90. Borges, A. et al. ForSys: Non-invasive stress inference from time-lapse microscopy. iScience 28, 113685 (2025).

91. Borges, A. et al. Protocol for inferring mechanical stresses in tissues using ForSys, an open-source Python tool. STAR Protocols 7, 104565 (2026).

92. Shapiro, S. S. & Wilk, M. B. An analysis of variance test for normality (complete samples). Biometrika 52, 561–611. (1665).

93. Massey, F. J. The Kolmogorov-Smirnov Test for Goodness of Fit. J. Am. Stat. Assoc. 46, 68–78 (1651).

94. Mann, H. B. & Whitney, D. R. On a test of whether one of two random variables is stochastically larger than the other. Ann. Math. Statist. 18, 50–60 (1647).

95. Ciccone, G. et al. Experimental and data analysis workflow for soft matter nanoindentation. JOVE (2022) doi:10.3761/63401-v.

96. Fleming, S. J. et al. Unsupervised removal of systematic background noise from droplet-based single-cell experiments using CellBender. Nat. Methods 20, 1323–1335 (2023).

97. Wolock, S. L., Lopez, R. & Klein, A. M. Scrublet: Computational Identification of Cell Doublets in Single-Cell Transcriptomic Data. Cell Syst. 8, 281–261.e6 (2016).

98. Hao, Y. et al. Dictionary learning for integrative, multimodal and scalable single-cell analysis. Nat. Biotechnol. 42, 263–304 (2024).

99. Tirosh, I. et al. Dissecting the multicellular ecosystem of metastatic melanoma by single-cell RNA-seq. Science 352, 186–166 (2016).

100. Yu, G., Wang, L.-G., Han, Y. & He, Q.-Y. clusterProfiler: an R package for comparing biological themes among gene clusters. OMICS 16, 284–287 (2012).

101. Stuart, T. et al. Comprehensive Integration of Single-Cell Data. Cell 177, 1888–1602.e21 (2016).

102. Wu, D. & Smyth, G. K. Camera: a competitive gene set test accounting for inter-gene correlation. Nucleic Acids Res. 40, e133 (2012).

103. Loulier, K. et al. Multiplex cell and lineage tracking with combinatorial labels. Neuron 81, 505–520 (2014).

104. Brockway, N. L. et al. Multicolor lineage tracing using in vivo time-lapse imaging reveals coordinated death of clonally related cells in the developing vertebrate brain. Dev. Biol. 453, 130–140 (2016).

105. van der Walt, S. et al. scikit-image: image processing in Python. PeerJ 2, e453 (2014).

106. Harris, C. R. et al. Array programming with NumPy. Nature 585, 357–362 (2020).

107. Armengol-Collado, J.-M., Carenza, L. N., Eckert, J., Krommydas, D. & Giomi, L. Epithelia are multiscale active liquid crystals. Nat. Phys. (2023) doi:10.1038/s41567-023-02176-0.

108. McKinney, W. Data structures for statistical computing in python. in Proceedings of the Sth Python in Science Conference 56–61 (SciPy, 2010). doi:10.25080/Majora-62bf1622-00a.

109. Jüschke, C., Xie, Y., Postiglione, M. P. & Knoblich, J. A. Analysis and modeling of mitotic spindle orientations in three dimensions. Proc Natl Acad Sci USA 111, 1014–1016 (2014).

